# A novel cross talk of AtRAV1, an ethylene responsive transcription factor with MAP kinases imparts broad spectrum disease resistance in plants

**DOI:** 10.1101/2020.01.08.899690

**Authors:** Ravindra Kumar Chandan, Rahul Kumar, Durga Madhab Swain, Srayan Ghosh, Prakash Kumar Bhagat, Sunita Patel, Ganesh Bagler, Alok Krishna Sinha, Gopaljee Jha

## Abstract

Plant diseases pose a serious threat to sustainable agriculture as controlling them in eco-friendly manner remains a challenge. In this study, we establish RAV1 as a master transcriptional regulator of defense genes in model plant Arabidopsis. The overexpression of *AtRAV1* provided disease resistance against necrotrophic fungal pathogen (*Rhizoctonia solani)* infection in *A. thaliana*. The transgenic lines exhibited enhanced expression of several defense genes including mitogen associated protein kinases (MAPKs) and the amplitude of their expression was further enhanced upon pathogen infection. Conversely, the *atrav1* mutant plants were unable to induce the expression of these defense genes and were highly susceptible to infection. Our data suggests that upon pathogen attack, AtRAV1 transcriptionally upregulate the expression of *MAPKs* (*AtMPK3, AtMPK4* and *AtMPK6*) and AtMPK3 and AtMPK6 are essential for AtRAV1 mediated disease resistance. Further, we demonstrate that AtRAV1 is a phosphorylation target of AtMPK3 (but not AtMPK6) and the phospho-defective variants of AtRAV1 are unable to induce disease resistance in *A. thaliana*. Considering the presence of AtRAV1 orthologs in diverse plant species, we propose that they can be gainfully deployed to control economically important diseases. In deed we observe that overexpression of tomato ortholog of *AtRAV1* (*SlRAV1*) provides broad spectrum disease resistance against bacterial (*Ralstonia solanacearum)*, fungal (*R. solani*) and viral (Tomato leaf curl virus) infections in tomato.

---

Plants have evolved specific receptors to perceive pathogen attack. The PRRs (Pattern recognition receptors) is deployed to perceive pathogen associated molecular cues while leucine rich repeat (LRR) receptors recognize effector proteins of the pathogens (1–3). In this process, plant mount strong defense response to ward off most of the pathogens (1, 4–6). Increase in production of reactive oxygen species (oxidative burst), alkalization of cytoplasm, production of phenolics, phytoalexins, deposition of lignin and callose, hypersensitive response associated programmed cell death, etc are part of plant defense strategies (7). The phytohormones such as jasmonic acid, salicylic acid and ethylene also play a critical role in elaborating the plant defense response (8–12). Moreover, an extensive crosstalk (both synergistic and antagonistic) between various phytohormones modulate the defense response (13, 14).

On the other hand, for successful colonization phytopathogens have evolved diverse strategies to suppress the induction of plant defense response. With extensive polymorphisms in various isolates/strains, some phytopathogens are able to cause disease on diverse host species. *Ralstonia solanacearum* is one of the notable examples which causes devastating bacterial wilt disease in tomato, potato and over two hundred other plant species (15–17). Similarly *Rhizoctonia solani* a necrotrophic fungal pathogen infects diverse plants including rice, potato, tomato etc. and imparts huge economic losses (18, 19). Notably *R. solani* and *R. solanacearum* share many common hosts, including agriculturally important crops such as tomato, potato, etc. Moreover viruses also pose a serious threat for crop production (20). Thus, strategy to simultaneously control bacterial, fungal as well as viral diseases in an eco-friendly and sustainable fashion will be important for ensuring food security.

Manipulation of some of the PRR receptors, LRR receptors and host defense related genes had been shown to provide broad spectrum disease resistance in plants (21, 22). The overexpression of *AtNPR1* (Non expressor of *PR* genes; encoding a positive regulator of SAR) provided broad spectrum disease resistance in various crop plants (23, 24). Similarly, the overexpression of an *AtEFR* (a PRR receptor) gene could enhance tolerance against bacterial pathogen (*R. solanacearum, Xanthomonas perforans*) infection in tomato (25). Further, overexpression of an anti-apoptotic vaculoviral p35 protein also imparted broad spectrum disease resistance in tomato against bacterial (*Pseudomonas syringae* pv. tomato) and fungal (*Alternaria alternata* and *Colletotrichum coccodes*) infections (26). However, the deployment of transgenes in disease management has to face strong biosafety regulations (27). In this regard, utilization of endogenous gene(s) with broad spectrum disease resistance will be helpful in preventing yield loss due to pathogen attack.

In this study, we endeavoured to identify master regulator of plant defense genes in model plant *A. thaliana* and explore the potential of identified gene to impart broad spectrum disease resistance in economically important crops such as tomato. Based upon network centrality parameters, we identified 16 proteins to be topologically central to Arabidopsis defense proteins interaction network. The RAV1 transcription factor binding motifs was present in the promoter region of each of the genes encoding them. It is worth mentioning that RAV1 is an ethylene responsive transcription factor which contains AP2 domain (which participates in activation of ethylene mediated signalling pathway) at its N-terminal region and B3 domain (involved in abscisic acid mediated signalling) at its C-terminus (28, 29). RAV1 has been shown to be a positive regulator of leaf senescence (30) and upon overexpression it provides ABA insensitive phenotype in *A. thaliana* (29).

In this study we identified AtRAV1 as a master regulator of defense gene expression in *A. thaliana* including mitogen activated protein kinases (MAPKs; AtMPK3, AtMPK4 and AtMPK6) and when overexpressed it provides disease resistance against *R. solani*. Similarly, overexpression of *SlRAV1* (ortholog of *AtRAV1*) confers broad spectrum disease resistance in tomato against fungal (*R. solani)*, bacterial (*R. solanacearum*) and viral (tomato leaf curl Joydebpur virus; *ToLCJoV*) infections. The data presented in this study highlights a novel cross talk between RAV1 with MAPKs in imparting disease resistance. The RAV1 transcriptionally induces the expression of *MAPKs* (*AtMPK3, AtMPK4* and *AtMPK6*) and the AtMPK3/AtMPK6 is essential for RAV1 mediated disease resistance. Further the AtMPK3 (but not AtMPK6) phosphorylates AtRAV1 and potentially stabilize it to facilitate sustained activation of defense response.

## Results

### Identification of *AtRAV1* as a key transcriptional regulator of plant defense genes

*In-silico* analysis of protein-protein interactions between Arabidopsis defense proteins (31); identified 16 proteins to be important for the topology and dynamics of the network (**Fig. S1)**. Here onwards, we refer these proteins as key defense proteins of Arabidopsis. Some of the previously reported plant defense proteins such as SKP1 (32), MAPKs (33, 34), heat shock proteins (35, 36) and cyclophilins (37, 38) were noteworthy in this list. Interestingly, the RAV1 binding sites were present in the promoter region of each of these genes (**Fig. S2**). The phylogenetic analysis revealed RAV1 to be conserved in different monocot as well as dicot plants (**Fig. S3**). Presence of RAV1 binding motifs in the *AtRAV1* promoter (**Fig. S4**) suggested it to be under auto-regulation.

### Overexpression of *AtRAV1* induces the expression of key defence genes

We reasoned that overexpression of *AtRAV1* would simultaneously induce the key defense genes and promote disease resistance in *A. thaliana*. To test this, transgenic *A. thaliana* (Col-0) lines that constitutively overexpress *AtRAV1* (At1G13260) under CaMV 35S promoter were generated. Two independent overexpression lines (OE1 and OE2) having relatively higher fold expression of *AtRAV1* along with an EV line were propagated to T_4_ generation for further analysis (**Fig. S5**). Compared to EV plants, the OE lines (OE1 and OE2) demonstrated enhanced expression of key defense genes (**Fig. 1A**).

**Fig. 1.**
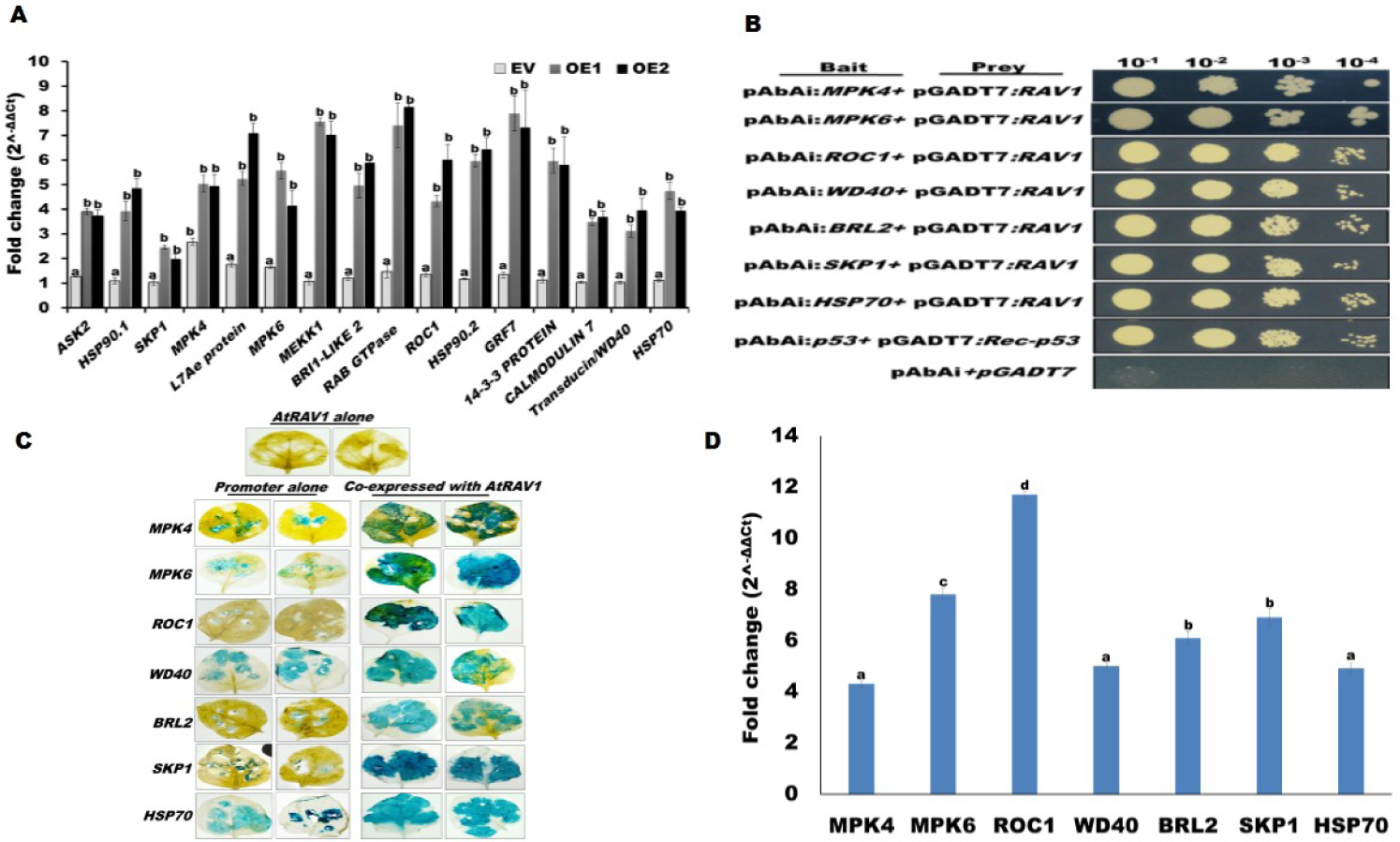
*AtRAV1* up-regulates the expression of key defence genes. (A) Relative expression pattern of sixteen key defense genes in *A. thaliana*. The differential expression of these genes in *AtRAV1* overexpression lines (OE1 and OE2) with respect to the wild type (WT) plants was calculated using beta actin gene as endogenous control. (B) Yeast one hybrid (Y1H) based transactivation assay. The full-length *AtRAV1* were ligated into the pGADT7 (Prey) vector and promoters of selected key defence genes were cloned in pAbAi bait vectors in upstream of Aureobasidine A (AbA). The growth of co-transformed (Prey+Bait) yeast strain on SD/-Leu/-Ura/AbA verifies the transactivation activity. (C) GUS reporter assay. Transcriptionally fused *GUS* under the promoter of selected key defense genes was found induced (appearance of blue color) when co-expressed with AtRAV1 in *N. benthamiana*. (D) qRT-PCR quantification of GUS gene expression in *N. benthamiana* leaves. The *Npt*II gene was used as internal control and relative expression was quantified upon normalization with promoter: GUS infiltrated samples. Graph shows mean values ± standard error of at least three technical replicates. For each gene, different letters indicate significant difference at *P*<*0.05* (estimated using one-way ANOVA). Similar results were obtained in at least three biological repeats.

To validate that AtRAV1 can bind to the promoter and induces expression, we randomly selected few of key defense genes namely *MPK4, MPK6, ROC1, WD40, BRL2, SKP1* and *HSP70* and performed yeast one hybrid (Y1H) as well as GUS reporter assays. For Y1H assay potential AtRAV1 binding motifs in the promoter region of these genes (**Table S1**) were individually cloned in a bait vector (pAbAi) while the full length *AtRAV1* was cloned in a prey vector (pGADT7-AD). The Y1H Gold bait reporter yeast strain expressing both the plasmids grew on Aureobasidin A (AbA) containing double drop out (SD-URA-LEU) plates while the strain co-expressing empty vectors (pAbAi and pGADT17) failed to grow on such plates (**Fig. 1B**). GUS reporter assay was performed in *N. benthamiana* plants to validate that co-expression of AtRAV1 modulates GUS expression through the promoters of selected key defense genes. Limited GUS expression was observed in pBI101:promoter:*GUS* infiltrated leaves, while significantly enhanced GUS expression was observed when promoter:*GUS* and *AtRAV1* constructs were co-infiltrated (**Fig. 1C**). The qRT-PCR further reinforced that expression of *AtRAV1* enhances *GUS* expression (**Fig. 1D**). Taken together, our result suggests that AtRAV1 modules the expression of various key defense genes including AtMPK4 and AtMPK6. As AtMPK3 is an important player in plant defense (39), we performed Y1H and promoter:*GUS* reporter assays to test whether AtRAV1 can modulate *AtMPK3* gene expression. As shown in **Fig. S6**, the AtRAV1 did bind to the promoter of *AtMPK3* and induced its expression.

### Overexpression of *AtRAV1* confers disease resistance in *A. thaliana*

We further analysed whether the overexpression of *AtRAV1* enhances disease resistance against *Rhizoctonia solani*, a notorious necrotrophic fungal pathogen infection. Both OE1 and OE2 lines demonstrated only mild necrotic symptoms when infected with *R. solani*, however severe necrosis was observed in the infected *atrav1* mutant (a previously reported mutant line *Salk_021865*; obtained from Arabidopsis Biological Resource Center, ABRC; **Fig. S7**), EV as well as WT plants (**Fig. 2A**). Compared to others the extent of host cell death (**Fig. 2B**) and ROS accumulation was relatively less in the infected OE lines (**Fig. 2C**). Moreover, the disease severity index (**Fig. 2D**) and abundance of fungal (estimated through monitoring the abundance of *R. solani* 18S ribosomal gene through qRT-PCR) biomass (**Fig. 2E**) was significantly less in infected OEs plants, compared to the infected *atrav1* mutant, WT and EV plants. Also the chlorophyll content was relatively higher in infected OE lines compared to that of WT, EV and atrav1 mutant plants (**Fig. 2F**).The confocal microscopic analysis revealed limited growth of *R. solani* and absence of infection cushion in OE lines (**Fig. 2G**). Taken together, these results reinforced that overexpression of *AtRAV1* imparts enhanced resistance against *R. solani* infection.

**Fig. 2.**
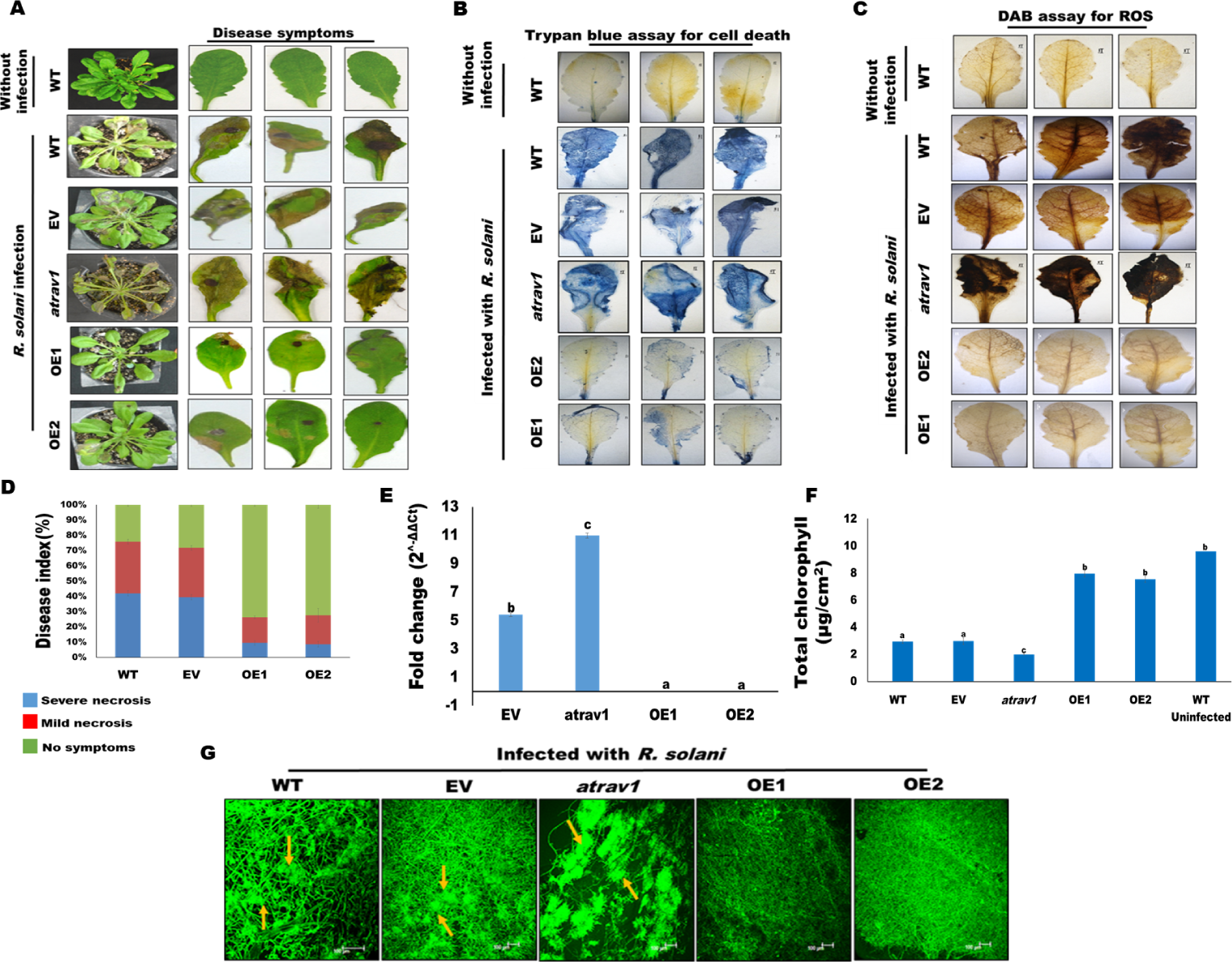
Overexpression of *AtRAV1* provides disease resistance against *R. solani* infection in *A. thaliana*. (A) Disease symptoms observed as brown necrotic lesions in infected leaves at 4 dpi. (B) Trypan blue staining for visualization of cell death and (C) DAB staining for ROS accumulation (brown coloration) in the infected leaves. (D) Disease index (based upon observed necrotic symptoms). (E) Bar graph showing qRT-PCR quantification of 18S RNA gene of *R. solani* (reflecting pathogen load). (F) Total chlorophyll content in *R. solani* infected *A. thaliana* leaves. (G) Confocal imaging of WGA-FITC stained *R. solani* mycelium in the infected leaves. The arrows indicate infection cushions in WT and EV plants. Graph shows mean values ± standard error of at least three technical replicates. Values with different letters are significantly different at *P*<*0.05* (estimated using one-way ANOVA). Similar results were obtained in at least three biological repeats.

### Expression of key defense genes gets enhanced upon pathogen infection in *AtRAV1* overexpressing lines

In comparison to WT and EV plants, the expression of *AtRAV1* was up-regulated upon pathogen *(R. solani*) infection in OE lines but not in the *atrav1* mutant lines (**Fig. S8**). Similarly, the expression of most of the selected key defense genes (*BRL2, ROC1, SKP1, WD40* and *HSP70*) as well as previously reported Salicylic acid (SA), Jasmonic acid (JA) and Ethylene (ET) mediated defense marker genes (**Table S2**) were significantly enhanced upon *R. solani* infection in the OE lines but not in the *atrav1* mutant plants (**Fig. S9A and B**). Also the enhanced expression of *AtMPK3, AtMPK4* and *AtMPK6* was observed in *R. solani* infected OE lines (**Fig. 3A**). Western blot analysis further revealed enhanced accumulation of AtMPK3, AtMPK4 and AtMPK6 proteins in the infected OE lines but not in the atrav1 mutant plants (**Fig. 3B**). Here it is worth mentioning that compared to AtMPK4, the extent of up-regulation of AtMPK3 and AtMPK6 was significantly high (**Fig. 3)**.

**Fig. 3.**
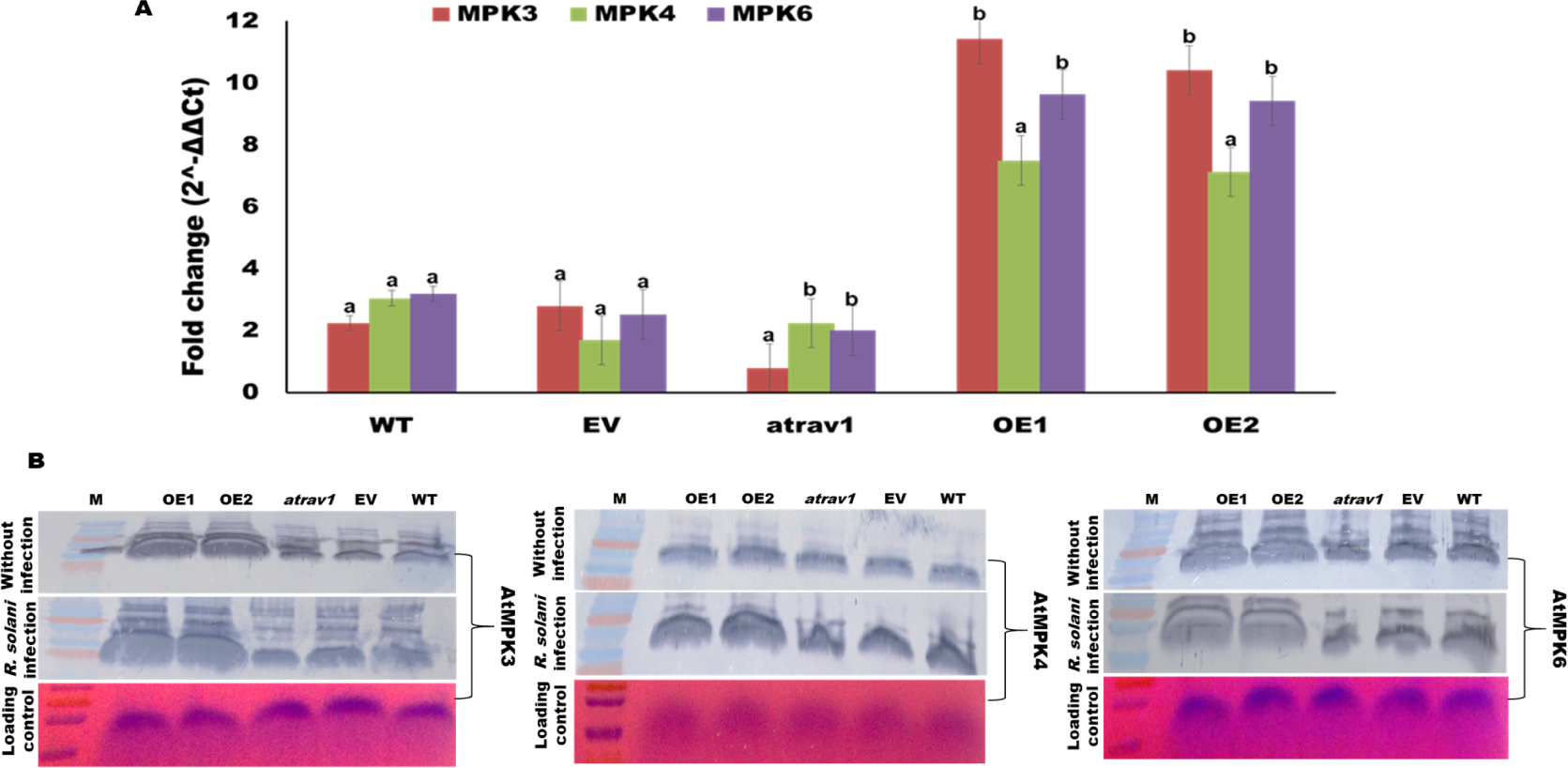
The MAP kinases are induced in *R. solani* infected *AtRAV1* overexpressing lines. (A) Bar graph represents qRT-PCR based expression analysis of different *MAP* kinase genes in *R.solani* infected *A. thaliana* leaves. The relative expression was quantified by normalizing the expression with uninfected samples using beta actin as endogenous control. (B) Western blot analysis showing the expression of AtMPK3, AtMPK4 and AtMPK6 proteins in different *A. thaliana* plants with or without *R. solani* infection. Graph shows mean values ± standard error of at least three technical replicates. For each gene, different letters indicate significant difference at *P*<*0.05* (estimated using one-way ANOVA). Similar results were obtained in at least three biological repeats.

### AtMPK3 and AtMPK6 are required for AtRAV1 mediated disease resistance in *A. thaliana*

We obtained *AtMPK3* (*atmpk3*, SALK_100651), *AtMPK4* (*atmpk4-2*, SALK_056245) and *AtMPK6* (*atmpk6-2*, SALK_073907) mutants from ABRC stock centre and subjected them to *R. solani* infection. The *mpk3* and *mpk6-2* mutants were hyper susceptible to *R. solani* infection while the *mpk4-2* mutant was moderately susceptible (**Fig. 4A**). We crossed each of MAP kinase mutants individually with the *AtRAV1* OE1 plants to obtain the AtRAV1^OE1^/mpk3, AtRAV1^OE1^/mpk4-2 and AtRAV1^OE1^/mpk6-2 lines; wherein the respective MAP kinase protein has been knocked out (**Fig. S10**). Interestingly, the AtRAV1^OE1^/ mpk3 as well as AtRAV1^OE1^/ mpk6-2 lines demonstrated hyper susceptibility to *R. solani* infection (**Fig. 4A**). On the other hand the AtRAV1^OE1^/mpk4-2 showed moderate disease tolerance; however the amplitude of tolerance was significantly less compared to that observed in OE1 lines (**Fig. 4A-C**). Also ROS accumulation, extent of host cell death and pathogen load (*R. solani* biomass estimated through qRT-PCR) were significantly high in AtRAV1^OE1^/mpk3 and AtRAV1^OE1^/mpk6-2 lines, compared to the AtRAV1^OE1^/mpk4-2 or OE1 lines (**Fig. 4A and 4B**). The total chlorophyll content of infected AtRAV1^OE1^/mpk3 and AtRAV1^OE1^/ mpk6-2 lines was significantly less compared to that of AtRAV1^OE1^/mpk4-2 or OE1 line (**Fig. 4C**). Taken together these results highlighted that AtMPK3/AtMPK6 is predominantly required for AtRAV1 mediated enhanced disease resistance in *A. thaliana*.

**Fig. 4.**
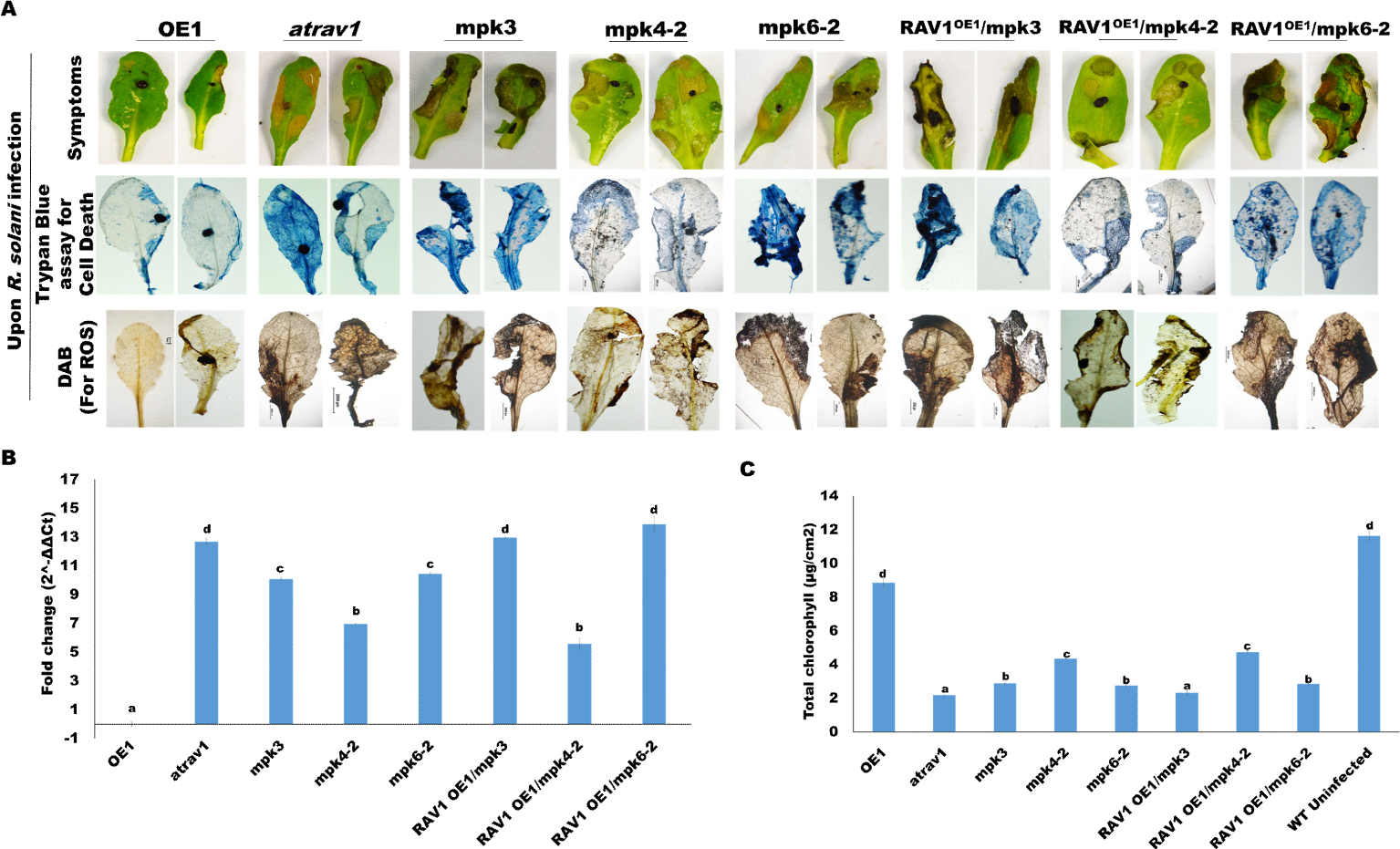
AtRAV1 mediated disease resistance requires functional AtMPK3 and AtMPK6 proteins. (A) The disease symptoms, cell death (trypan blue staining) and ROS accumulation (DAB staining) in the *R. solani* infected leaves of *A. thaliana* plants at 4 dpi. (B) Bar graph showing qRT-PCR based quantification of 18S ribosomal RNA of *R. solani* (reflecting pathogen load) in the infected plants. (C) Total chlorophyll content in the infected *A. thaliana* leaves. Data are reflected as mean ± SE of at least three technical. Values with different letters are significantly different at *P*<*0.05* (estimated using one-way ANOVA). Similar results were obtained in three biological repeats.

### AtRAV1 is phosphorylated by AtMPK3 under in-vitro condition

Bioinformatics analysis revealed that AtRAV1 protein contains three TP and one SP amino acid residues as putative MAP kinase phosphorylation sites (**Fig. S11**). We ectopically overexpressed and purified AtRAV1 protein as well as its different variants wherein potential phosphorylation residues had been mutated (SDM1: Ser310Ala; SDM2: Thr19Ala; SDM3: Thr23Ala; SDM4: Thr193Ala; SDM5: having all four potential phosphorylation sites mutated) from *E. coli* cells to analyse their phosphorylation by AtMPK3 and AtMPK6 under in-vitro condition. The assay revealed that AtMPK3 but not AtMPK6 phosphorylate the AtRAV1 (**Fig. 5A and 5B**). Compared to others, the SDM2 demonstrated weak phosphorylation signal by AtMPK3 whereas the phosphorylation was completely abolished in case of SDM5.

**Fig. 5.**
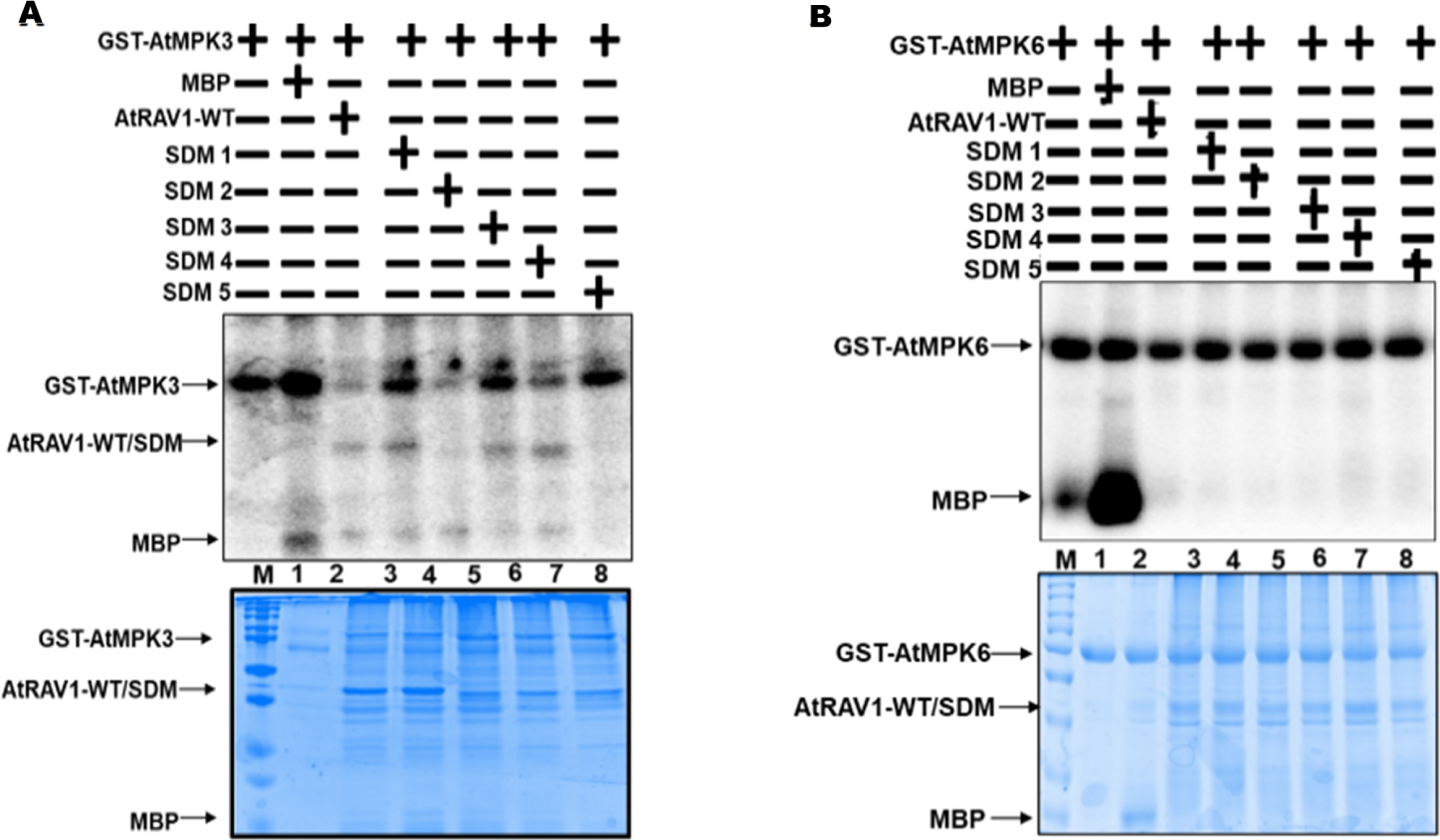
AtRAV1 is phosphorylated by AtMPK3 under in-vitro condition. (A) and (B) Upper panel: Autoradiogram showing in-vitro phosphorylation of bacterially expressed AtRAV1 with AtMPK3-GST and AtMPK6-GST. The phosphorylation of MBP was used as a positive control. (A) and (B) Lower panel: coomassie brilliant blue stained gel (12%) with positions of different proteins indicated by arrows. The plus and minus signs indicate the presence and absence of proteins, respectively during the assay.

### Overexpression of phospho-defective variants of AtRAV1 is unable to induce disease resistance in *A. thaliana*

In order to test whether phosphorylation of AtRAV1 is required for inducing disease resistance, we overexpressed phospho-defective variants of AtRAV1 (SDM2 and SDM5) in the WT and *atrav1* mutant *A. thaliana* plants. Interestingly in both the cases (WT^SDM2^/^SDM5^ and atrav1^SDM2^/^SDM5^) the plants were highly susceptible to *R. solani* infection (**Fig. 6A**). The severity of disease symptoms, extent of host cell death and pathogen load were also significantly higher in WT^SDM2^/^SDM5^ and atrav1^SDM2^/^SDM5^ plants, compared to the *AtRAV1* OE1 plants (**Fig. 6A and 6B**). In contrary to the OE1 line, the WT^SDM2^/^SDM5^ and atrav1^SDM2^/^SDM5^ lines were unable to induce the expression of key defense genes including MAP kinases (*AtMPK3, AtMPK4* and *AtMPK6*) (**Fig. 6C)** and defense marker genes (**Fig. 6D**) upon pathogen infection. Overall, this suggested that phosphorylation of AtRAV1 is required for eliciting defense response and imparting disease resistance in *A. thaliana*.

**Fig. 6.**
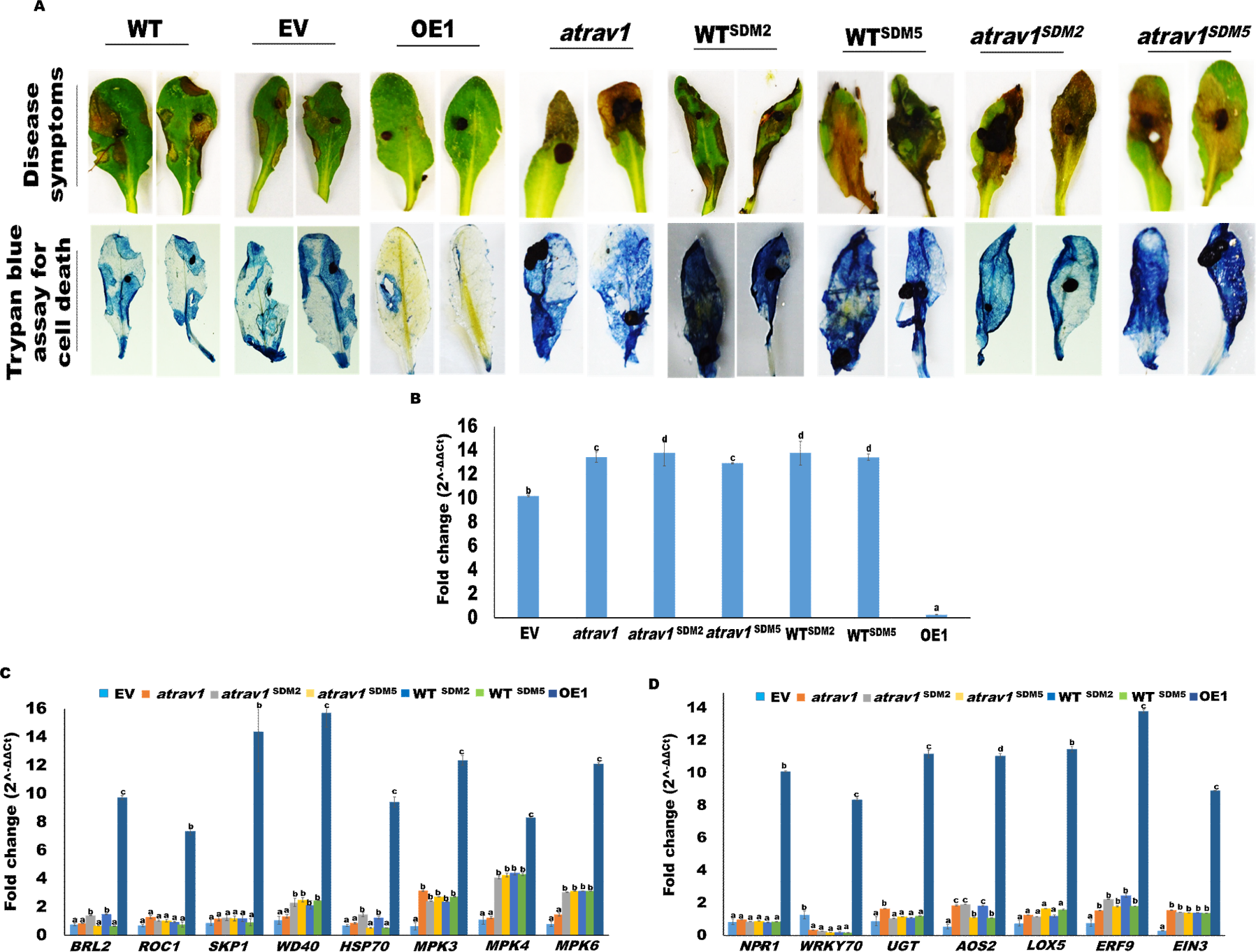
The overexpression of phospho-defective variants of *AtRAV1* fail to impart resistance against *R. solani* infections in *A. thaliana*. (A) The disease symptoms and extent of host cell death (trypan blue staining) in the *R. solani* infected leaves of different A. *thaliana* plants at 4 dpi. (B) Bar graph showing qRT-PCR based quantification of 18S ribosomal RNA of *R. solani* (reflecting pathogen load) in the infected plants. The expression of (C) selected key defense genes and (D) Defense marker genes in *R. solani* infected lines. The relative expression was quantified by normalizing the expression with that of *R. solani* infected wild type plants using beta actin as endogenous control. Graph shows mean values ± standard error of at least three technical replicates. For each gene, different letters indicate significant difference at *P*<*0.05* (estimated using one-way ANOVA). Similar results were obtained in two different biological repeats.

### Overexpression of *SlRAV1* imparts broad spectrum disease resistance in tomato

To further substantiate the role of RAV1 in plant defense, we analysed whether the *AtRAV1* ortholog of tomato (*SlRAV1;* EU164416) can impart disease resistance in tomato (*Lycopersicon esculentum* mill). Two independent *SlRAV1* OE (OE:L1 and OE:L2) lines were generated in tomato cultivar Pusa Ruby (**Fig. S12**). The qRT-PCR revealed both the OE lines to have higher fold expression of *SlRAV1* (**Fig. S12E**) and the expression were significantly enhanced upon *R. solani* infection in OE lines (**Fig. S13A**). The western blot analysis also suggested enhanced accumulation of SlRAV1 in pathogen infected OE lines (**Fig. S13B**). The disease symptoms (**Fig. 7A**), disease severity index (**Fig. 7B**) and chlorophyll content of the infected leaves (**Fig. 7C**) suggested enhanced disease tolerance in OE lines, compared to EV and WT tomato plants. Also, the expression of most of the selected Salicylic acid (SA), Jasmonic acid (JA) and Ethylene (ET) mediated defense marker genes were significantly enhanced upon *R. solani* infection in the OE lines (**Fig. 7D**).

**Fig. 7.**
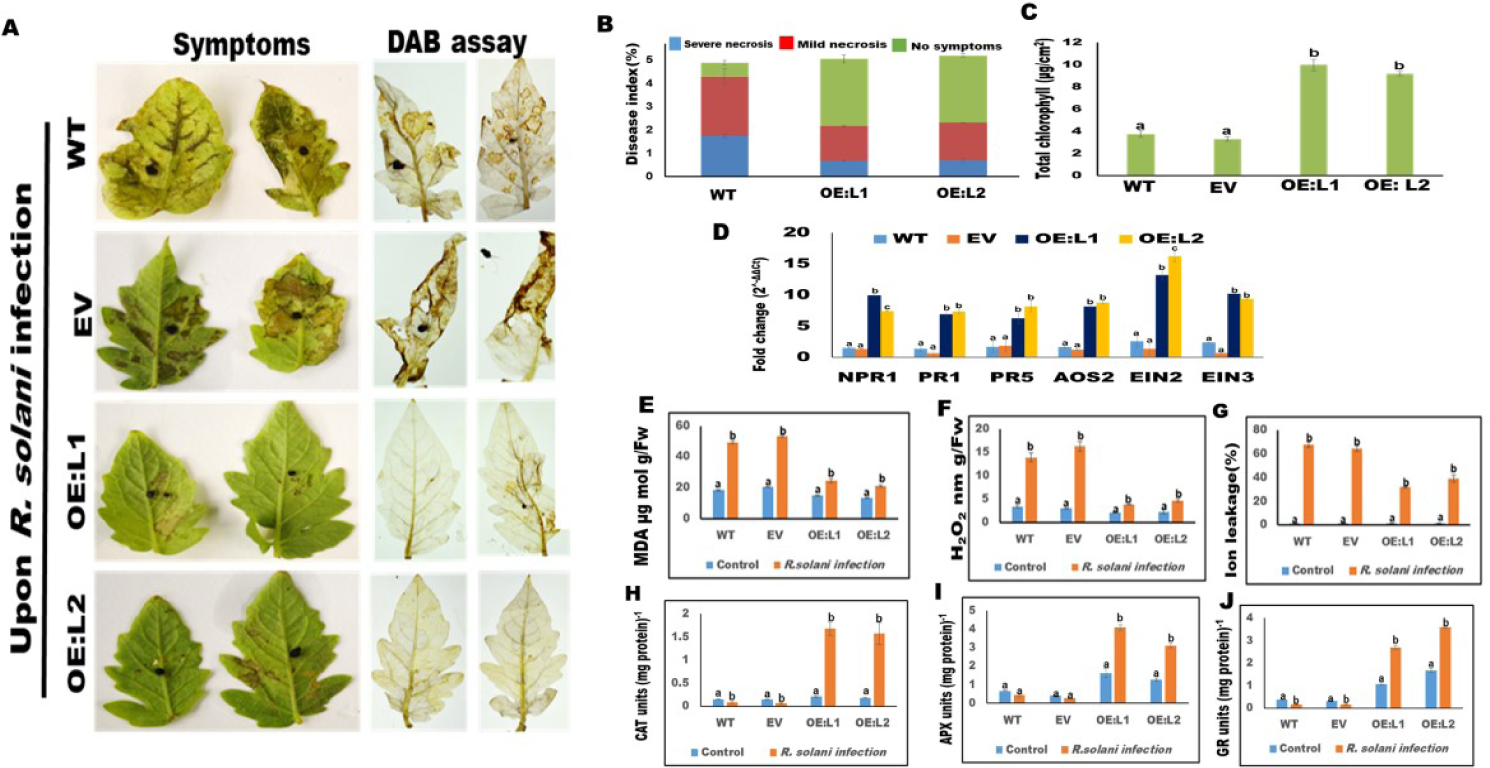
Overexpression of *SlRAV1* provides resistance against *R. solani* infection in tomato. (A) Disease symptoms, (B) Disease index (in terms observed necrotic symptoms) and (C) Total chlorophyll content in the *R. solani* infected tomato leaves at 4 dpi. (D) Expression analysis of SA, JA and ET mediated marker genes in the infected samples. The relative expression was quantified with respect to uninfected samples using beta actin as internal control. (E) MDA content, (F) H_2_O_2_ content, (G) ion leakage (%) and the enzymatic activities of various antioxidant markers, (H) CAT, (I) APX and (J) GR in the infected plants. Graph shows mean values ± standard error of at least three technical replicates. For each gene, different letters indicate significant difference at *P*<*0.05* (estimated using one-way ANOVA). Similar results were obtained in at least three biological repeats.

We further tested the susceptibility of OE lines against a deadly pathogen (*Ralstonia solanacearum*) which causes bacterial wilt disease in tomato. As shown in **Fig. 8A**, the wilting symptoms was remarkably less in infected OE lines (OE:L1 and OE:L2), compared to WT plants. Notably, the pathogen (*R. solanacearum*) load and disease severity index were also significantly less in the infected OE:L1 and OE:L2 lines (**Fig. 8B** and 8**C**). The enhanced accumulation of SlRAV1 protein in *R. solanacearum* (**Fig. 8D**) infected tomato OE lines, supported its pathogen inducible nature. Also gene expression (**Fig S14**) as well as enzymatic activity of some of the antioxidant markers such as catalase (CAT), ascorbate peroxidase (APX) and glutathione reductase (GR) were significantly higher in *R. solani* (**Fig. 7H-J**) and *R. solanacearum* (**Fig. 8H-J**) infected OE lines while the MDA content, H_2_O_2_ content and ion leakage were reduced in these lines (**Fig. 7E-G** and **8E-G**).

**Fig. 8.**
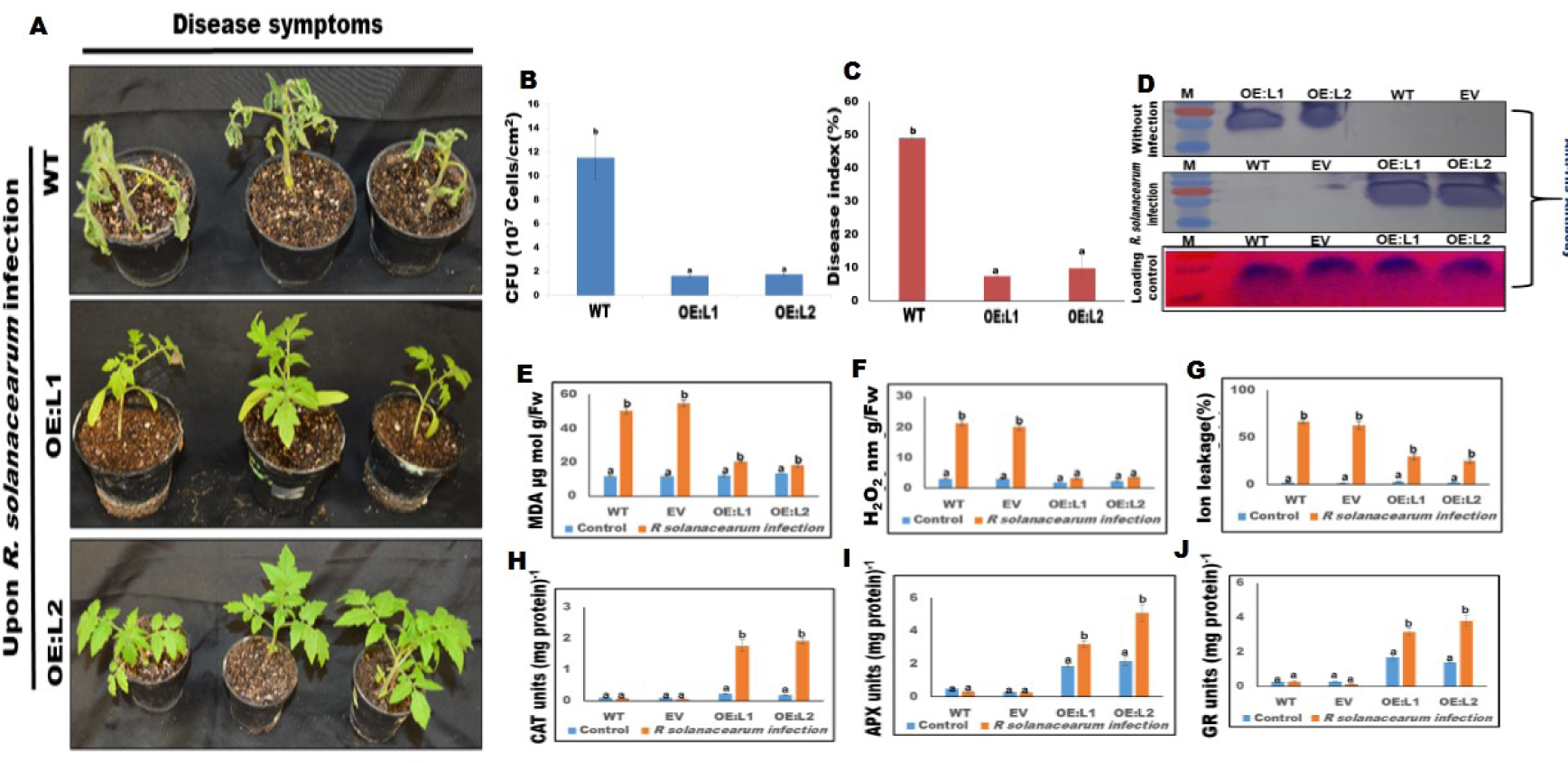
Overexpression of *SlRAV1* provides resistance against *R. solanacearum* infection in tomato. (A) Disease symptoms, (B) The pathogen load (CFU/ml) and (C) disease index (% of plants with wilting symptoms) in drench inoculated *R. solanacearum* infected tomato plants at 7dpi. (D) Western-blot analysis reflecting the accumulation of His-tagged SlRAV1 protein in tomato plants with or without *R. solanacearum* infection. (E) MDA content, (F) H_2_O_2_ content, (G) ion leakage (%) and the enzymatic activities of various antioxidant markers, (H) CAT, (I) APX and (J) GR in the infected plants. Graph shows mean values ± standard error of at least three technical replicates. For each gene, different letters indicate significant difference at *P*<*0.05* (estimated using one-way ANOVA). Similar results were obtained in three biological repeats.

We further observed the OE lines to have high level of tolerance against Tomato Leaf Curl Joydepur Virus (*ToLCJoV*) infection. The leaf curling symptom was negligible in the OE lines but severe in case of WT as well as EV plants (**Fig. S15A**). ROS accumulation (**Fig. S15B**), disease severity index (**Fig. S15C**) and total chlorophyll content (**Fig. S15D**) further reinforced enhanced tolerance against *ToLCJoV* infection in OE lines.

## Discussion

Plants are susceptible to various bacterial, fungal and viral diseases. Controlling them in an eco-friendly manner is a challenge for sustainable agriculture. We endeavoured to identify gene(s) that can provide broad spectrum disease resistance in plants. Initially, while studying the defense protein interaction network, we identified RAV1, an ethylene responsive transcription factor as a master transcriptional regulator of defense genes in *Arabidopsis thaliana*. Previous studies had shown that overexpression of pepper RAV1 provides resistance against *Pseudomonas syringae* pv. tomato DC3000 (a hemi-biotrophic bacterial pathogen) infection in *A. thaliana* by induction of PR genes (40). Here we observed that overexpression of *AtRAV1* confers remarkable resistance against fungal (*Rhizoctonia solani*) infection in *A. thaliana*. Similarly, overexpression of *SlRAV1* (the *AtRAV1* ortholog in tomato) imparted remarkable level of protection against fungal (*R. solani*), bacterial (*R. solanacearum*) and viral (Tomato leaf curl virus) diseases in tomato. We observed that AtRAV1 acts as a master transcriptional activator of various key defense genes (that are topologically central to defense protein interaction network) as well as JA, SA and ET responsive defense marker genes in *A. thaliana*. The enhanced expression of *AtMPK3, AtMPK4* and *AtMPK6* transcripts as well as their proteins in the pathogen infected OE lines suggests the activation of MAP kinase signalling. The AtMPK3/AtMPK6 signalling seems essential for mediating AtRAV1 mediated defense; as knocking out AtMPK3 or AtMPK6 (but not AtMPK4) rendered the *AtRAV1* OE lines hyper susceptible to *solani* infection. The MPK3/MPK6 mediated signalling plays an important role in elucidation of pathogen triggered immunity (PTI) and effector triggered immunity (ETI) as well as production of defense associated phytoalexin, camalexin in plants (41–44). Also the MPK3/MPK6 are known to phosphorylate some plant transcription factors and regulate important cellular response including defense response (45). Notably, the AtMPK3/AtMPK6 mediated phosphorylation of some members of ERF family (ERF6/ERF104) assist in elucidation of plant defense response (46). In this study, we observed that AtMPK3 but not AtMPK6 can phosphorylate the AtRAV1 under in-vitro condition. Recently, it has been reported that AtMPK3 but not AtMPK6 regulates submergence tolerance by phosphorylation of SUB1A1 (Submergence1A1) transcription factor (47). We anticipate that the AtMPK3 mediated phosphorylation of AtRAV1 may stabilize the protein during pathogen attack and activates it to stimulate the defense response. Thus on one hand, the AtRAV1 binds to the promoter of different MAP kinases (AtMPK3/AtMPK4/AtMPK6) and induces their signalling while on the other hand AtMPK3 phosphorylates AtRAV1 and modulates its function. In deed we observed that phospho-defective variants of AtRAV1 (SDM2 and SDM5) are unable to impart disease resistance against *R. solani* infection in *A. thaliana*.

There has been trade-off between disease resistance and plant growth (48). The constitutive activation of defense genes can negatively impact plant growth and development (49). However in this study, we did not observe any apparent growth or developmental defects in the RAV1 overexpressing *A. thaliana* as well as tomato lines. We attribute this may be due to observed dynamics of induction of defense genes expression in the OE lines. Although several defense genes including MAP kinases were overexpressed in OE lines, the extent of their up-regulation was further enhanced upon pathogen infections. Similarly, the antioxidant machinery of the host (tomato) was significantly enhanced upon pathogen (*R. solani* and *R. solanacearum*) infection in OE lines. Such dynamic expression of defense genes may ensure strong protection during pathogen attack while averting the negative effect of induced defense responses under control (uninfected) conditions. Further, due to a relatively short (<60 min) half-life (50) and potential ability to auto-regulate its own transcription, the level of RAV1 is limited under normal conditions. Conversely with pathogen inducible nature, the RAV1 level gets enhanced in the infected OE lines and upon potential phosphorylation by AtMPK3 the protein gets stabilized, leading to sustained activation of MPK3/MPK6 signalling and thereby induction of defense genes. In support of this, we observed limited induction of defense genes including MAPKs in the *A. thaliana* lines overexpressing the phosphor-defective variants of AtRAV1 (**Fig. 6**).

Overall, the present study reports RAV1, an ethylene responsive transcription factor as a master regulator of plant defense and is a novel phosphorylation target of AtMPK3. A cross talk between RAV1 and MAPKs is required for inducing disease resistance against *R. solani* infection in *A. thaliana*. Furthermore, the overexpression of tomato RAV1 provides remarkable level of protection against bacterial, fungal and viral infections in tomato. Considering that RAV1 orthologs are conserved in different monocot and dicot plants and we do not observed any trade-off between enhanced disease resistance with apparent growth/developmental defects in the overexpression *A. thaliana* and tomato lines, our study emphasizes that RAV1 can be gainfully deployed as a biotechnological intervention to develop broad spectrum disease resistance in a variety of crops.

## Methods

### Identification of key defense proteins in *Arabidopsis thaliana*

Mukhtar and colleagues have reported the list of plant (*A. thaliana*) and pathogen proteins (*Pseudomonas syringae* and *Hyaloperonospora arabidopsidis*) that are involved in plant pathogen interactions (51). The plant proteins (n=392 proteins) potentially involved in defense responses were obtained from that list (**Table S3**) and were mapped on to AtPIN (*A. thaliana* Protein Interaction Network) database (52) to construct a Arabidopsis Defense Protein Interaction Network (ADPIN). Visualization of protein-protein interaction network was performed using Cytoscape (53). Furthermore the ADPINv1 (The Arabidopsis Defense Proteins Interaction Network vicinity 1) was constructed by extending ADPIN to include its first interacting partners.

Network centrality measures were computed for ADPINv1 proteins by graph-theoretical analysis. Degree (hubs), betweeness (bottlenecks) and average shortest path (swift communicators) were studied to identify proteins that are central to the interactome and might be critical for executing plant defense. Top 10 proteins with the best values for each of these network parameters, were selected. Notably few proteins were commonly predicted to have best values for each of the network parameter. After removing these redundant proteins from the list, 16 unique proteins, topologically and dynamically central to the network, were obtained (**Table S3**). These proteins are considered as key defense proteins. Gene Investigator tool (https://genevestigator.com/) revealed that genes encoding these proteins are induced during biotic stress (**Table S4**).

### Identification of *AtRAV1* as master transcriptional regulator of key defense genes

The Arabidopsis Information Resources (TAIR) id of each of the 16 identified key defense gene was used as query in Plant PAN database (Plant PAN; http://PlantPAN2.itps.ncku.edu.tw). The transcription start site/5’UTR-End limit was fixed as 100bp, while transcription stop site/3’UTR-End was fixed as 1000bp. The analysis predicted putative transcription binding sites present in the 5’UTR (promoter) region in each of the genes. The *AtRAV1* (At1G13260) transcriptional factor binding sites were common in each of them.

### Plant Materials and Growth Conditions

Different *A. thaliana* (Colombia-0; wild type, transgenic and Salk_ 021865 mutant) lines used in this study were grown on soilrite at 22°C with 8/16-h photoperiod, 70 % relative humidity in growth chamber. Similarly, different tomato (*S. lycopersicum;* cultivar Pusa ruby) plants and tobacco (*N. benthamiana*) plants were grown on soilrite at 26°C with 12/12-h photoperiod, 70 % relative humidity in growth chamber.

### Generation of *AtRAV1* overexpressing *A. thaliana* and *SlRAV1* overexpressing tomato

The CDS of *AtRAV1* (AT1G13260) gene was cloned in pGJ100 (a modified pBin19 binary vector having MCS of pBSKS vector) and transformed into GV3101 strain of Agrobacterium. The Agrobacterium harbouring 35S:*RAV1* construct was inoculated into three weeks old *A. thaliana* (Col-0) plants by floral dip method (54). Presence of transgene was reconfirmed by PCR using *CaMV35S F* and *RAV1OX-R* primer. The *NptII-F* and *NptII-R* primer pair was further used to confirm the presence of vector in both OE and EV lines. The expression of *AtRAV1* gene in different *A. thaliana* lines was verified by qRT PCR using primers mentioned in **Table S5**.

Similarly, the full length gene sequence of *SlRAV1* (EU164416) was cloned into gateway binary vector (pGWB408) using SlRAV1OX-F and SlRAV1OX-R primer pairs (**Table S5**) and transgenic tomato lines (OE:L1 and OE:L2) were generated through Agrobacterium (GV3101 strain) mediated transformation. The presence of transgene was confirmed by PCR using CaMV35S F and SlRAV1OX-R primer while T-DNA was verified by NptII*-F* and NptII*-R* primer. The expression of *SlRAV1* was validated by qRT-PCR using gene specific (SlRAV1 RTF and SlRAV1 RTR) primer. Total crude protein was isolated from 100 mg of plant leaves using P-PER plant protein extraction kit (Thermo scientific: 89803) and the western blot was carried out by electro blotting the 20µg of total crude protein onto polyvinylidene fluoride (PVDF) membrane and probed with mouse polyclonal anti-His-antibody (1:30000 dilutions). The blot was developed as per manufacturer’s protocol (Sigma-Aldrich, Japan).

### Generation of MAP kinase mutants in AtRAV1 overexpressing OE1 lines

The *AtMPK3* (*atmpk3*, SALK_100651), *AtMPK4* (*atmpk4-2*, SALK_056245) and *AtMPK6* (*atmpk6-2*, SALK_073907) mutants were obtained from ABRC stock centre. Presence of T-DNA insertion in mutant (*atmpk3, atmpk4-2 and atmpk6-2)* was individually conformed by PCR using T-DNA border primer and MAP kinase gene specific reverse primer (BP+RP) (**Table S5**). The cross between the MAP kinase mutant plants (as male plant) and the AtRAV1 OE1 (as female) was set up and obtained seeds were grown on soilrite. Upon PCR validation of presence of AtRAV1 overexpression construct (using CaMV35S_forwerd and AtRAV1 reverse primer) and T-DNA insertion in particular MAP kinase (using BP+RP primer), we propagated the seeds to T2 generation and used them for further studies.

### Yeast one hybrid assay

Y1H assay was performed using Matchmaker Gold Yeast One-Hybrid Library Screening System (Clontech, USA). The nucleotide sequences of promoter region of selected key defense genes having potential RAV1 binding motifs were retrieved by using online tool (https://bioinformatics.psb.ugent.be/plaza/). The RAV1 motif enriched promoter sequence of each of the gene (**Table S1**) was cloned in pAbAi bait vector (Clontech, USA) in the upstream of Aureobasidine A. The plasmid was linearized with *Bst*BI restriction enzyme and transformed into Y1H Gold yeast strain, as per manufacturer’s protocol and the positive transformants (Y1H-Bait) was selected on AbA (200mg ml^-1^ AbA) plate. Subsequently the full-length copy of *AtRAV1* was cloned in pGADT7-AD (Clontech, USA) prey vector as GAL4 transcription activation domain (GAL4 AD) fusion protein. It was transformed in the Y1H-Bait strain and positive transformants (Y1H-Bait+Prey) were selected by growing serially diluted (10^−1^, 10^−2^, 10^−3^ and 10^−4^) cells at 30°C for 3 days on AbA (200 mg/ml) containing double drop out (SD-URA-LEU) plates.

### GUS based reporter assay

The promoter region of the selected key defense genes (as described above) was fused with *GUS* gene in pBI101 (pBI101:promoter:*GUS*) and *AtRAV1* full length gene (AT1G13260) was cloned under CaMV35S promoter in pGJ100 (pGJ100:AtRAV1). The primer used for cloning is enlisted in **Table S5**. Both the recombinant plasmids were individually transformed into GV3101 strain of Agrobacterium and were co-infiltrated into the leaves of *N. benthamiana*. After 48 hours of infiltration, GUS expression was analyzed by staining the leaves with GUS solution (1 mg/ml) for 16 hours at 37°C. Upon destaining in a solution (1:3 ratio of glacial acetic acid: ethanol) for 3 to 4 hrs at room temperature and washing with distilled water (55).

### cDNA synthesis and expression analysis

Total RNA was isolated from plant tissues using RNeasy Plant RNA isolation kit (Qiagen, Valencia, CA). 1µg of total RNA was used for cDNA synthesis using Verso cDNA synthesis kit (Thermo Fisher Scientific Inc, USA), as per the manufacturer’s protocol. qRT-PCR of 16 key defence genes and various defense marker genes (SA, JA and ET) was performed using primers mentioned in **Table S5**. The relative fold change was calculated by using 2^-ΔΔCt^ method (56).

### Pathogen infection assays

*R. solani* AG1-IA (BRS1) strain (18) was used to infect *A. thaliana* and tomato. The *R. solani* sclerotia pre-germinated in potato dextrose broth (PDB, Himedia, Mumbai, India) at 28°C for 6 hours were used to infect the leaves of *A. thaliana* plants. A minimum of three leaves per plant and minimum 10 plants of each line per experiment was infected. However for tomato infection, the detached leaves (n=3) of at least 10 plants were used in each experiments (57). On the basis of observed symptom patterns (severe or mild or no symptoms), we categorised percentage of leaves having particular disease symptom as disease index. Total chlorophyll was calculated per square cm area according to Arnon’s equation as mentioned earlier (58). Further, the *R. solani* biomass in the infected samples was estimated by monitoring the expression of its 18S rRNA gene through qRT-PCR using primers mentioned in **Table S5**.

The deadly bacterial pathogen *Ralstonia solanacearum* strain F1C1 was also used to infect tomato. The pathogen was grown in BG media (Peptone 10g, yeast extract 1g, casamino acid 1g, agar 1.5g per litre) at 28°C for 48 h and 2×10^8^ cfu/ml bacterial cells were drench inoculated in the 3 weeks grown nursery pots of tomato as per method described in (16). A minimum of 25 plants of each lines were infected in each experiment and the experiment was independently repeated twice. Disease symptoms were monitored at 7 dpi and percentage of plants with wilting symptoms was plotted as bar chart. The abundance of *R. solanacearum* was estimated by serial dilution plating method and counting the CFU per gram fresh weight of leaf as described earlier (59).

Also, the Tomato leaf curl Joydebpur virus (*ToLCJoV*) was infected to the 3 weeks old tomato leaves as described (60). The disease symptoms were observed at 21 dpi and the disease index, DAB staining and total chlorophyll content were estimated.

### Microscopic analysis

The infected leaves were harvested and stained with WGA-FITC as described earlier to monitor the growth of *R. solani* mycelia (61). The samples were observed under GFP filter of Confocal Laser Scanning Microscope (AOBS TCS-SP5, Leica, Germany). The images were analysed using LAS AF Version: 2.6.0 build 7266 software.

### Biochemical assays

DAB staining was used for detection of ROS while trypan blue staining was performed to detect cell death assay in *R. solani* infected leaves of *A. thaliana* and tomato leaves (62).

Further, the MDA, H_2_O_2_ content and ion leakage (%) were quantified using the earlier described method (61). Similarly the activities of various antioxidant enzymes (CAT, APX and GR) were estimated using the protocol described in (62).

### Expression and purification of AtRAV1 protein

The *AtRAV1* gene was cloned in pET28a bacterial expression vector using *RAV1OX-F* and *RAV1OX-R* gene specific primers and transformed into *E. coli* (BL-21 strain, DE3-codon^+^) cells. The protein was purified using affinity chromatography (Ni^+2^-NTA) following the method described earlier (57). Similarly, the different variants of *AtRAV1*, having different phosphorylation residues mutated (SDM1: Ser310Ala; SDM2: Thr19Ala; SDM3: Thr23Ala; SDM4: Thr193Ala; SDM5: having all four potential phosphorylation sites mutated) were synthesized commercially (Gene Universal Inc; http://www.geneuniversal.com/) and cloned in pET28a to purify different variant proteins. The western blot was performed by electro blotting protein onto polyvinylidene fluoride (PVDF) membrane and probed with mouse polyclonal anti-His-antibody (1:30000 dilutions). Also western blotting of AtMPK3, AtMPK4 and AtMPK6 was performed using anti-MAP kinase-antibody (1:20000 dilutions) as primary antibody and anti-mouse IgG (Sigma) protein (1:15000 dilutions) as secondary antibody.

### *In-vitro* phosphorylation of AtRAV1 by MAP Kinases

The bacterially purified AtRAV1 protein and its variants (as described above) were used for in-vitro phosphorylation assay. The CDS of *AtMPK3* and *AtMPK6* were cloned in pGEX4t2 vector in-frame with amino-terminal GST tag and transformed into *E. coli* (BL21). Upon 1mM IPTG induction for 4h, the proteins were purified using GST-beads as per manufacturer’s protocols (63). The in-vitro kinase assay was performed as described in (47). Briefly, the MAP kinases and RAV1 variant proteins (1:10) were incubated in a 20µl kinase reaction buffer (25 mM Tris-Cl (pH 7.5), 10 mM MgCl2, 5 mM MnCl2, 1 mM DTT, 1 mM β-glycerol-phosphate, 1 μM Na3VO4, 0.5 mg/ml MBP, 25 μM ATP and 1 μCi [γ^-32^ P] ATP) at 30°C for 30 minutes. The reaction was stopped by addition of 2x-SDS-loading buffer and heating at 95°C for 5 minutes. The samples were fractionated in 10% to 12% SDS-PAGE. The phosphorylation signals were detected by Typhoon phosphor imaging system (GE Health Care, Life Sciences, USA).

### Overexpression of phospho-defective variants of *AtRAV1* in *A. thaliana*

The SDM2 and SDM5 variants of *AtRAV1* which were defective in *in-vitro* phosphorylation by AtMPK3 were PCR amplified and cloned in the pGJ100 plant transformation vector. The constructs were subsequently transformed into GV3101 strain of Agrobacterium and the recombinant bacterial strains were used for transformation in the WT (Col-0) and atrav1 (*Salk_021865)* mutant lines. The transgenic lines were confirmed through PCR and sequence analysis.

### Statistical analysis

One-way analysis of variance was performed using Sigma Plot 12.0 (SPSS, Inc. Chicago, IL, USA) with *P ≤ 0.05* considered statistically significant. The statistical significance is mentioned in the figure legend, wherever required.

## Acknowledgements

We acknowledge Ms Ekta Manglesh for making contributions during early stage of the envisaged research. RK acknowledges CSIR, SRA fellowship. Financial support from the DBT-RA programme in Biotechnology and Life Sciences is gratefully acknowledged by DMS. SG acknowledges SPM fellowship from CSIR. PKB thanks UGC for fellowship. The work has been supported by the research findings from Department of Biotechnology, Govt of India under NIPGR Flagship programme (imparting sheath blight disease tolerance in rice) and NIPGR core research grant. We are grateful to Dr Ramesh Sonti, NIPGR for providing critical comments on the manuscript and helping us to improve its quality. Dr S.K. Ray from Tezpur University for his help in providing F1C1 wild type strain of *Ralstonia solanacearum.* We also thank Dr A.K. Singh, ICAR-IIVR, Varanasi for providing infectious clone of *ToLCJoV*.

## Authors’ contributions

GJ conceived the work and coordinated its progress, GJ as well as GB planned while GB supervised the network analysis which leads to identification of key defense proteins. RKC carried out the detail characterization of *AtRAV1* as well as *SlRAV1* to establish its role in providing broad spectrum disease resistance against bacterial, fungal and viral infection. RK assisted in making various constructs and performed YIH. DMS performed protein purification, antioxidant assays and western blot analysis. SG assisted in confocal microscopic analysis and bioinformatics analysis. PKB performed the *in-vitro* phosphorylation assays, AKS contributed different MAPK mutants and antibodies, SP had provided valuable comments on the manuscript. GJ, RKC and DMS had written the manuscript and all authors have approved the final manuscript.

## Supplementary information

**Fig. S1.**
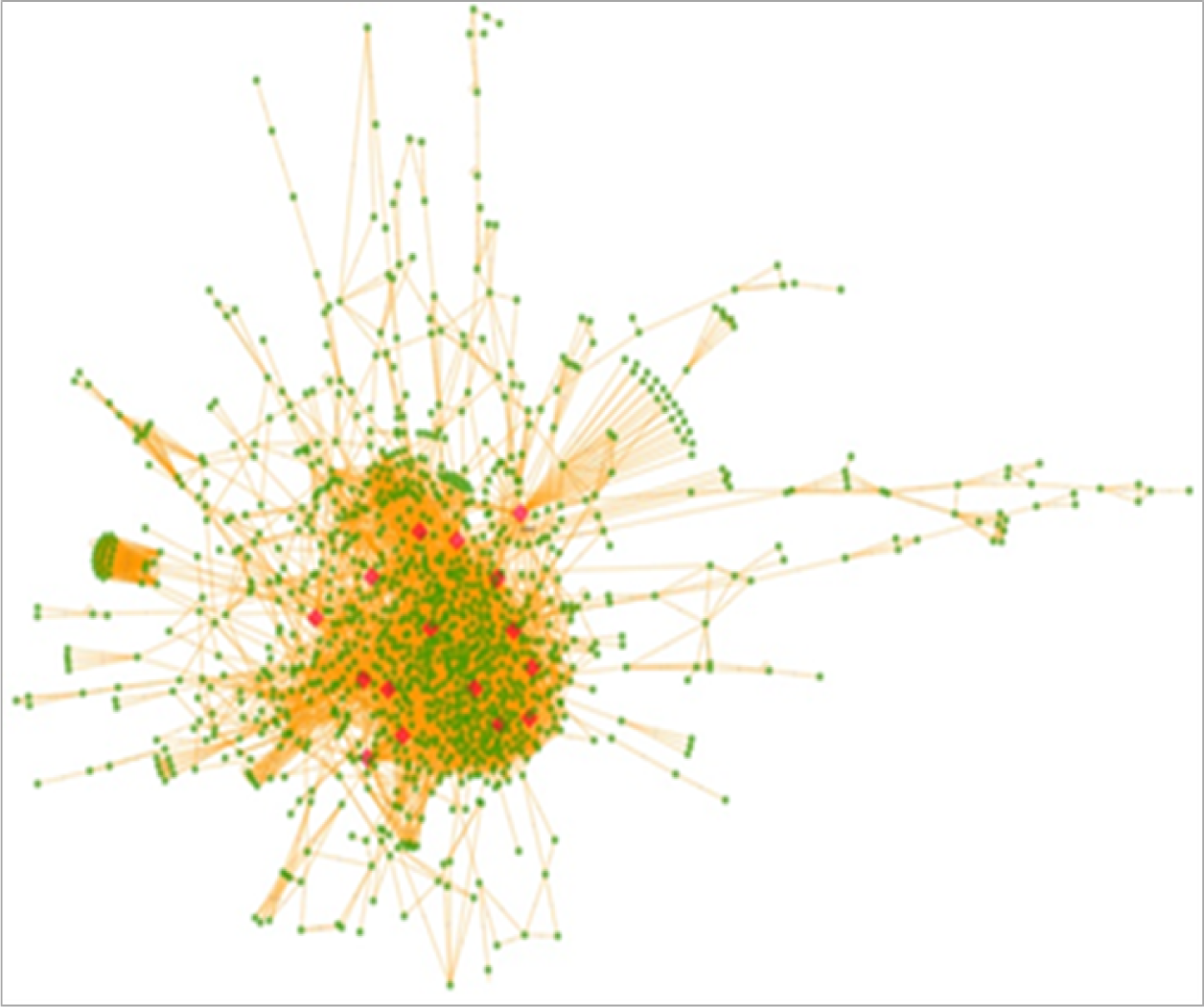
Centrality of key defense proteins in Arabidopsis defense protein interaction network. Protein-protein interaction network of Arabidopsis defense proteins and their immediate interacting partners: ADPINv1. The Arabidopsis defense proteins were mapped onto AtPIN and the interactome of these proteins and their immediate interactors was extracted using Cytoscape. A complex network with 6051 interactions amongst 1343 proteins was observed. Central nature of key plant defense interactome proteins was identified using network metrics. Sixteen key defense proteins are highlighted onto ADPINv1 in red.

**Fig. S2.**
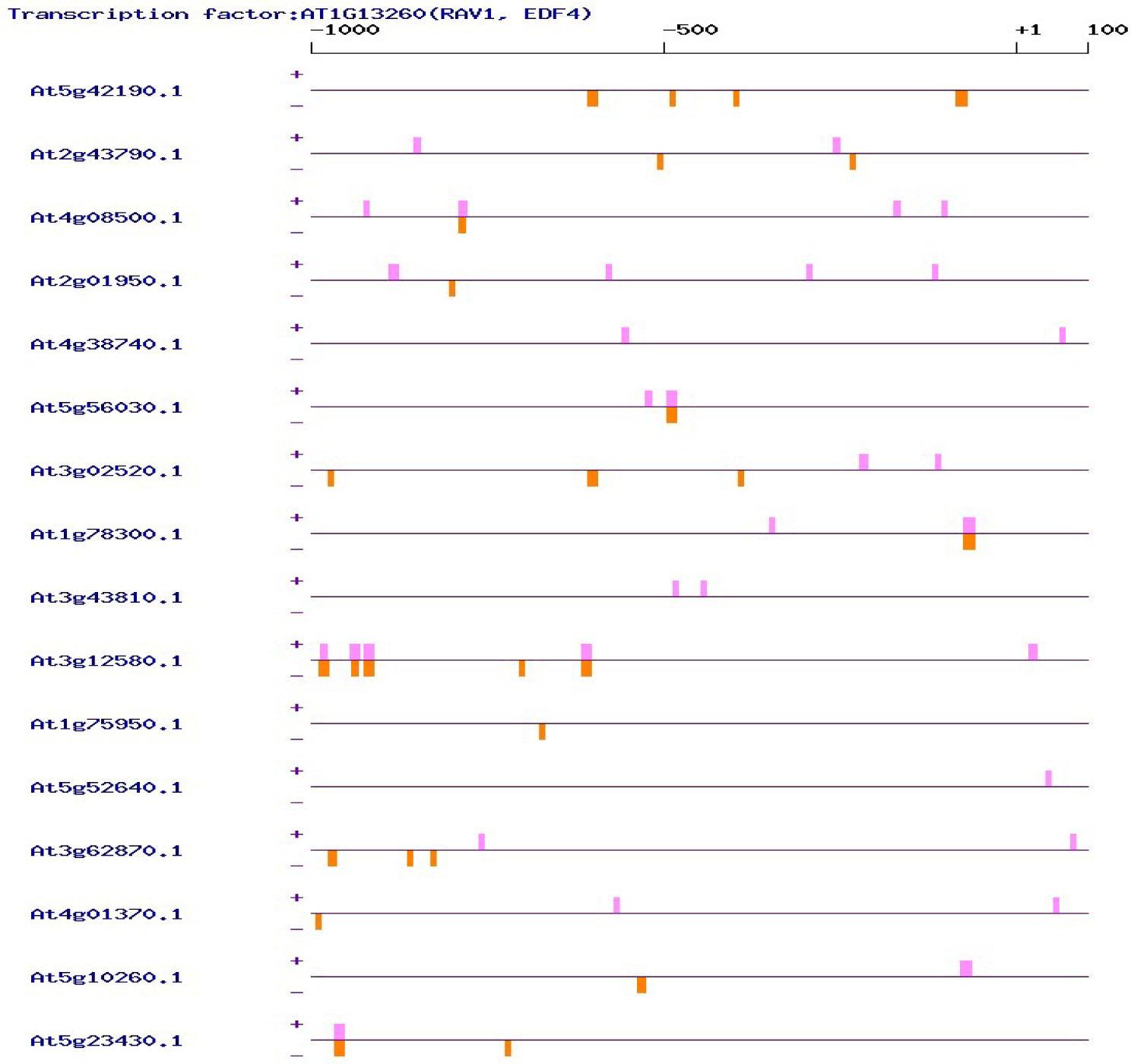
The AtRAV1 transcription factor (AT1G13260) binding sites in the promoter region of key defense genes. The AtRAV1 transcription factor (AT1G13260) binding sites in the promoter region in each of the 16 key defense genes are represented as vertical bars. The violet colour indicates binding sites in the negative strand while pink colour indicates binding sites in the positive strand.

**Fig. S3.**
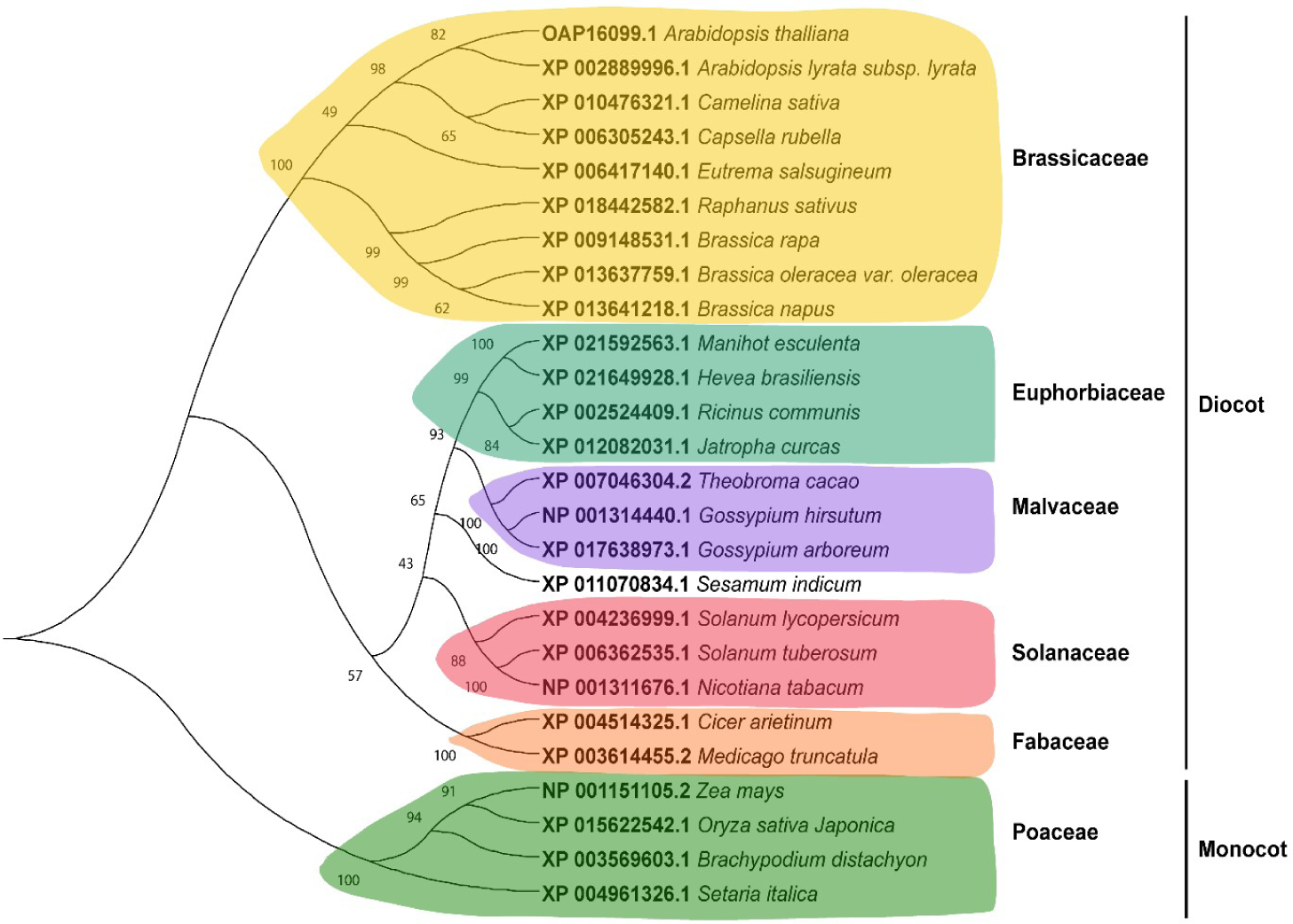
Phylogenetic analysis of RAV1 proteins. The bootstrap values are indicated at each branch node. The evolutionary distances were computed using the Poisson correction method and are in the units of the number of amino acid substitutions per site. The evolutionary analysis was conducted in MEGA X using Neighbor-Joining method.

**Fig. S4.**
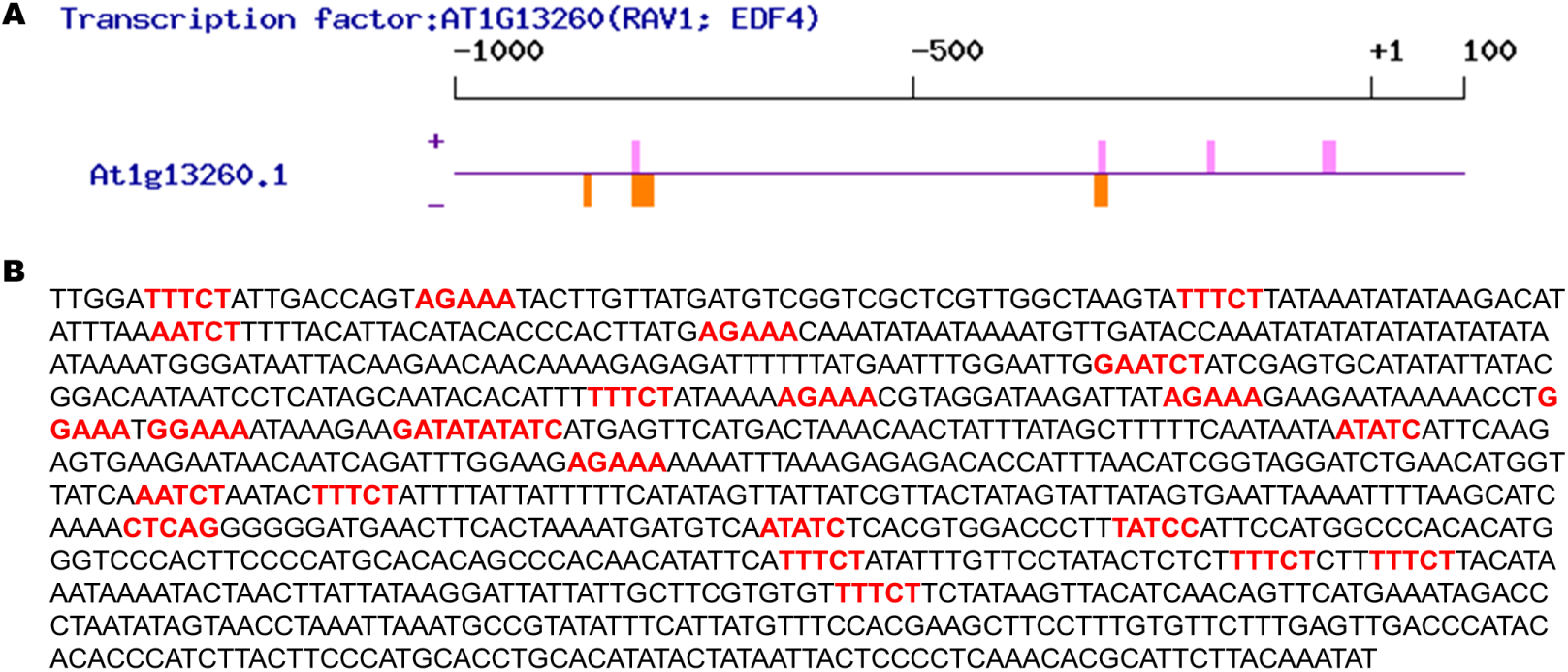
Potential RAV1 binding sites present in the promoter of *AtRAV1.* (A) Schematic view depicting the presence of potential RAV1 transcription factor binding sites in the *AtRAV1* promoter. The violet colour indicates binding sites in negative strand while pink colour indicates binding sites in the positive strand. (B) The potential RAV1 binding motifs are highlighted in the *AtRAV1* promoter region. Eight distinct RAV1 binding motifs [AGAAA (5), TTTCT (8), AATCT (3), GGAAA (2), GATAT (1), ATATC (3), CTCAG (1) and TATCC (1)], are highlighted.

**Fig. S5.**
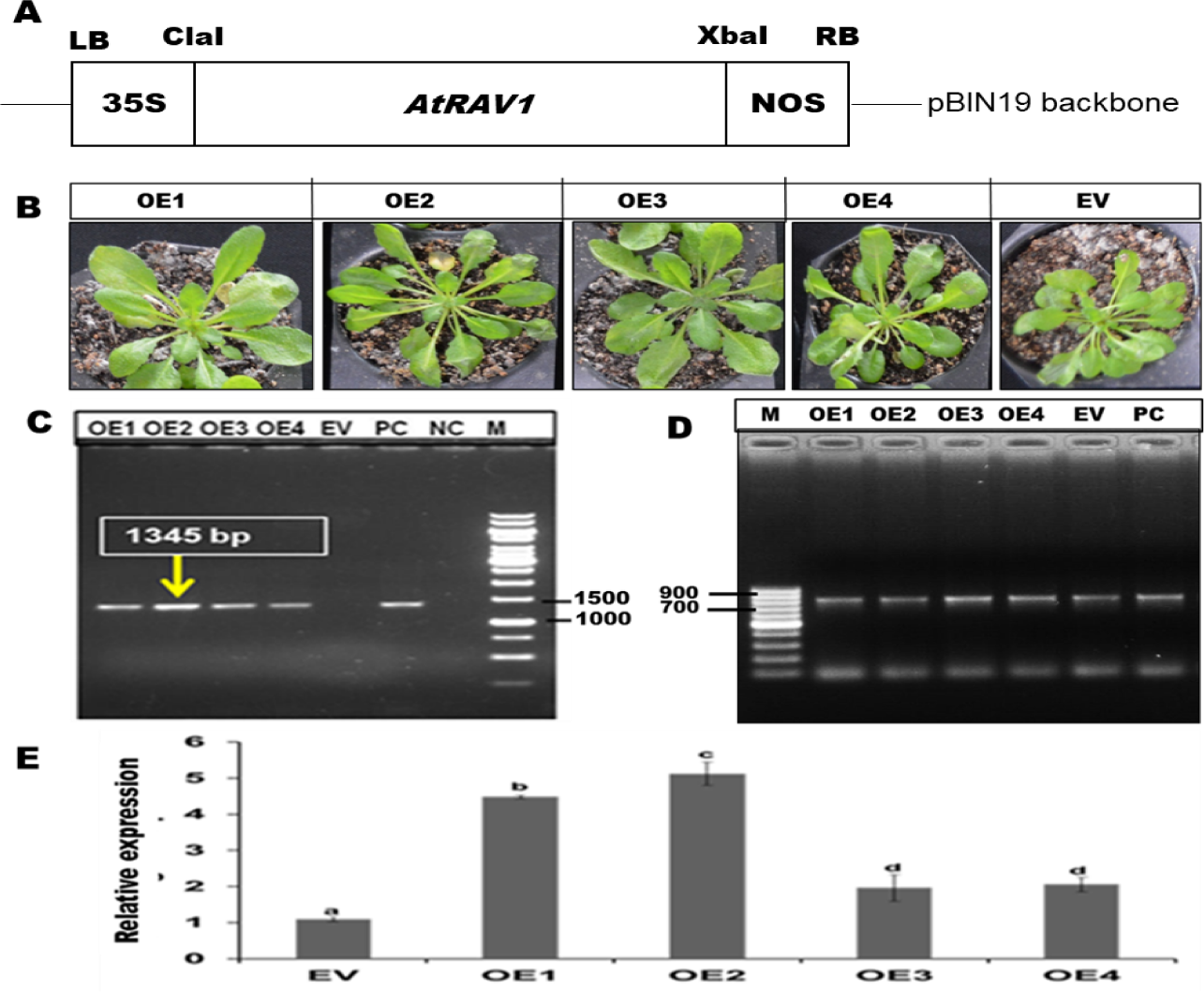
Characterization of transgenic *A. thaliana* lines. (A) The pGJ100 binary vector map that was used for generating transgenic lines. (B) The representative photographs of *AtRAV1* overexpressing (OEs) and empty vector (EV) transgenic plants. (C) PCR product using CaMV 35S F and RAV1OX-R primer pair. PC (positive control) represents PCR product obtained using plasmid of pGJ100 containing 35S:AtRAV1; NC (negative control) reflects no template control and M: represents DNA marker. (D) PCR product using NptII-F and NptII-R primers, highlighting the integration of T-DNA in both OE and EV transgenic lines. (E) Relative gene expression of *AtRAV1* in different OE and EV lines, when normalized with the expression in wild type (WT) plants using beta-actin as housekeeping gene. Graph shows mean values ± standard error of at least three biological replicates. Values with different letters are significantly different at *P* < *0.05* (estimated using one-way ANOVA)

**Fig. S6.**
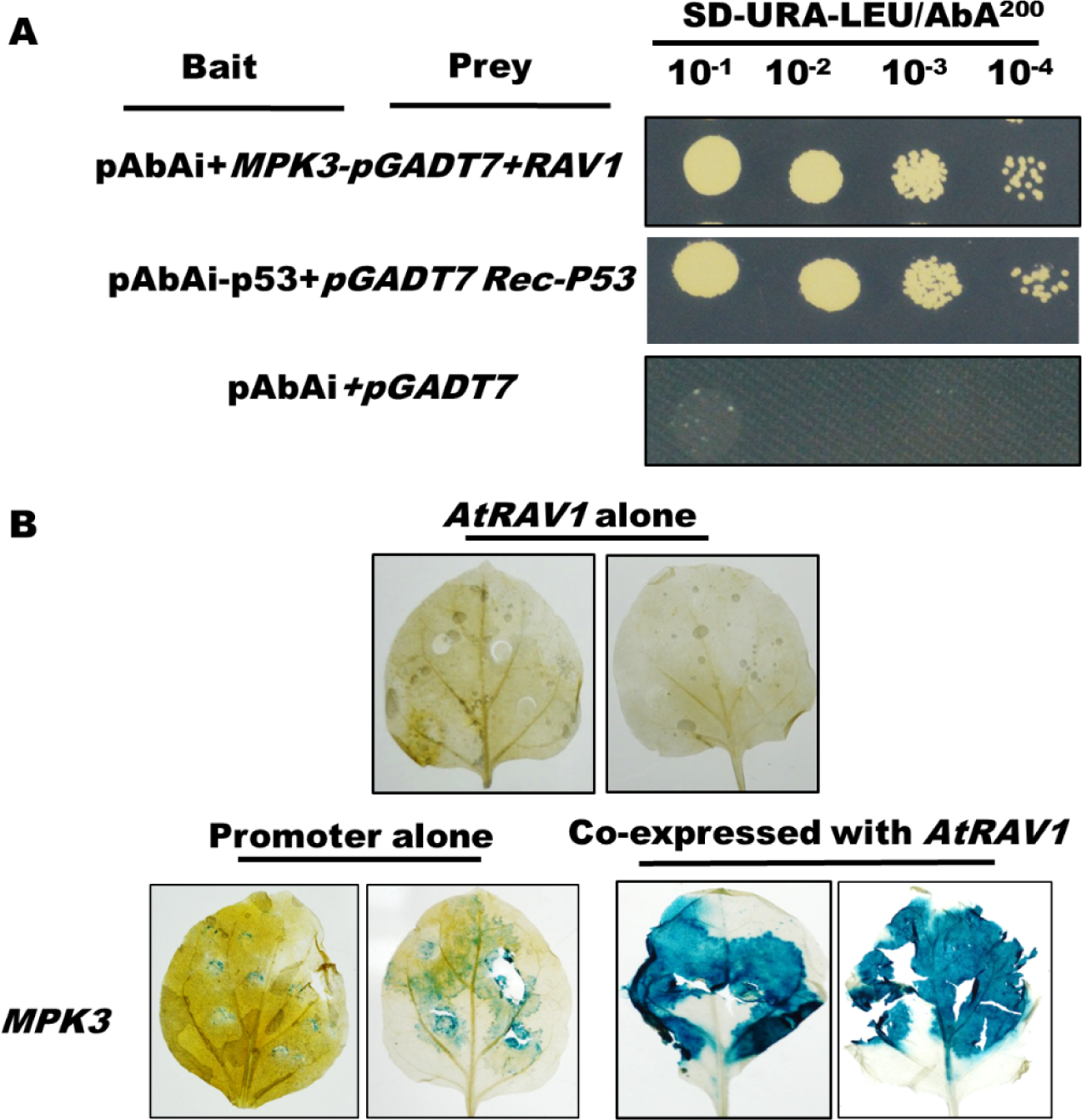
*AtRAV1* induces the expression of AtMPK3 by potentially binding to its promoter. (A) Yeast one hybrid assay: growth of yeast cells on SD-URA-LEU medium in presence of AbA (Aureobasidin A) reflects that AtRAV1 transactivates the expression of AbA gene under *AtMPK3* promoter. The pAbAi-p53+pGADT7Rec-P53 was used as a positive control while the empty vector of pAbAi + pGADT7 was used as negative control. (B) GUS based reporter assay suggesting that AtRAV1 activates GUS expression driven through the promoter of *AtMPK3* gene in *N. benthamiana*.

**Fig. S7.**
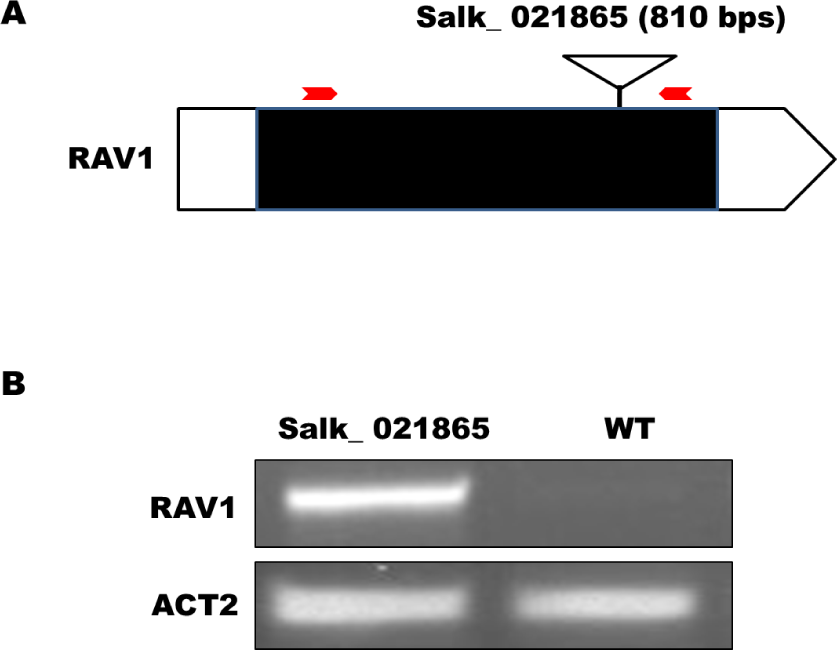
Validation of *atrav1* mutant line (Salk_021865). (A) The position of T-DNA insertion (marked by inverted triangles) in *AtRAV1* gene in. Red arrow indicates the positions of primers used to validate T-DNA insertion. (B) PCR product showing T-DNA insertion in *atrav1* mutant. *ACT2* (beta actin) gene of *A. thaliana* was used as loading control.

**Fig. S8.**
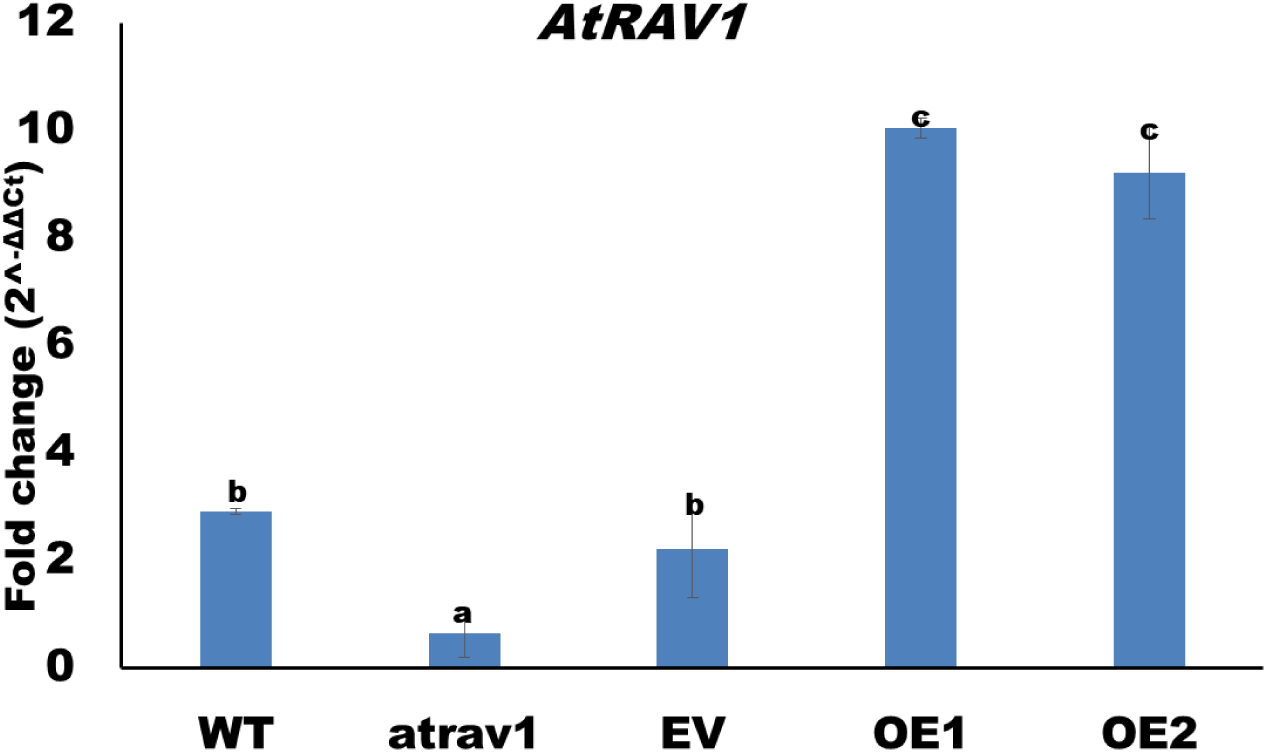
The pathogen infection induces the expression of *AtRAV1*. The relative expression of *AtRAV1* gene in *R. solani* infected *A. thaliana* plants are summarized as bar chart. The relative expression was quantified by normalizing the expression with the uninfected samples using beta actin as endogenous control. Graph shows mean values ± standard error of at least three technical replicates. Values with different letters are significantly different at *P* < *0.05* (estimated using one-way ANOVA). Similar results were obtained in three biological repeats.

**Fig. S9.**
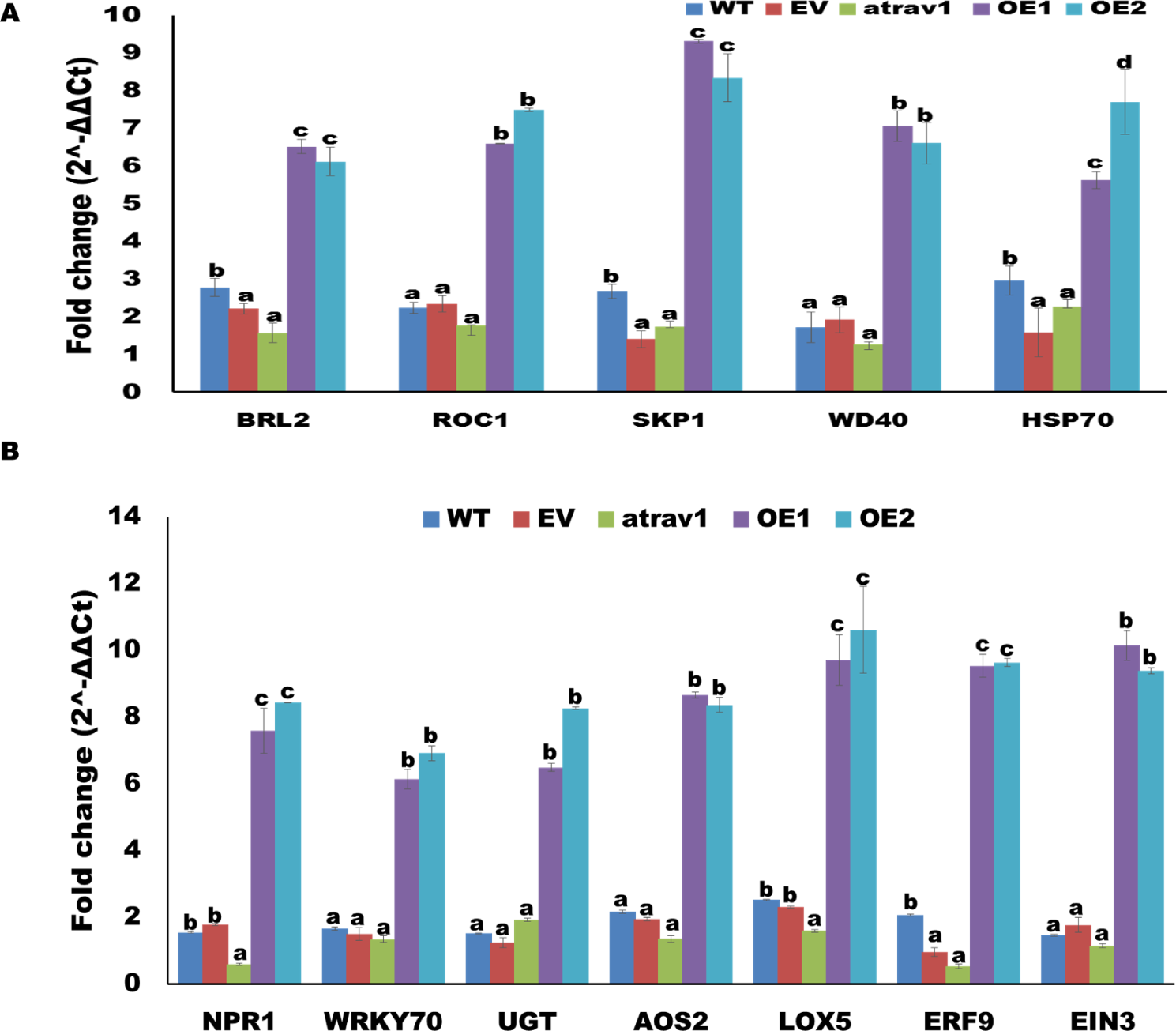
*R. solani* infection enhances the expression of key defense and defense marker genes in *AtRAV1* overexpression lines. The expression of (A) selected key defense genes and (B) Defense marker genes in *R. solani* infected lines. The relative expression was quantified by normalizing the expression with uninfected samples using beta actin as endogenous control. Graph shows mean values ± standard error of at least three technical replicates. For each gene, different letters indicate significant difference at *P*<*0.05* (estimated using one-way ANOVA). Similar results were obtained in three biological repeats.

**Fig. S10.**
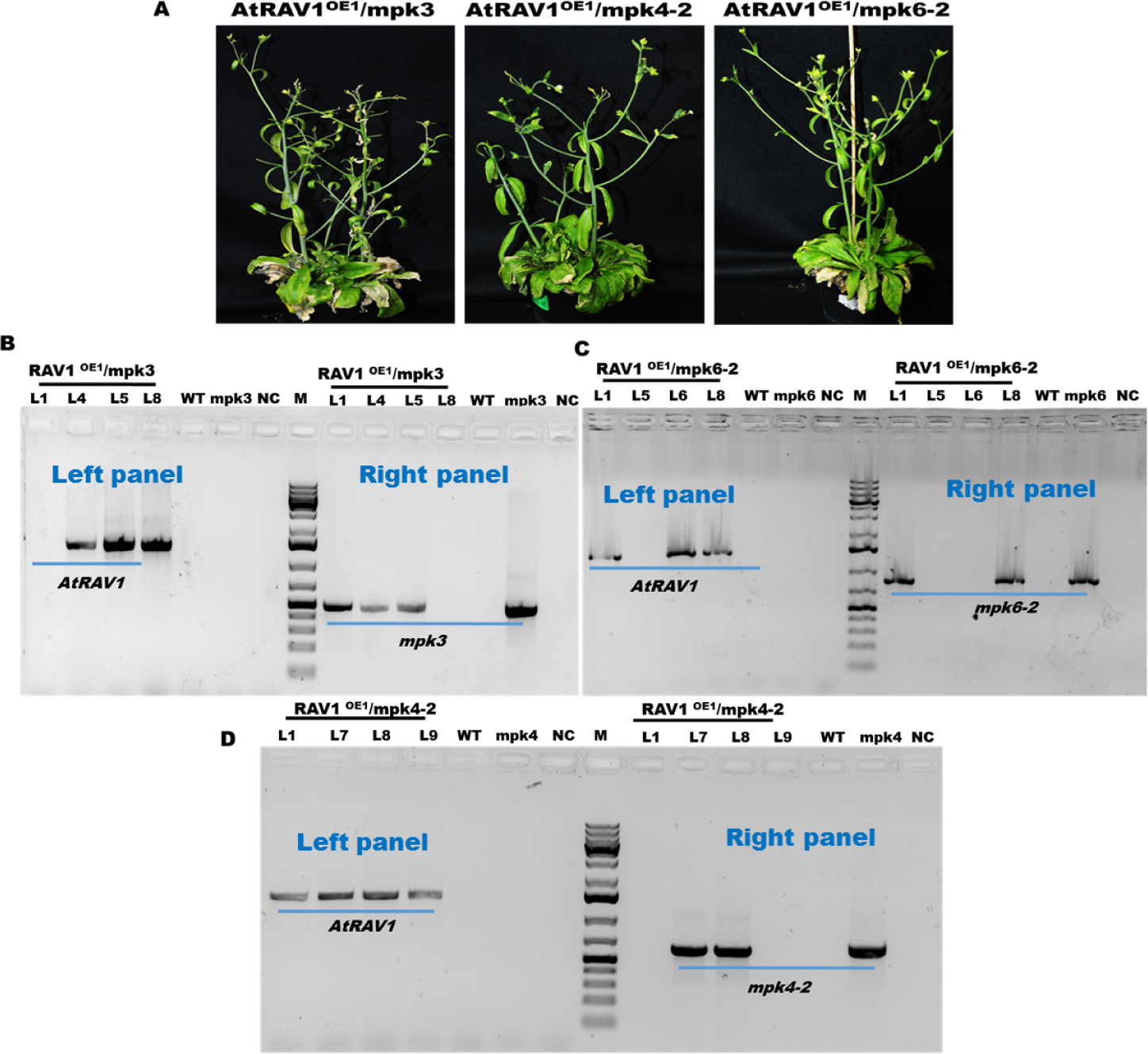
Validation of AtRAV1^OE1^/mpk3, AtRAV1^OE1^/mpk4-2 and AtRAV1^OE1^/mpk6-2 lines in *A. thaliana.* (A) The representative images of different plants (AtRAV1^OE1^/mpk3, AtRAV1^OE1^/mpk4-2 and AtRAV1^OE1^/mpk6-2). B, C and D right panel represent PCR based validation of T-DNA insertion of different MAP kinase knockout mutant (mpk3, mpk4-2 and mpk6-2) in *AtRAV1* OE1 lines using T-DNA border primer and gene specific reverse primer (BP+RP). B, C and D left panel represent presence of *AtRAV1* transgene was confirmed by PCR using CaMV 35S F and RAV1OX-R primer. Negative control (NC) reflects no template control.

**Fig. S11.**
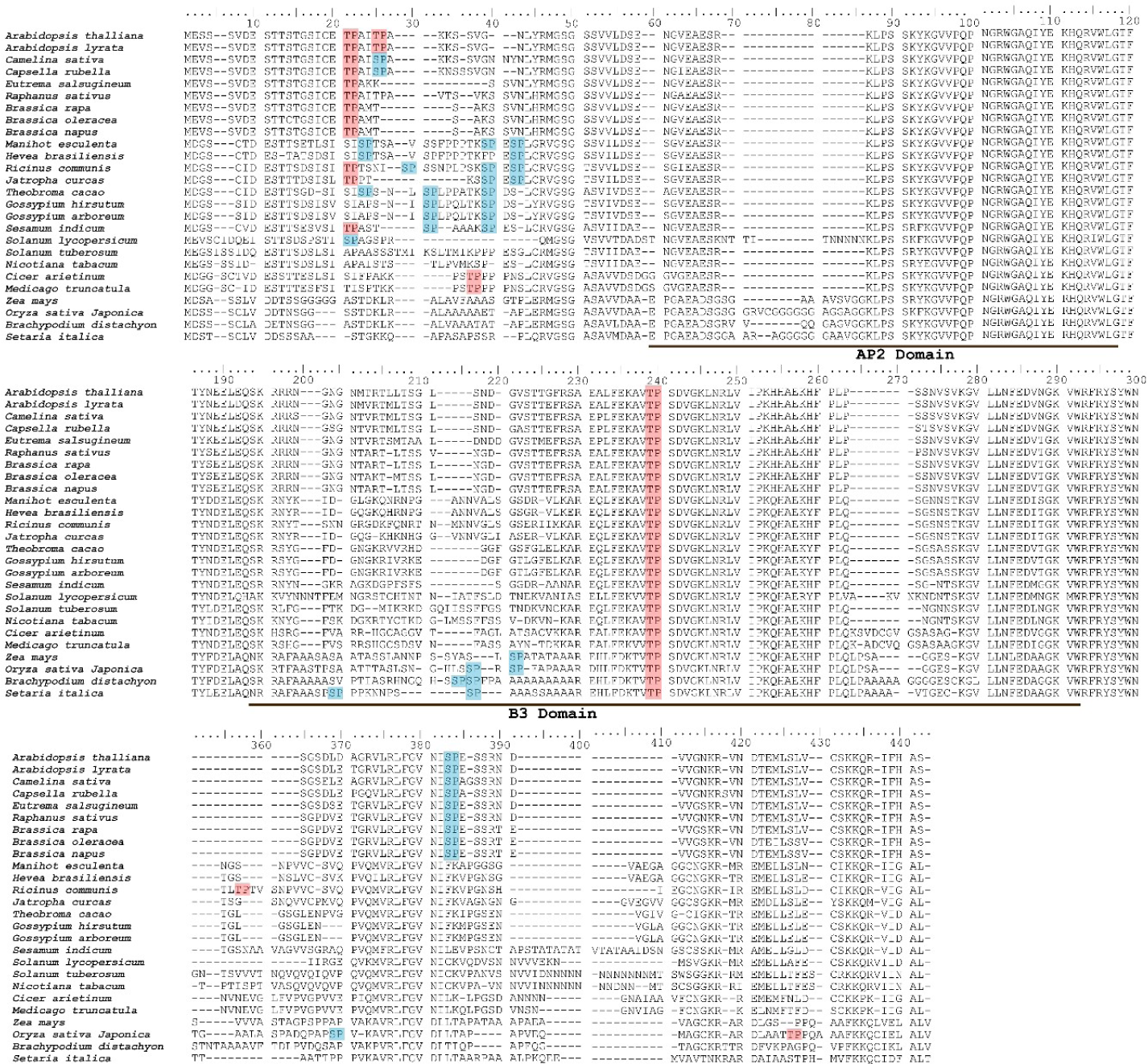
ClustalW alignment of RAV1 protein sequences in different plants. Presence of conserved AP2 and B3 domain region and potential MAP kinase phosphorylation sites (TP in red and SP in blue) in RAV1 amino acid sequence is highlighted.

**Fig. S12.**
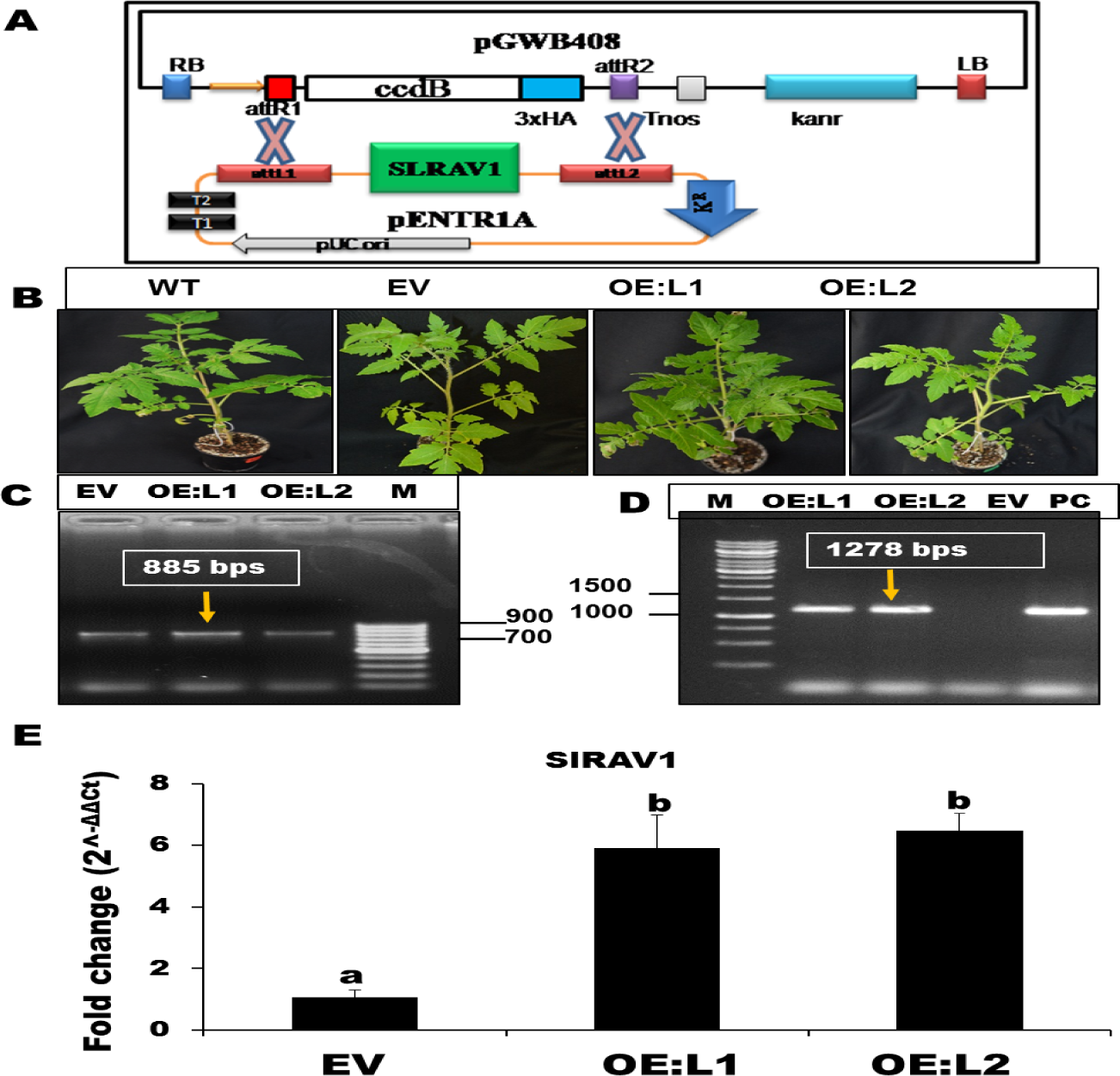
Characterization of transgenic tomato lines. (A) T-DNA map of pGWB408 gateway binary vector that was used for generating transgenic lines in tomato. (B) The representative images of empty vector (EV) and *SlRAV1* overexpressing (OEs) transgenic plants. (C) Presence of T-DNA (885 bp) confirmed by PCR using NptII-F and NptII-R primer. (D) Presence of *SlRAV1* transgene (1278bp) confirmed by PCR using CaMV 35S F and SlRAV1OE-R primer. Positive control (PC) represents PCR product obtained from recombinant plasmid (pGWB408 containing 35S:*SlRAV1*). (E) Relative gene expression of *SlRAV1* in different OE and EV lines, upon normalization with respect to WT plants. Graph shows mean values ± standard error of at least three technical replicates. Values with different letters are significantly different at *P* < *0.05* (estimated using one-way ANOVA). Similar results were obtained in three biological repeats.

**Fig. S13.**
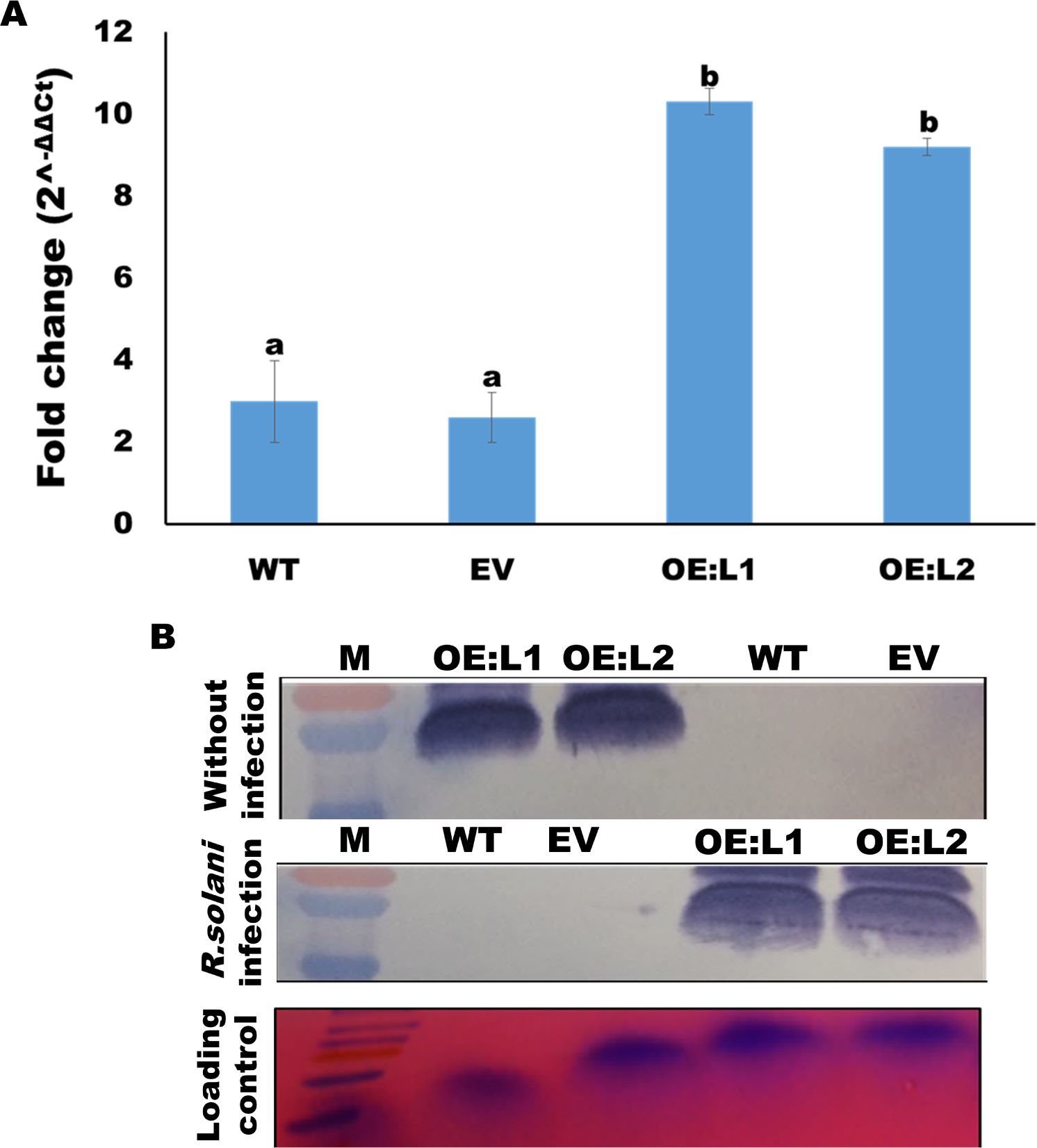
*R. solani* infection upregulates the expression of SlRAV1 in tomato. (A) The relative expression of *SlRAV1* gene in *R. solani* infected (4 dpi) tomato leaves are summarized as bar chart. The relative expression was quantified by normalizing the expression with uninfected samples using beta actin as endogenous control. Graph shows mean values ± standard error of at least three technical replicates. Values with different letters are significantly different at *P* < *0.05* (estimated using one-way ANOVA). (B) Western-blot analysis reflecting the accumulation of His-tagged version of SlRAV1 protein in different tomato plants with or without *R. solani* infection. Similar results were obtained in three biological repeats.

**Fig. S14.**
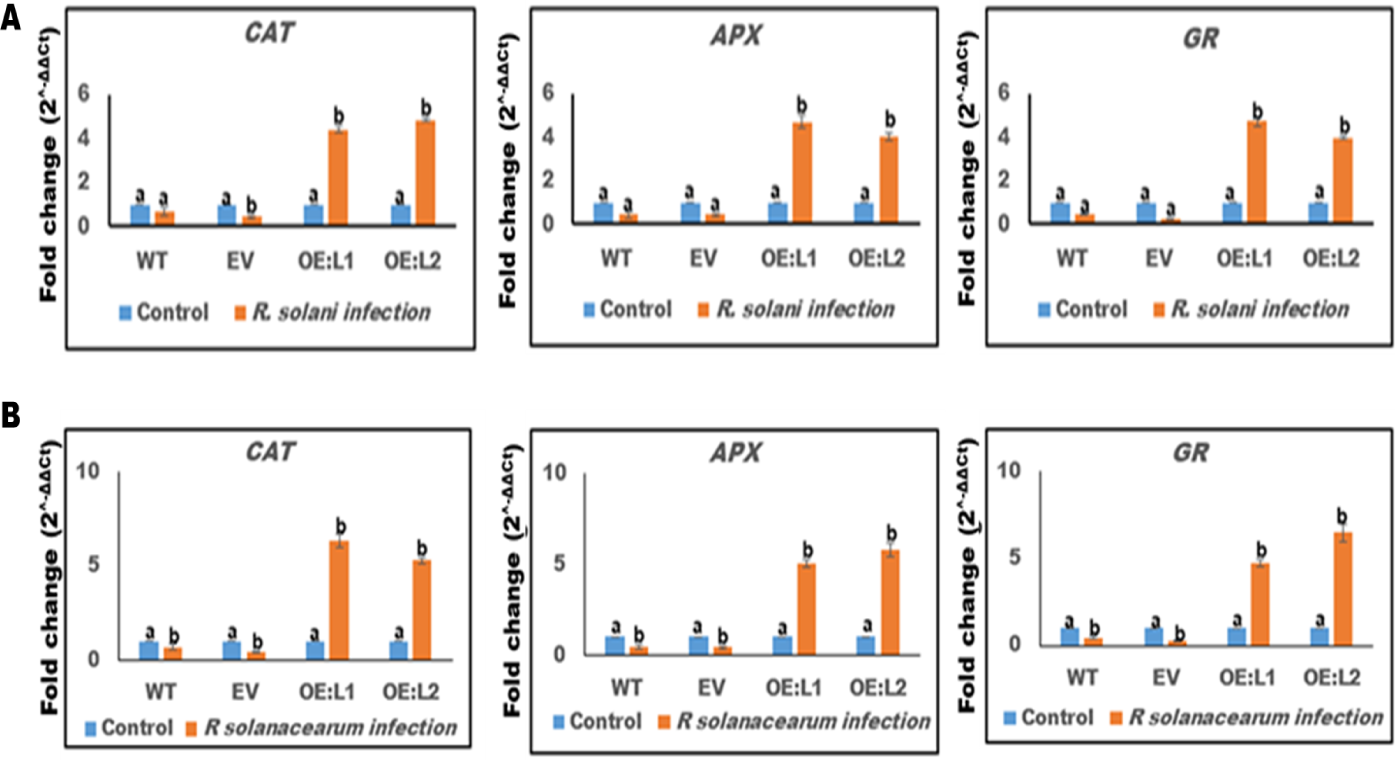
Expression analysis of different antioxidant marker genes upon *R. solani* and *R. solanacearum* infection. The relative expression of various antioxidant marker genes (*CAT, APX* and *GR*) in (A) *R. solani* (4 dpi) and (B) *R. solanacearum* (7 dpi) infected tomato leaves. The relative expression was quantified by normalizing the expression with uninfected samples using beta actin as endogenous control. Graph shows mean values ± standard error of at least three technical replicates. Values with different letters are significantly different at *P* < *0.05* (estimated using one-way ANOVA). Similar results were obtained in three biological repeats.

**Fig. S15.**
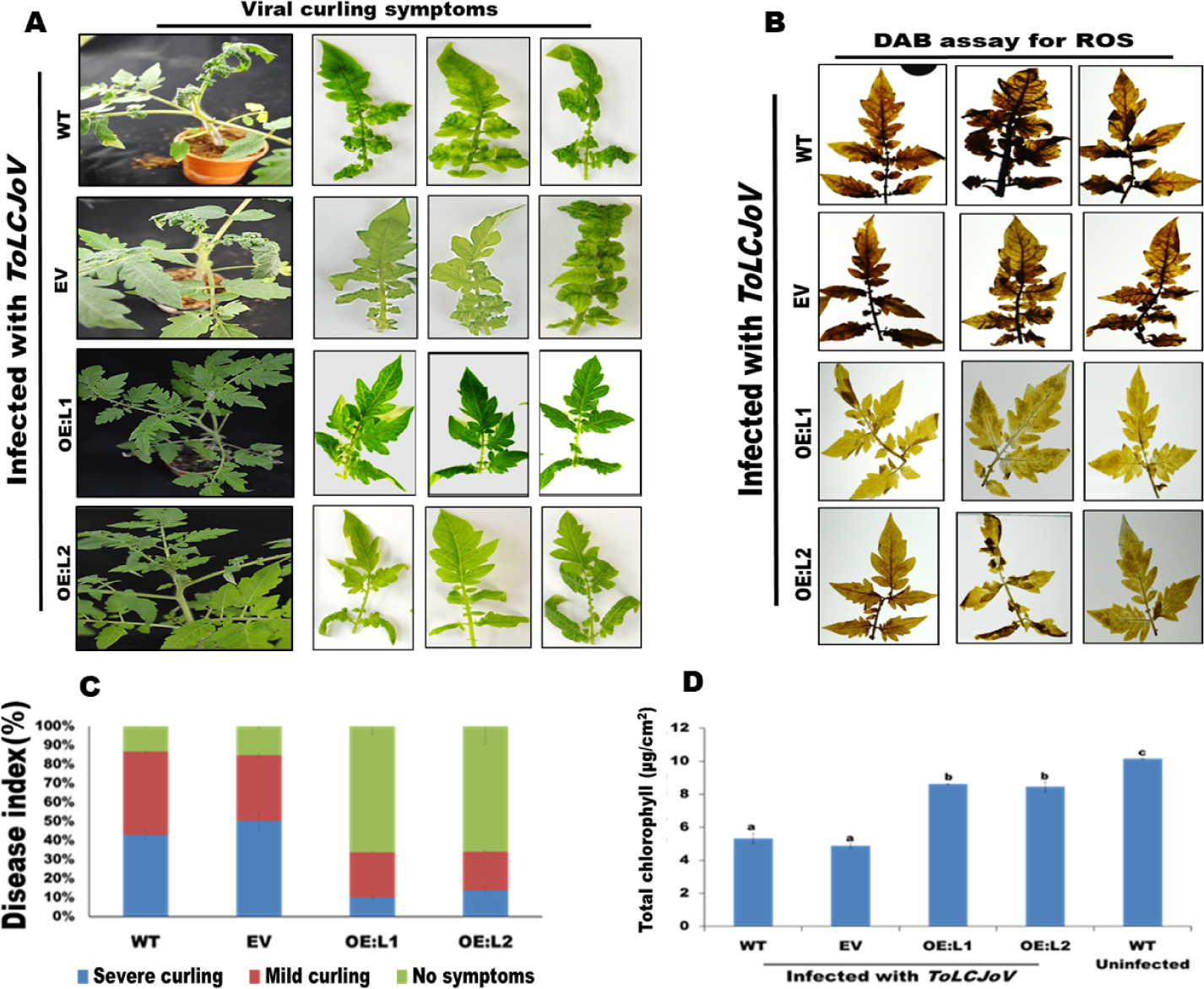
*SlRAV1* overexpression provides tolerance against *Tomato leaf curl Joydepur virus (ToLCJoV)* infections in tomato. (A) Disease symptoms (leaf curling) in *ToLCJoV* infected tomato at 21 dpi. (B) DAB staining of *ToLCJoV* infected tomato leaves. (C) Observed disease symptoms in *ToLCJoV* infected tomato leaves plotted as disease index. (D) Total chlorophyll content in *ToLCJoV* infected tomato leaves at 21 dpi. Graph shows mean values ± standard error of at least three technical replicates. Values with different letters are significantly different at *P* < *0.05* (estimated using one-way ANOVA). Similar results were obtained in three biological repeats.

**Table S1.** RAV1 binding motifs in the promoter region of selected key defence genes.

| Gene name | Promoter region having potential<br>AtRAV1 binding sites | Number of RAV1<br>binding motifs |
| --- | --- | --- |
| MPK4 | -124 to -1858 | 3 |
| ROC1 | -169 to -1812 | 11 |
| WD40 | -133 to -1775 | 6 |
| BRL2 | -40 to -1916 | 9 |
| SKP1 | -167 to -1906 | 6 |
| HSP70 | -103 to -1806 | 11 |

**Table S2.**
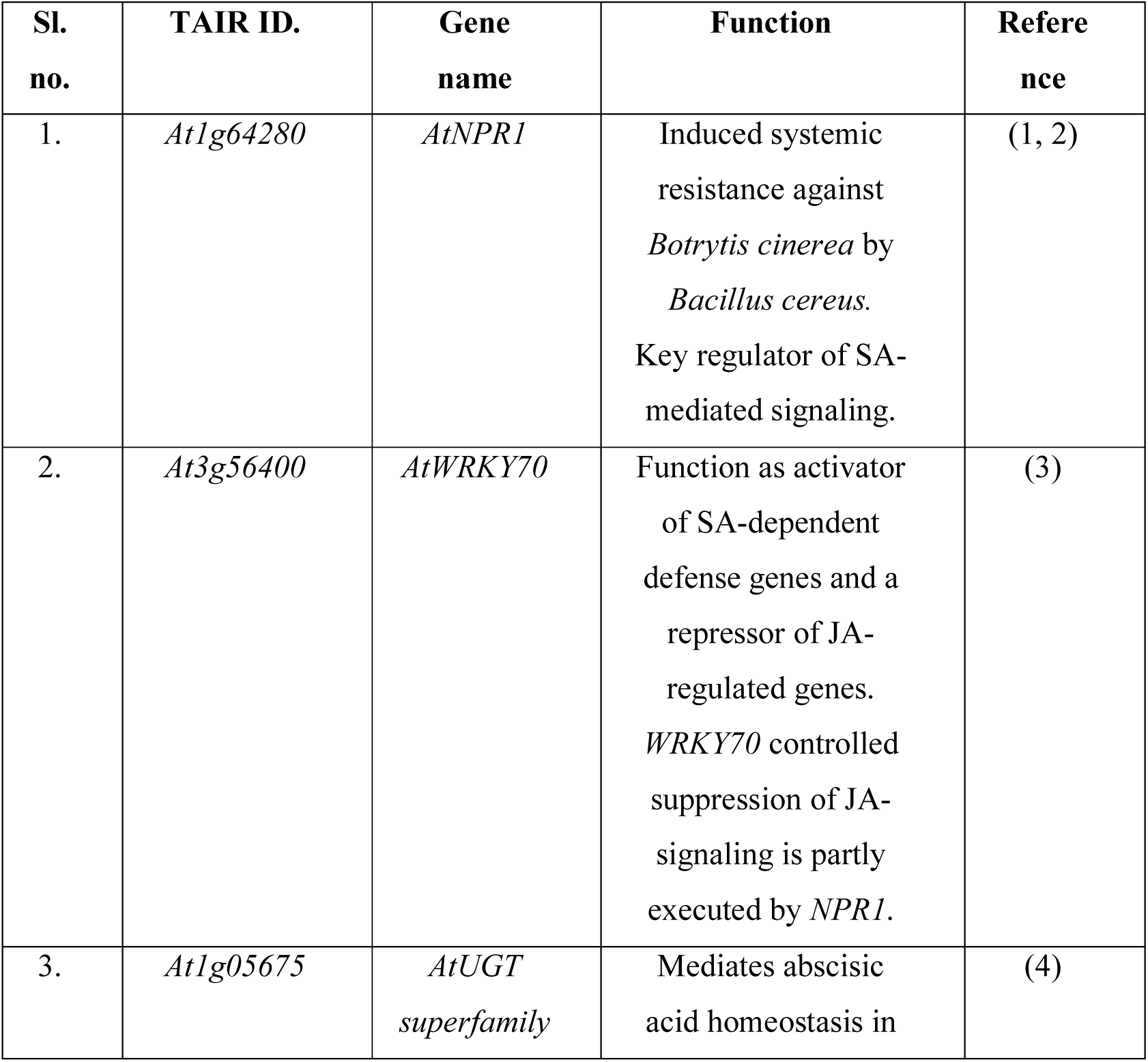
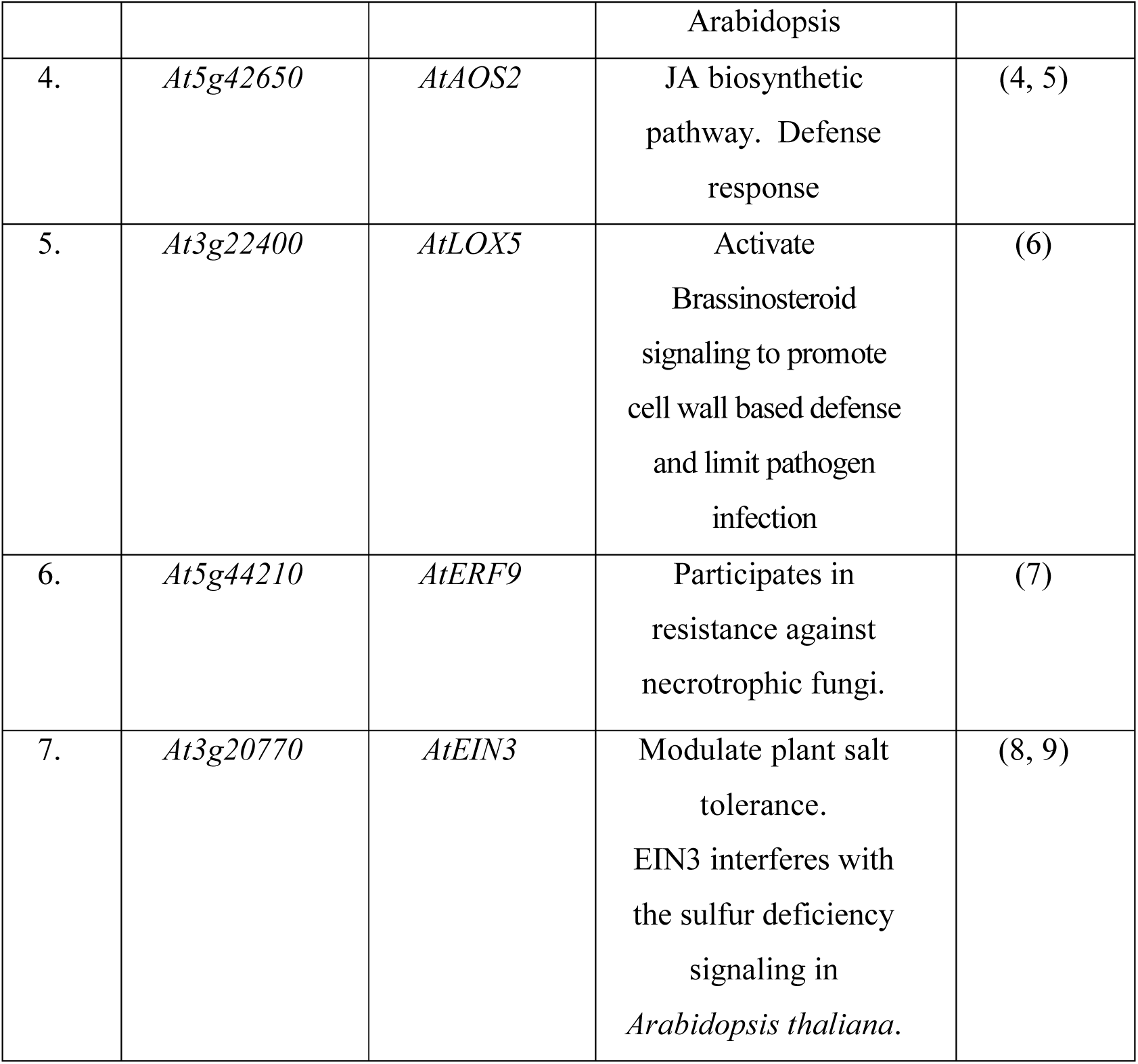
List of Salicylic acid, Jasmonic acid and Ethylene (SA, JA and ET) mediated defense marker gene used in this study.

**Table S3.** As XLS sheet (attached separately)

**Table S4.**
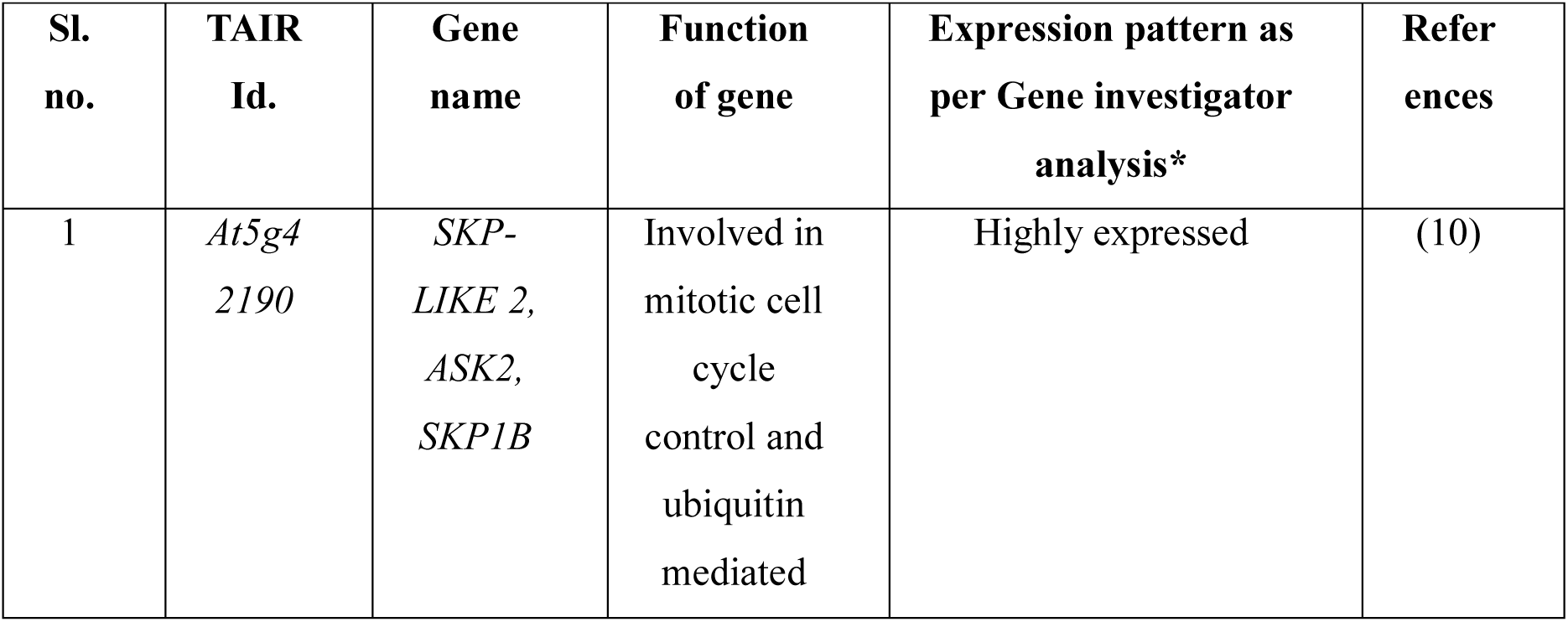
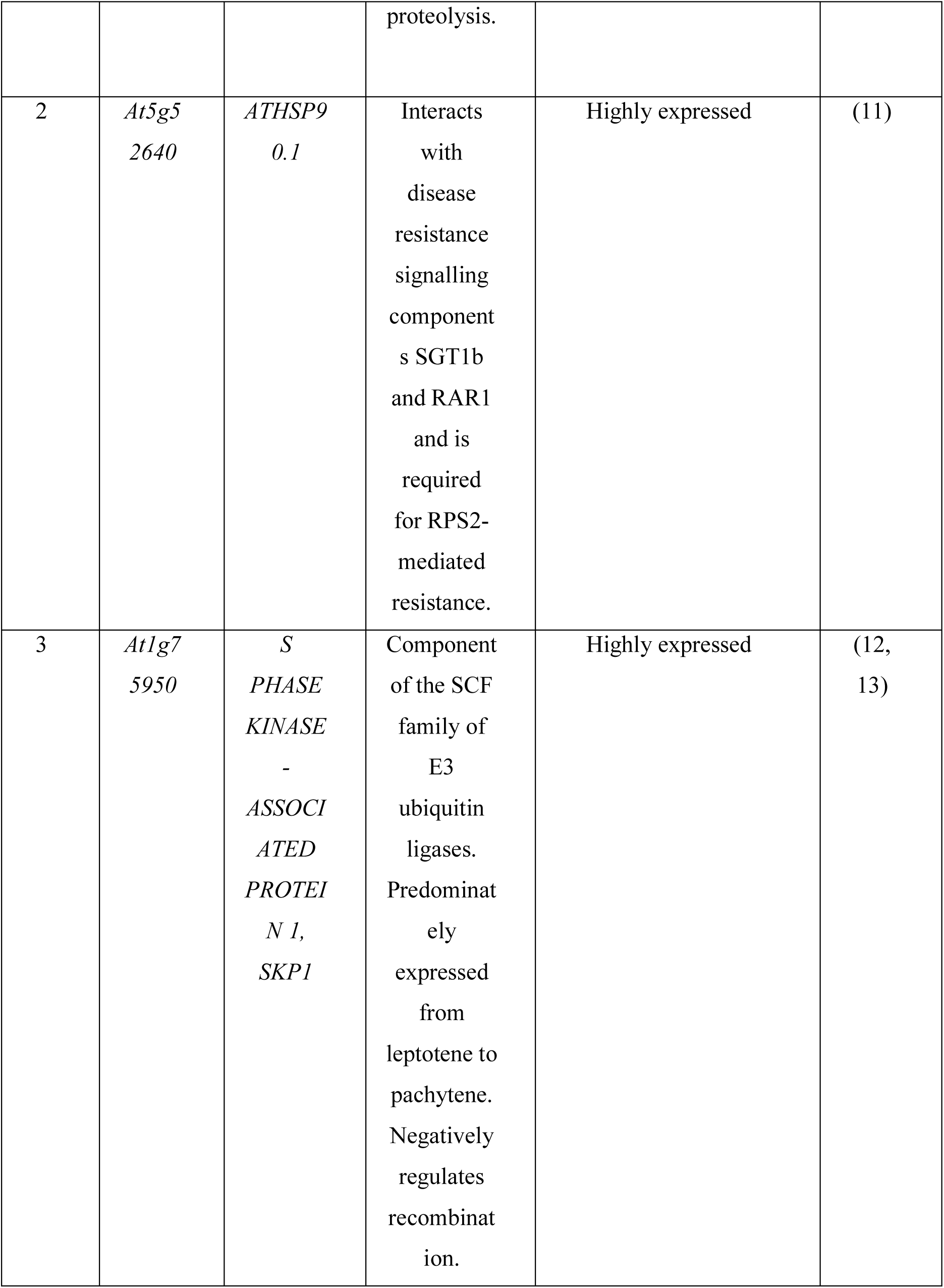
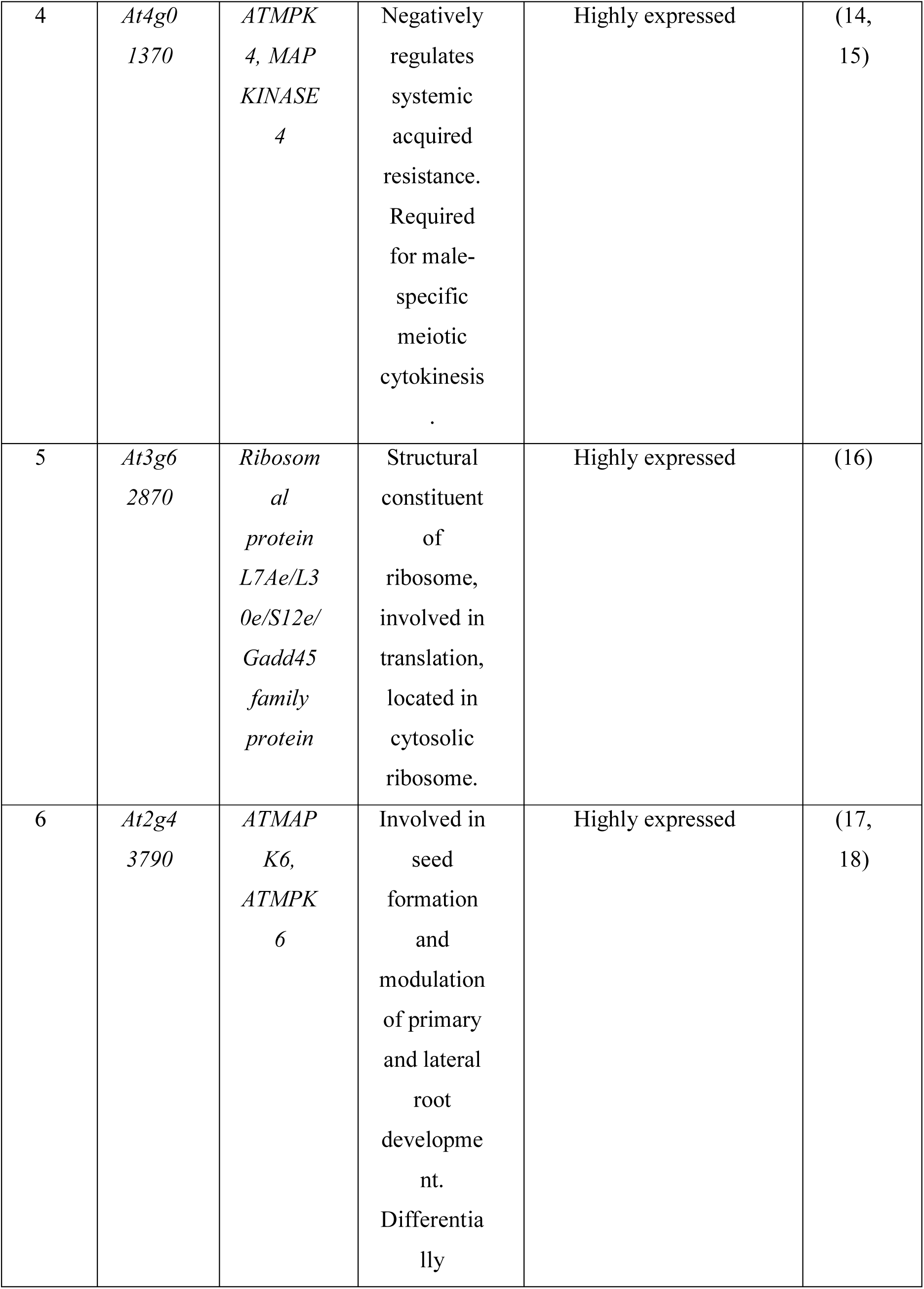
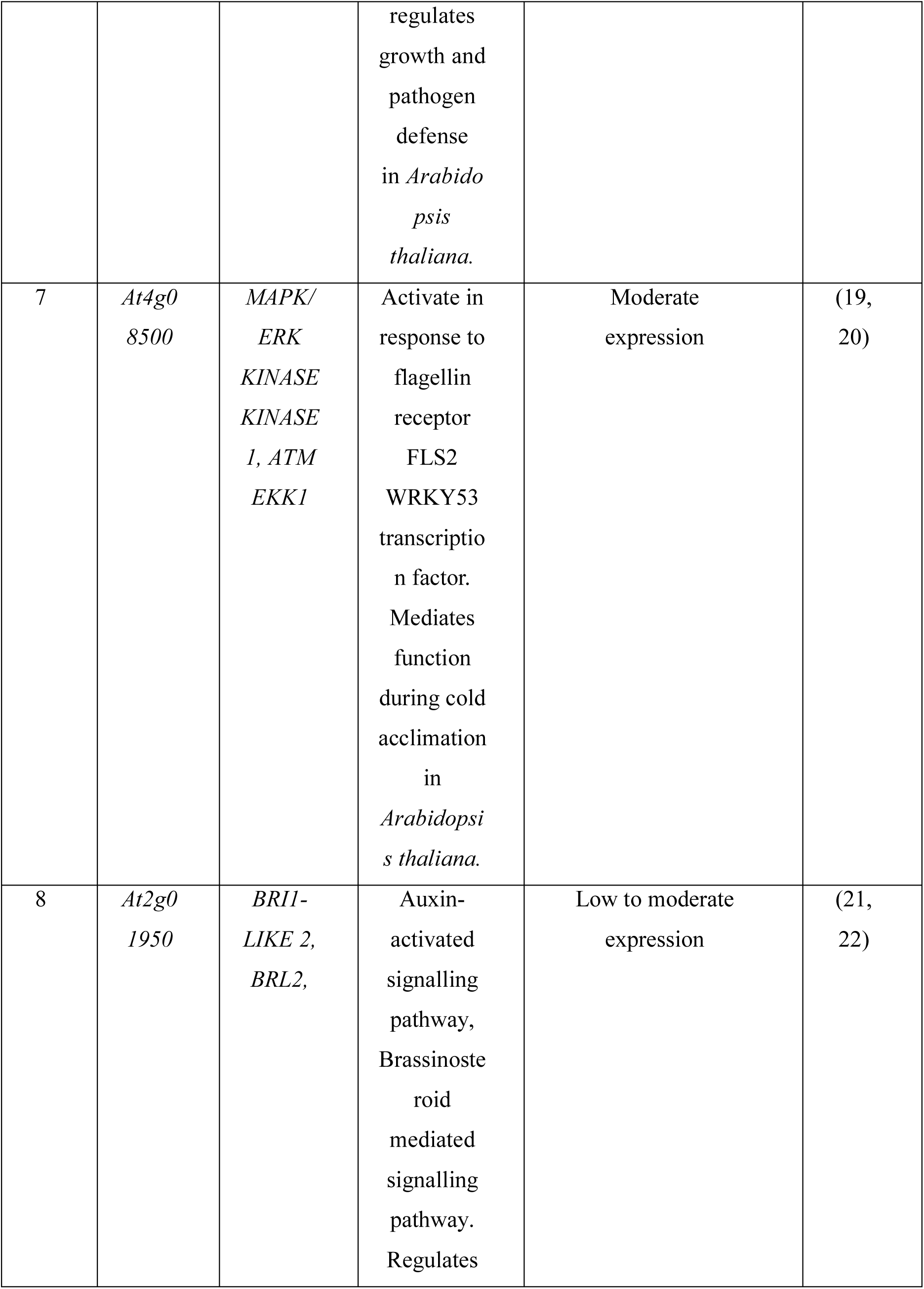
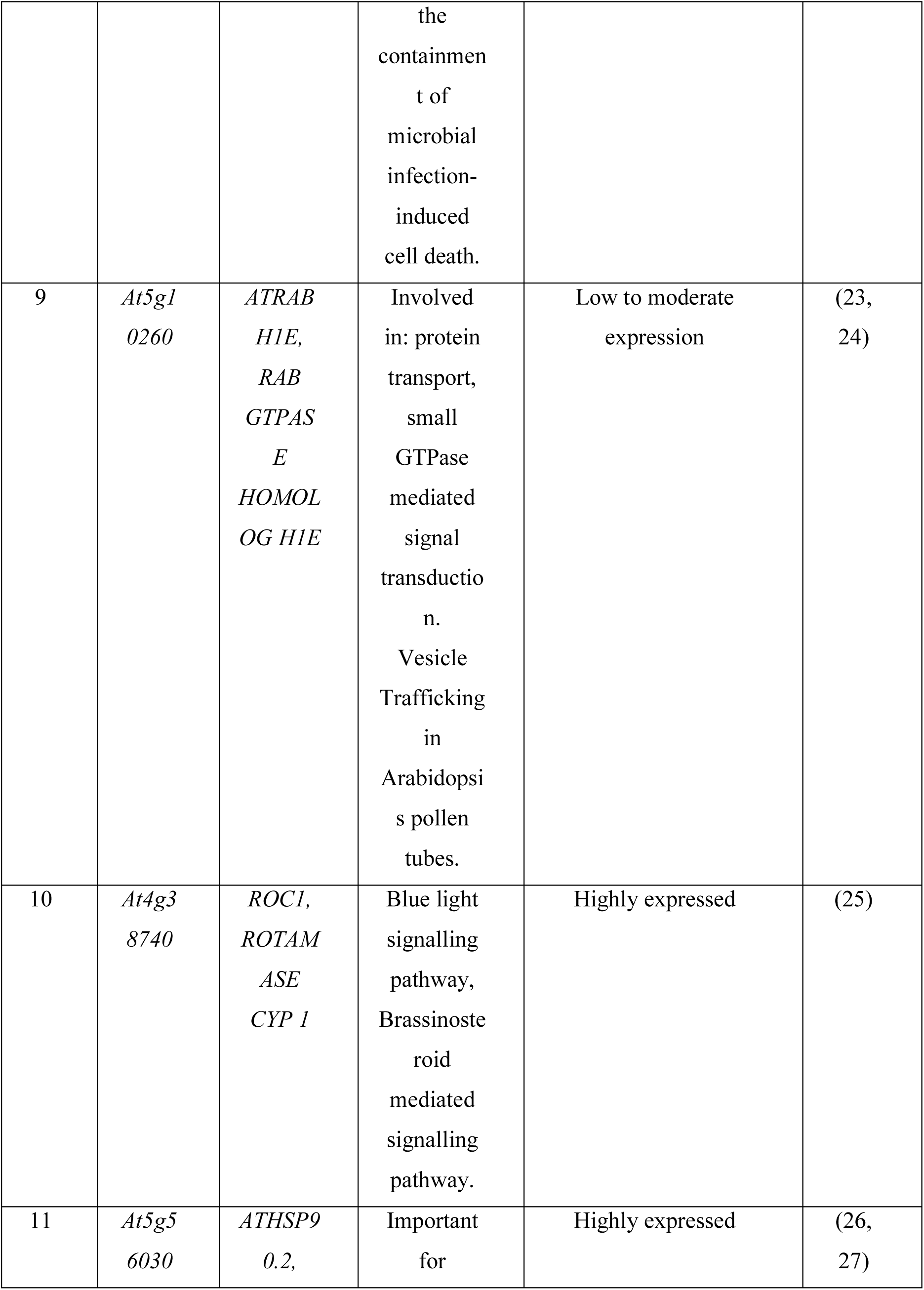
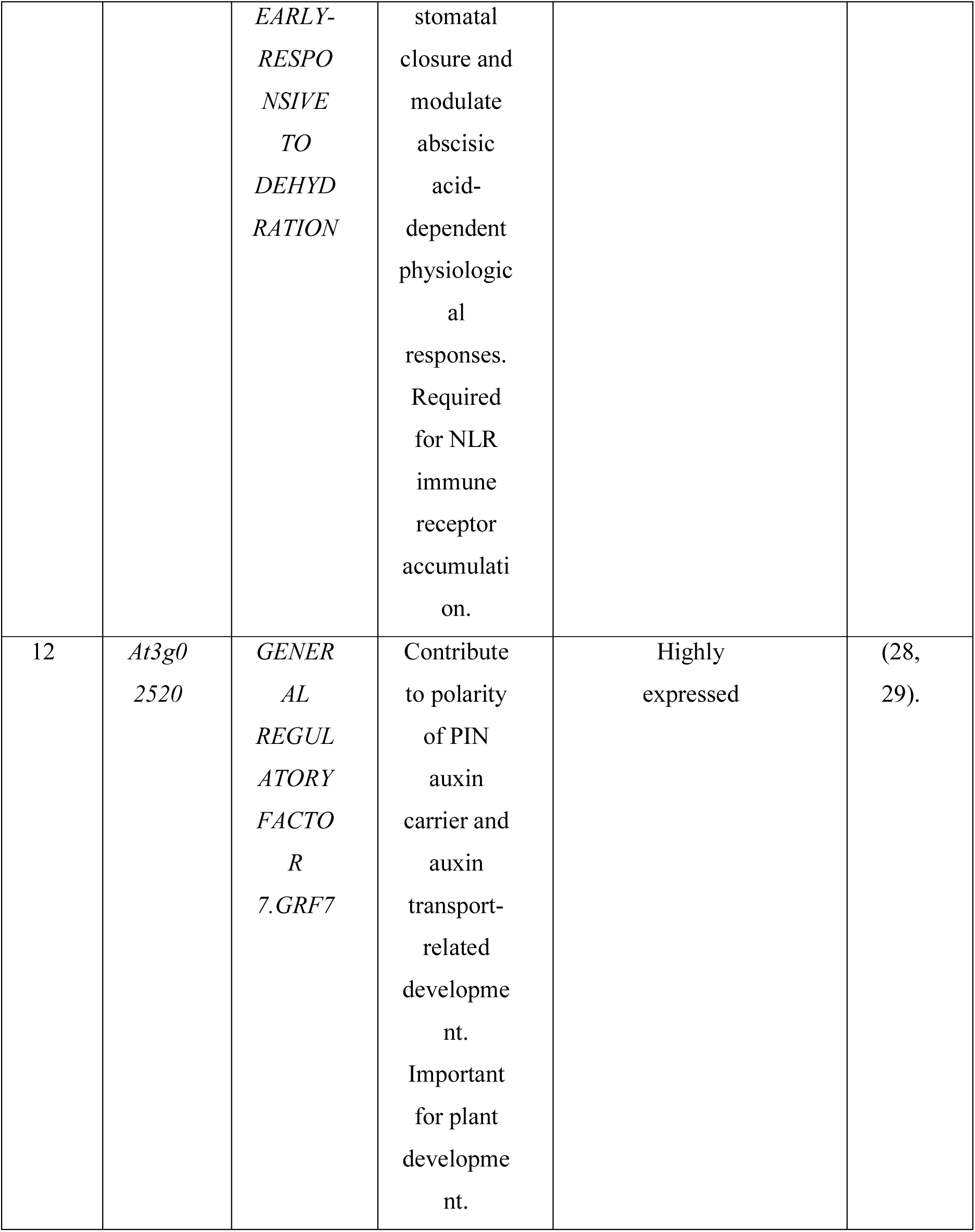
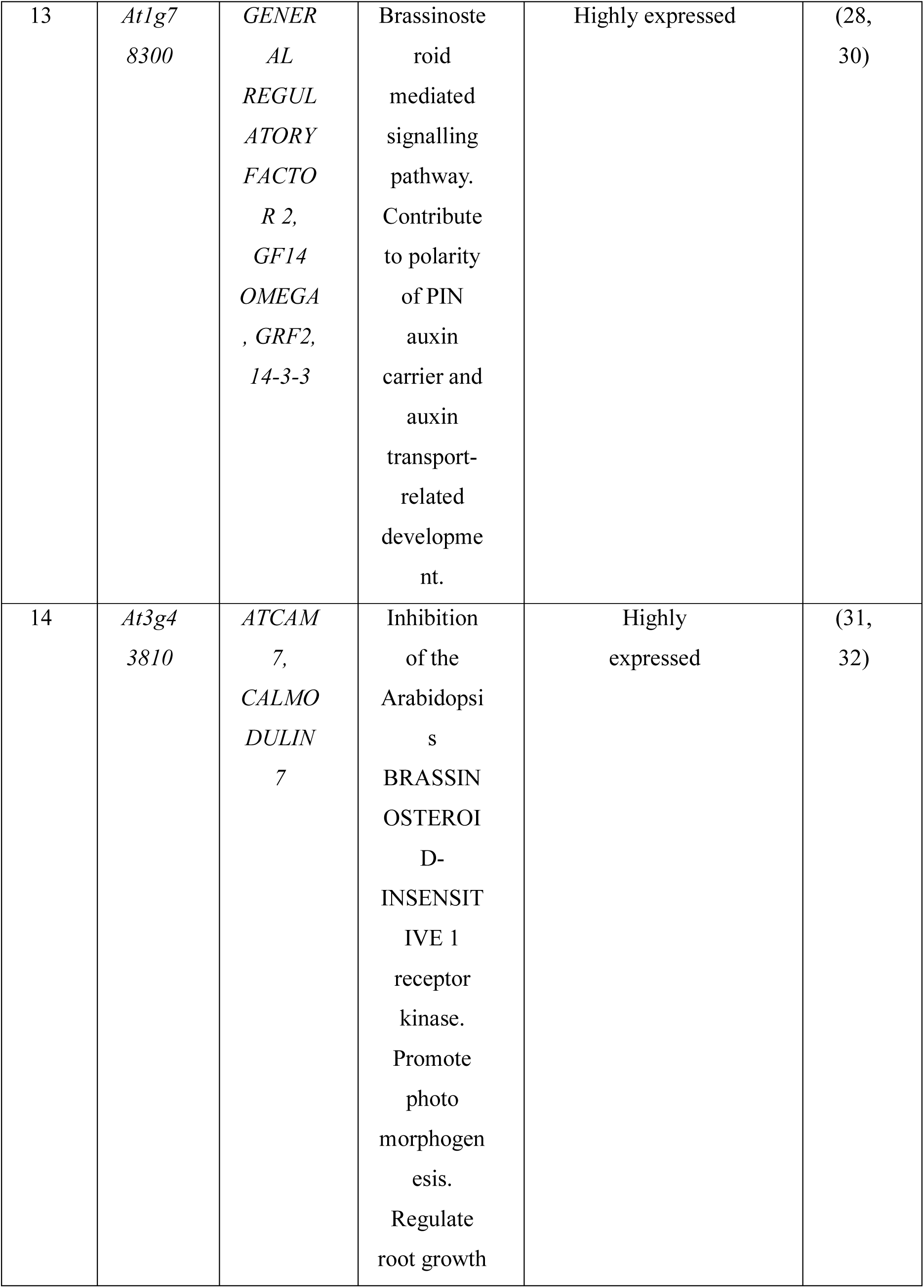
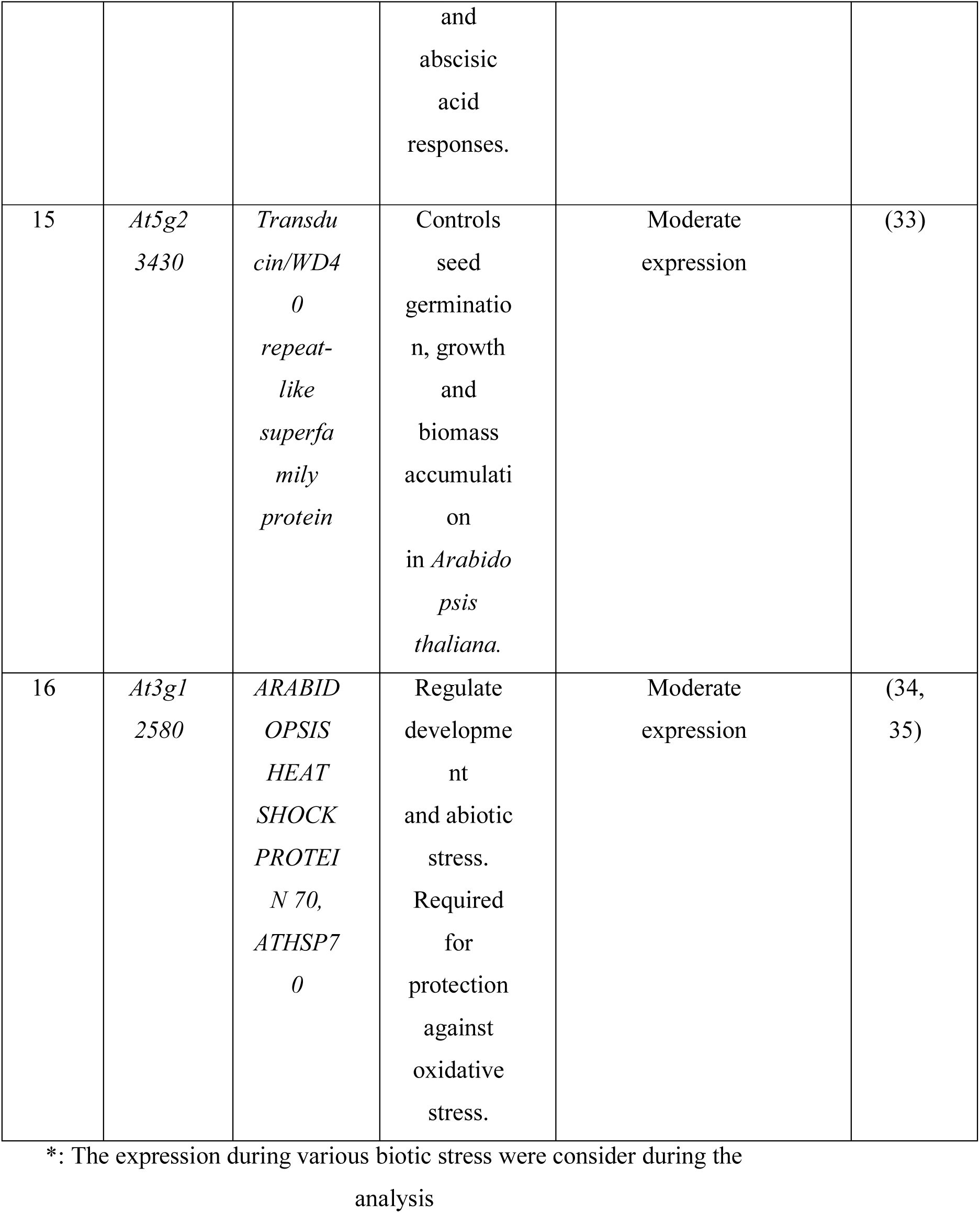
List of 16 key defense genes.

**Table S5.**
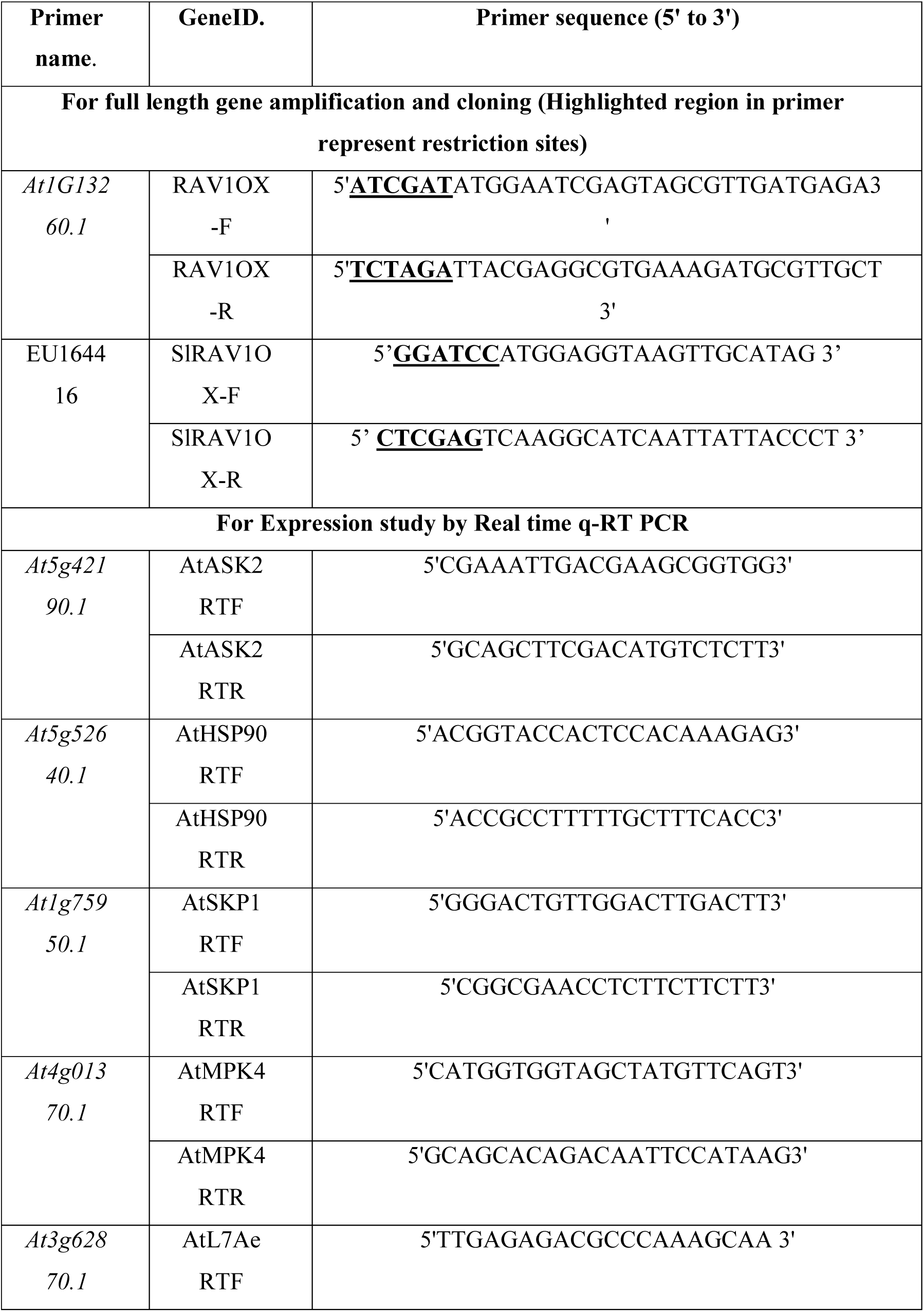
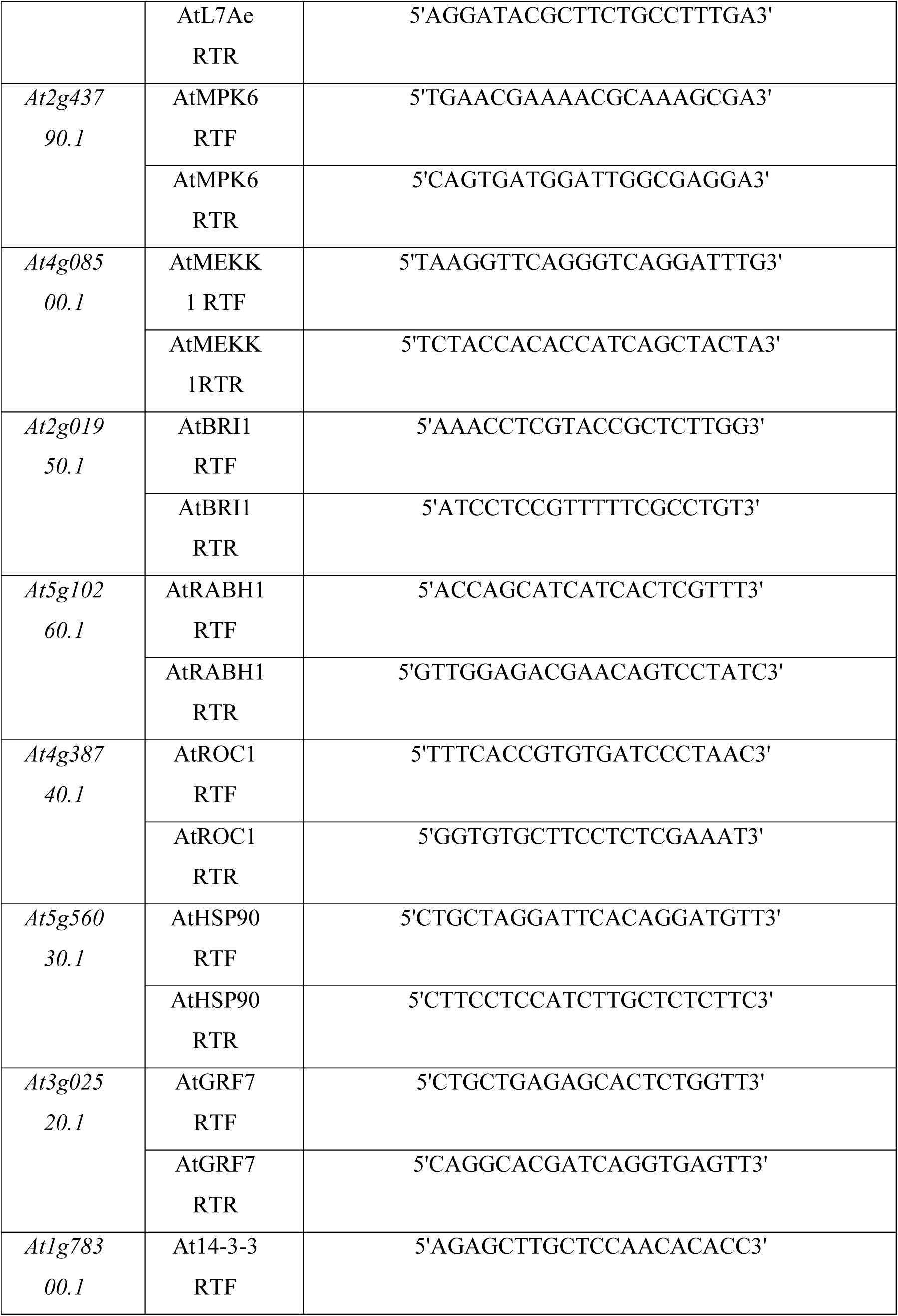
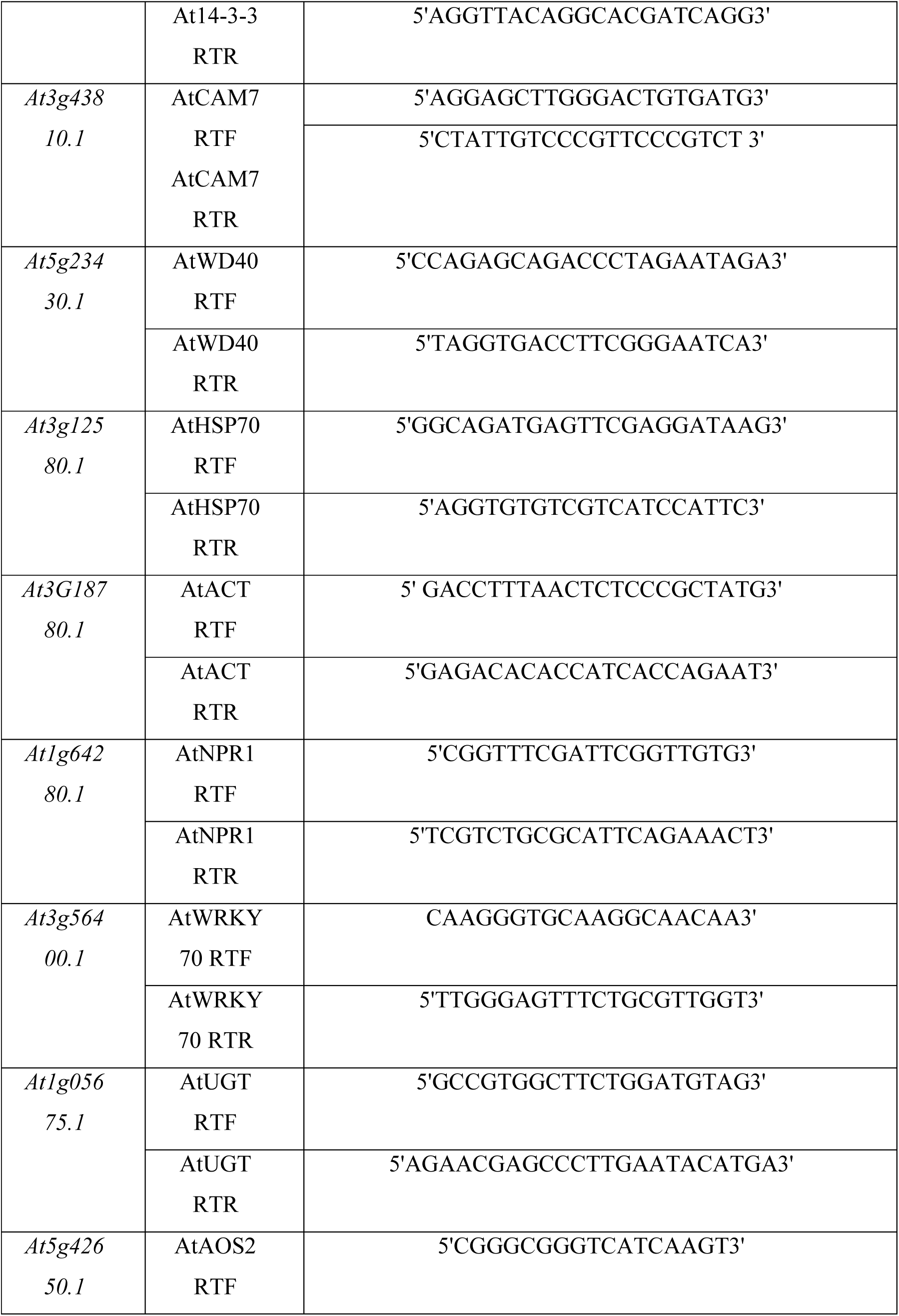
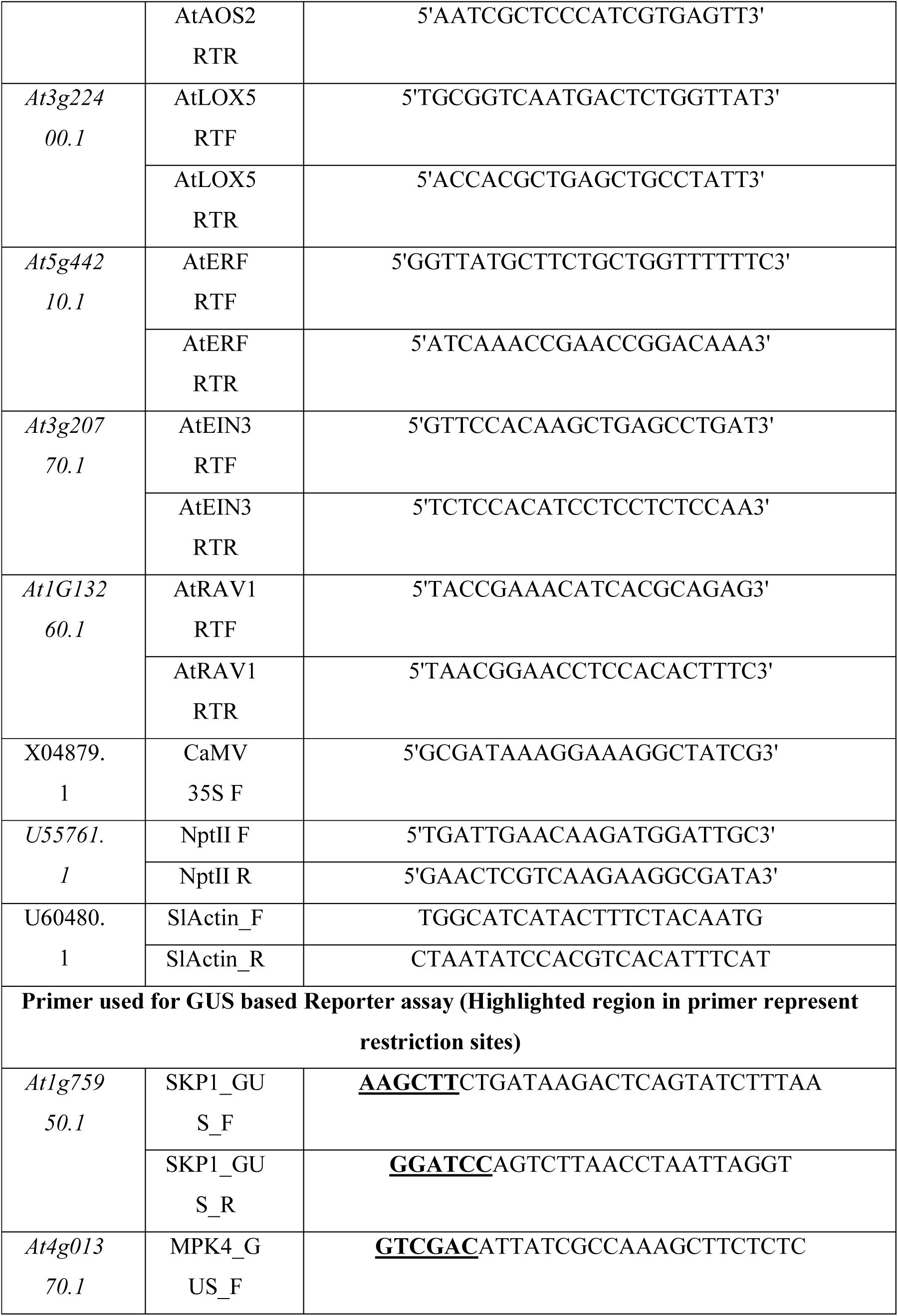
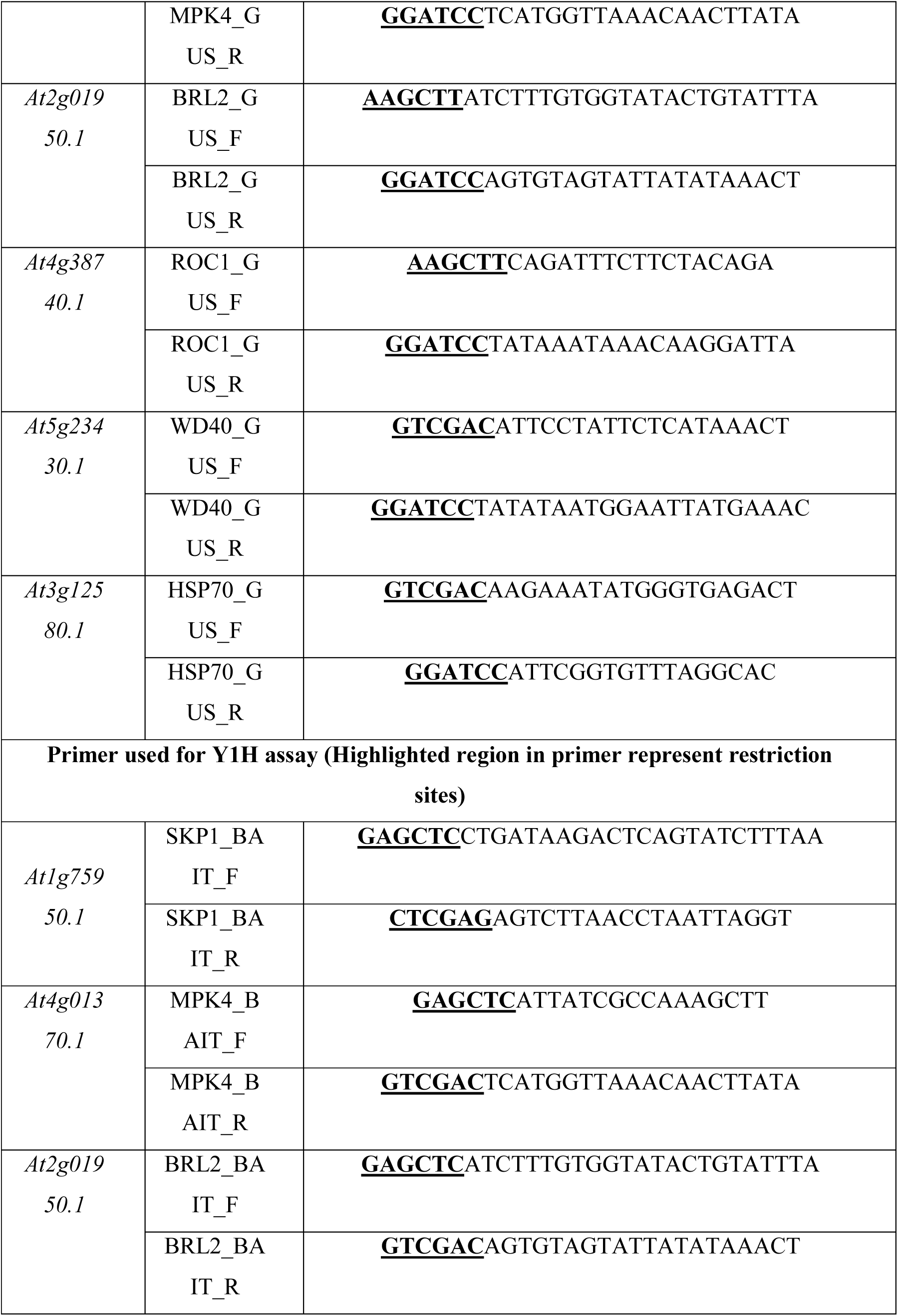
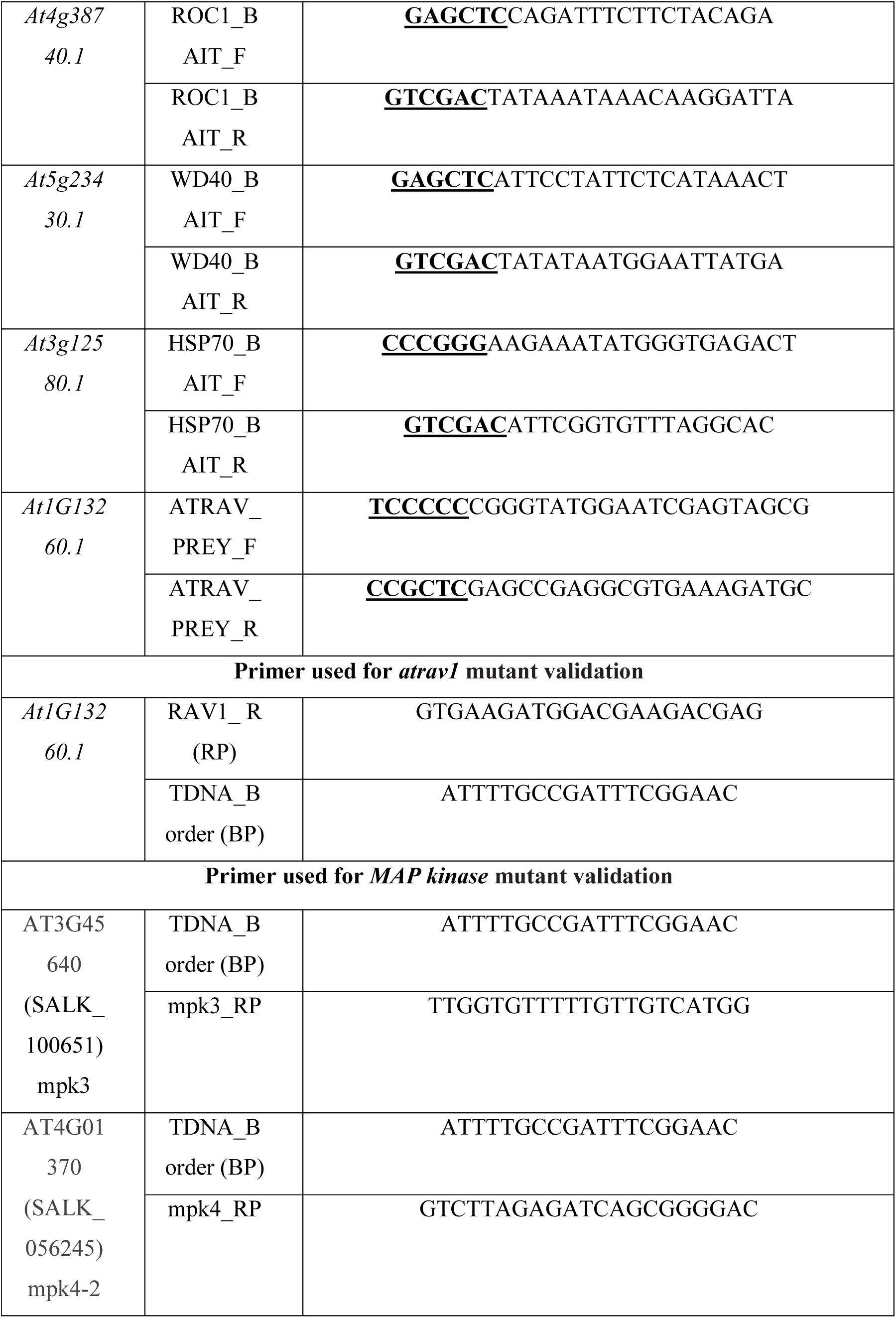
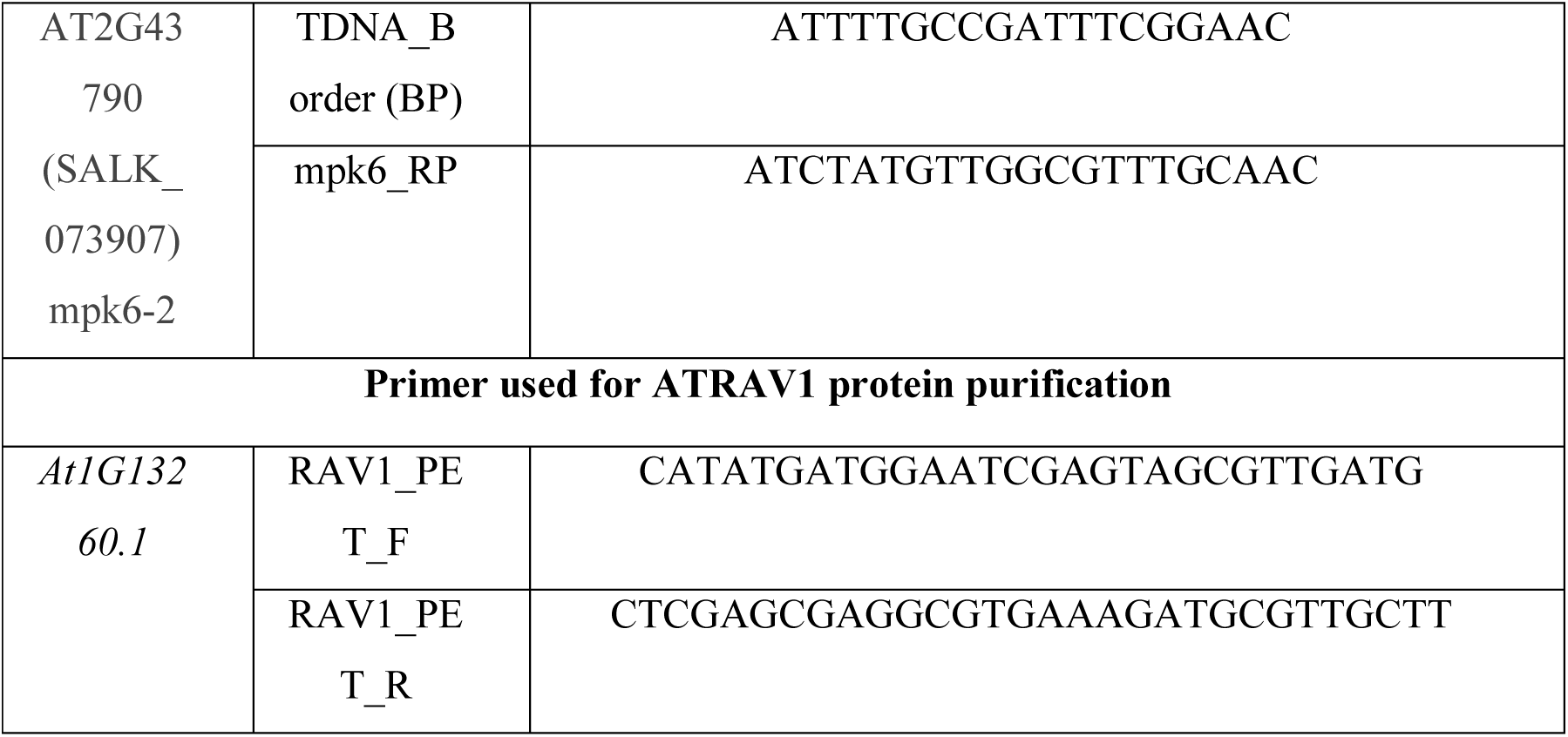
List of primer used in this study.

## References

1. J. D. G. Jones, J. L. Dangl, The plant immune system. Nature 444, 323–9 (2006).

2. Z. Ma, et al., A Phytophthora sojae Glycoside Hydrolase 12 Protein Is a Major Virulence Factor during Soybean Infection and Is Recognized as a PAMP. Plant Cell (2015) https://doi.org/10.1105/tpc.15.00390.

3. C. Zipfel, et al., Perception of the Bacterial PAMP EF-Tu by the Receptor EFR Restricts Agrobacterium-Mediated Transformation. Cell (2006) https://doi.org/10.1016/j.cell.2006.03.037.

4. S. Stael, et al., Plant innate immunity - sunny side up? Trends Plant Sci. (2015) https://doi.org/10.1016/j.tplants.2014.10.002.

5. S. H. Spoel, X. Dong, How do plants achieve immunity? Defence without specialized immune cells. Nat. Rev. Immunol. 12, 89–100 (2012).

6. S. T. Chisholm, G. Coaker, B. Day, B. J. Staskawicz, Host-microbe interactions: shaping the evolution of the plant immune response. Cell 124, 803–814 (2006).

7. L. Bacete, H. Mélida, E. Miedes, A. Molina, Plant cell wall-mediated immunity: cell wall changes trigger disease resistance responses. Plant J. (2018) https://doi.org/10.1111/tpj.13807.

8. L. Wang, et al., Integrated transcriptome and hormone profiling highlight the role of multiple phytohormone pathways in wheat resistance against fusarium head blight. PLoS One (2018) https://doi.org/10.1371/journal.pone.0207036.

9. V. Verma, P. Ravindran, P. P. Kumar, Plant hormone-mediated regulation of stress responses. BMC Plant Biol. (2016) https://doi.org/10.1186/s12870-016-0771-y.

10. L. Caarls, et al., Assessing the role of ETHYLENE RESPONSE FACTOR transcriptional repressors in salicylic acid-mediated suppression of jasmonic acid-responsive genes. Plant Cell Physiol. (2017) https://doi.org/10.1093/pcp/pcw187.

11. W. F. Broekaert, S. L. Delauré, M. F. C. De Bolle, B. P. A. Cammue, The role of ethylene in host-pathogen interactions. Annu. Rev. Phytopathol. 44, 393–416 (2006).

12. M. T. Nishimura, J. L. Dangl, Arabidopsis and the plant immune system. Plant J. (2010) https://doi.org/10.1111/j.1365-313X.2010.04131.x.

13. M. L. Berens, H. M. Berry, A. Mine, C. T. Argueso, K. Tsuda, Evolution of Hormone Signaling Networks in Plant Defense. Annu. Rev. Phytopathol. 55, annurev-phyto-080516-035544 (2017).

14. A. Robert-Seilaniantz, M. Grant, J. D. G. Jones, Hormone Crosstalk in Plant Disease and Defense: More Than Just JASMONATE-SALICYLATE Antagonism. Annu. Rev. Phytopathol. 49, 317–343 (2011).

15. N. Peeters, A. Guidot, F. Vailleau, M. Valls, Ralstonia solanacearum, a widespread bacterial plant pathogen in the post-genomic era. Mol. Plant Pathol. (2013) https://doi.org/10.1111/mpp.12038.

16. S. Genin, Molecular traits controlling host range and adaptation to plants in Ralstonia solanacearum. New Phytol. (2010) https://doi.org/10.1111/j.1469-8137.2010.03397.x.

17. A. C. Hayward, Biology and Epidemiology of Bacterial Wilt Caused by Pseudomonas Solanacearum. Annu. Rev. Phytopathol. 29, 65–87 (1991).

18. S. Ghosh, S. K. Gupta, G. Jha, Identification and functional analysis of AG1-IA specific genes of Rhizoctonia solani. Curr. Genet. 60, 327–341 (2014).

19. G. Yang, C. Li, “General Description of Rhizoctonia Species Complex” in Plant Pathology, (2012) https://doi.org/10.5772/39026.

20. E. Moriones, S. Praveen, S. Chakraborty, Tomato leaf curl new delhi virus: An emerging virus complex threatening vegetable and fiber crops. Viruses (2017) https://doi.org/10.3390/v9100264.

21. C. Albrecht, et al., Brassinosteroids inhibit pathogen-associated molecular pattern-triggered immune signaling independent of the receptor kinase BAK1. Proc. Natl. Acad. Sci. 109, 303–308 (2012).

22. H. He, et al., Pm21, Encoding a Typical CC-NBS-LRR Protein, Confers Broad-Spectrum Resistance to Wheat Powdery Mildew Disease. Mol. Plant (2018) https://doi.org/10.1016/j.molp.2018.03.004.

23. R. Backer, S. Naidoo, N. van den Berg, The NONEXPRESSOR OF PATHOGENESIS-RELATED GENES 1 (NPR1) and Related Family: Mechanistic Insights in Plant Disease Resistance. Front. Plant Sci. 10, 102 (2019).

24. K. M. Pajerowska-Mukhtar, D. K. Emerine, M. S. Mukhtar, Tell me more: Roles of NPRs in plant immunity. Trends Plant Sci. (2013) https://doi.org/10.1016/j.tplants.2013.04.004.

25. S. Lacombe, et al., Interfamily transfer of a plant pattern-recognition receptor confers broad-spectrum bacterial resistance. Nat. Biotechnol. 28, 365–369 (2010).

26. J. E. Lincoln, et al., Expression of the antiapoptotic baculovirus p35 gene in tomato blocks programmed cell death and provides broad-spectrum resistance to disease. Proc. Natl. Acad. Sci. (2002) https://doi.org/10.1073/pnas.232579799.

27. R. Nelson, T. Wiesner-Hanks, R. Wisser, P. Balint-Kurti, Navigating complexity to breed disease-resistant crops. Nat. Rev. Genet. (2018) https://doi.org/10.1038/nrg.2017.82.

28. Y. Kagaya, K. Ohmiya, T. Hattori, RAV1, a novel DNA-binding protein, binds to bipartite recognition sequence through two distinct DNA-binding domains uniquely found in higher plants. Nucleic Acids Res. 27, 470–478 (1999).

29. C. Z. Feng, et al., Arabidopsis RAV1 transcription factor, phosphorylated by SnRK2 kinases, regulates the expression of ABI3, ABI4, and ABI5 during seed germination and early seedling development. Plant J. 80, 654–668 (2014).

30. H. R. Woo, et al., The RAV1 transcription factor positively regulates leaf senescence in Arabidopsis. J. Exp. Bot. (2010) https://doi.org/10.1093/jxb/erq206.

31. M. S. Mukhtar, et al., Plant Immune System Network. Science (80-.). 333, 596–601 (2011).

32. Y. T. Cheng, et al., Stability of plant immune-receptor resistance proteins is controlled by SKP1-CULLIN1-F-box (SCF)-mediated protein degradation. Proc. Natl. Acad. Sci. 108, 14694–14699 (2011).

33. Y. Liu, et al., Phosphorylation of an ERF Transcription Factor by Arabidopsis MPK3/MPK6 Regulates Plant Defense Gene Induction and Fungal Resistance. Plant Cell (2013) https://doi.org/10.1105/tpc.112.109074.

34. M. Zhang, J. Su, Y. Zhang, J. Xu, S. Zhang, Conveying endogenous and exogenous signals: MAPK cascades in plant growth and defense. Curr. Opin. Plant Biol. 45, 1–10 (2018).

35. J. Jelenska, J. A. van Hal, J. T. Greenberg, Pseudomonas syringae hijacks plant stress chaperone machinery for virulence. Proc. Natl. Acad. Sci. (2010) https://doi.org/10.1073/pnas.0910943107.

36. C. J. Park, Y. S. Seo, Heat shock proteins: A review of the molecular chaperones for plant immunity. Plant Pathol. J. (2015) https://doi.org/10.5423/PPJ.RW.08.2015.0150.

37. G. Kong, et al., The Activation of Phytophthora Effector Avr3b by Plant Cyclophilin is Required for the Nudix Hydrolase Activity of Avr3b. PLoS Pathog. 11, 1–22 (2015).

38. J. R. Dominguez-Solis, et al., A cyclophilin links redox and light signals to cysteine biosynthesis and stress responses in chloroplasts. Proc. Natl. Acad. Sci. U. S. A. 105, 16386–91 (2008).

39. G. J. M. Beckers, et al., Mitogen-Activated protein kinases 3 and 6 are required for full priming of stress responses in Arabidopsis thaliana. Plant Cell (2009) https://doi.org/10.1105/tpc.108.062158.

40. K. H. Sohn, S. C. Lee, H. W. Jung, J. K. Hong, B. K. Hwang, Expression and functional roles of the pepper pathogen-induced transcription factor RAV1 in bacterial disease resistance, and drought and salt stress tolerance. Plant Mol. Biol. 61, 897–915 (2006).

41. A. Djamei, A. Pitzschke, H. Nakagami, I. Rajh, H. Hirt, Trojan horse strategy in Agrobacterium transformation: Abusing MAPK defense signaling. Science (80-.). (2007) https://doi.org/10.1126/science.1148110.

42. B. Li, X. Meng, L. Shan, P. He, Transcriptional Regulation of Pattern-Triggered Immunity in Plants. Cell Host Microbe (2016) https://doi.org/10.1016/j.chom.2016.04.011.

43. G. Mao, et al., Phosphorylation of a WRKY Transcription Factor by Two Pathogen-Responsive MAPKs Drives Phytoalexin Biosynthesis in *Arabidopsis*. Plant Cell 23, 1639–1653 (2011).

44. J. Su, et al., Active photosynthetic inhibition mediated by MPK3/MPK6 is critical to effector-triggered immunity. PLoS Biol. (2018) https://doi.org/10.1371/journal.pbio.2004122.

45. S. C. Popescu, et al., MAPK target networks in Arabidopsis thaliana revealed using functional protein microarrays. Genes Dev. (2009) https://doi.org/10.1101/gad.1740009.

46. X. Meng, et al., Phosphorylation of an ERF Transcription Factor by Arabidopsis MPK3/MPK6 Regulates Plant Defense Gene Induction and Fungal Resistance. Plant Cell 25, 1126–1142 (2013).

47. P. Singh, A. K. Sinha, A Positive Feedback Loop Governed by SUB1A1 Interaction with MITOGEN-ACTIVATED PROTEIN KINASE3 Imparts Submergence Tolerance in Rice. Plant Cell (2016) https://doi.org/10.1105/tpc.15.01001.

48. B. Huot, J. Yao, B. L. Montgomery, S. Y. He, Growth-defense tradeoffs in plants: A balancing act to optimize fitness. Mol. Plant 7, 1267–1287 (2014).

49. K. J. P. Silva, A. Brunings, N. A. Peres, Z. Mou, K. M. Folta, The Arabidopsis NPR1 gene confers broad-spectrum disease resistance in strawberry. Transgenic Res. 24, 693–704 (2015).

50. R. A. Gutierrez, R. M. Ewing, J. M. Cherry, P. J. Green, Identification of unstable transcripts in Arabidopsis by cDNA microarray analysis: Rapid decay is associated with a group of touch- and specific clock-controlled genes. Proc. Natl. Acad. Sci. 99, 11513–11518 (2002).

51. M. S. Mukhtar, et al., Independently evolved virulence effectors converge onto hubs in a plant immune system network. Science 333, 596–601 (2011).

52. M. M. Brandão, L. L. Dantas, M. C. Silva-Filho, AtPIN: Arabidopsis thaliana protein interaction network. BMC Bioinformatics 10, 454 (2009).

53. P. Shannon, A. Markiel, O. Ozier, N. Baliga, Cytoscape: a software environment for integrated models of biomolecular interaction networks. Genome, 2498–2504 (2003).

54. S. J. Clough, A. F. Bent, Floral dip: A simplified method for Agrobacterium-mediated transformation of Arabidopsis thaliana. Plant J. 16, 735–743 (1998).

55. Y. Chen, GUS staining of guard cells to identify localised guard cell gene expression. BIO-PROTOCOL (2017) https://doi.org/10.21769/bioprotoc.2446.

56. K. J. Livak, T. D. Schmittgen, Analysis of Relative Gene Expression Data Using Real-Time Quantitative PCR and the 2 Ϫ⌬⌬C T Method. METHODS 25, 402–408 (2001).

57. D. M. Swain, et al., A prophage tail-like protein is deployed by Burkholderia bacteria to feed on fungi. Nat. Commun. (2017) https://doi.org/10.1038/s41467-017-00529-0.

58. D. I. Arnon, Copper Enzymes in Isolated Chloroplasts. Polyphenoloxidase in Beta Vulgaris. Plant Physiol. 24, 1–15 (1949).

59. D. Khokhani, T. M. Lowe-Power, T. M. Tran, C. Allen, A single regulator mediates strategic switching between attachment/spread and growth/virulence in the plant pathogen Ralstonia solanacearum. MBio (2017) https://doi.org/10.1128/mBio.00895-17.

60. R. K. Chandan, et al., Silencing of tomato CTR1 provides enhanced tolerance against Tomato leaf curl virus infection. Plant Signal. Behav. (2019) https://doi.org/10.1080/15592324.2019.1565595.

61. D. M. Swain, et al., Concurrent overexpression of rice G-protein β and γ subunits provide enhanced tolerance to sheath blight disease and abiotic stress in rice. Planta (2019) https://doi.org/10.1007/s00425-019-03241-z.

62. R. Kant, K. Tyagi, S. Ghosh, G. Jha, Host Alternative NADH:Ubiquinone Oxidoreductase Serves as a Susceptibility Factor to Promote Pathogenesis of Rhizoctonia solani in Plants. Phytopathology (2019) https://doi.org/10.1094/phyto-02-19-0055-r.

63. B. Raghuram, A. H. Sheikh, Y. Rustagi, A. K. Sinha, MicroRNA biogenesis factor DRB1 is a phosphorylation target of mitogen activated protein kinase MPK3 in both rice and Arabidopsis. FEBS J. (2015) https://doi.org/10.1111/febs.13159.

## References

1. P. Nie, et al., Induced Systemic Resistance against Botrytis cinerea by Bacillus cereus AR156 through a JA/ET- and NPR1-Dependent Signaling Pathway and Activates PAMP-Triggered Immunity in Arabidopsis. Front. Plant Sci. 8, 1–12 (2017).

2. W. Fan, X. Dong, In Vivo Interaction between NPR1 and Transcription Factor TGA2 Leads to Salicylic Acid–Mediated Gene Activation in Arabidopsis. Plant Cell 14, 1377–1389 (2002).

3. J. Li, G. Brader, T. Kariola, E. Tapio Palva, WRKY70 modulates the selection of signaling pathways in plant defense. Plant J. 46, 477–491 (2006).

4. Z. Liu, et al., UDP-glucosyltransferase71C5, a major glucosyltransferase, mediates abscisic acid homeostasis in Arabidopsis. Plant Physiol. 167, 1659–70 (2015).

5. K. M. Pajerowska, J. E. Parker, C. Gebhardt, Potato homologs of Arabidopsis thaliana genes functional in defense signaling - Identification, genetic mapping, and molecular cloning. Mol. Plant-Microbe Interact. 18, 1107–1119 (2005).

6. R. Marcos, et al., 9-Lipoxygenase-derived oxylipins activate brassinosteroid signaling to promote cell wall-based defense and limit pathogen infection. Plant Physiol. 4, pp.00992.2015 (2015).

7. Y. Maruyama, et al., The Arabidopsis transcriptional repressor ERF9 participates in resistance against necrotrophic fungi. Plant Sci. 213, 79–87 (2013).

8. R. Quan, et al., EIN3 and SOS2 synergistically modulate plant salt tolerance. Sci Rep 7, 44637 (2017).

9. A. Wawrzyñska, A. Sirko, EIN3 interferes with the sulfur deficiency signaling in Arabidopsis thaliana through direct interaction with the SLIM1 transcription factor. Plant Sci. 253, 50–57 (2016).

10. F. Liu, et al., The ASK1 and ASK2 genes are essential for Arabidopsis early development. Plant Cell 16, 5–20 (2004).

11. A. Takahashi, C. Casais, K. Ichimura, K. Shirasu, HSP90 interacts with RAR1 and SGT1 and is essential for RPS2-mediated disease resistance in Arabidopsis. Proc. Natl. Acad. Sci. U. S. A. 100, 11777–11782 (2003).

12. E. Mazzucotelli, et al., The e3 ubiquitin ligase gene family in plants: regulation by degradation. Curr. Genomics 7, 509–22 (2006).

13. Y. Wang, M. Yang, The ARABIDOPSIS SKP1-LIKE1 (ASK1) protein acts predominately from leptotene to pachytene and represses homologous recombination in male meiosis. Planta 223, 613–617 (2006).

14. M. Petersen, et al., Arabidopsis map kinase 4 negatively regulates systemic acquired resistance. Cell 103, 1111–1120 (2000).

15. Q. Zeng, J. G. Chen, B. E. Ellis, AtMPK4 is required for male-specific meiotic cytokinesis in Arabidopsis. Plant J. 67, 895–906 (2011).

16. A. J. Carroll, J. L. Heazlewood, J. Ito, A. H. Millar, Analysis of the Arabidopsis cytosolic ribosome proteome provides detailed insights into its components and their post-translational modification. Mol. Cell. Proteomics 7, 347–369 (2008).

17. J. S. López-Bucio, et al., Arabidopsis thaliana mitogen-activated protein kinase 6 is involved in seed formation and modulation of primary and lateral root development. J. Exp. Bot. 65, 169–183 (2014).

18. M. A. T. Palm-Forster, L. Eschen-Lippold, J. Uhrig, D. Scheel, J. Lee, A novel family of proline/serine-rich proteins, which are phospho-targets of stress-related mitogen-activated protein kinases, differentially regulates growth and pathogen defense in Arabidopsis thaliana. Plant Mol. Biol. (2017) https://doi.org/10.1007/s11103-017-0641-5.

19. T. Asai, et al., Map kinase signalling cascade in Arabidopsis innate immunity. Nature 415, 977–983 (2002).

20. T. Furuya, D. Matsuoka, T. Nanmori, Membrane rigidification functions upstream of the MEKK1-MKK2-MPK4 cascade during cold acclimation in Arabidopsis thaliana. FEBS Lett. 588, 2025–2030 (2014).

21. J. L. Nemhauser, T. C. Mockler, J. Chory, Interdependency of brassinosteroid and auxin signaling in Arabidopsis. PLoS Biol. 2 (2004).

22. B. Kemmerling, et al., The BRI1-Associated Kinase 1, BAK1, Has a Brassinolide-Independent Role in Plant Cell-Death Control. Curr. Biol. 17, 1116–1122 (2007).

23. V. Vernoud, A. C. Horton, Z. Yang, E. Nielsen, Analysis of the small GTPase gene superfamily of Arabidopsis. Plant Physiol. 131, 1191–208 (2003).

24. N. Kato, H. He, A. P. Steger, A systems model of vesicle trafficking in Arabidopsis pollen tubes. Plant Physiol. 152, 590–601 (2010).

25. S. A. Trupkin, S. Mora-García, J. J. Casal, The cyclophilin ROC1 links phytochrome and cryptochrome to brassinosteroid sensitivity. Plant J. 71, 712–723 (2012).

26. M. Clement, et al., The cytosolic/nuclear HSC70 and HSP90 molecular chaperones are important for stomatal closure and modulate abscisic acid-dependent physiological responses in Arabidopsis. Plant Physiol. 156, 1481–1492 (2011).

27. S. Huang, et al., HSP90s are required for NLR immune receptor accumulation in Arabidopsis. Plant J. 79, 427–439 (2014).

28. J. Keicher, et al., Arabidopsis 14-3-3 epsilon members contribute to polarity of PIN auxin carrier and auxin transport-related development. Elife 6 (2017).

29. H. Fulgosi, et al., 14-3-3 proteins and plant development. Plant Mol. Biol. 50, 1019–1029 (2002).

30. S. S. Gampala, et al., An essential role for 14-3-3 proteins in brassinosteroid signal transduction in Arabidopsis. Dev. Cell 13, 177–189 (2007).

31. M.-H. Oh, et al., Calcium/calmodulin inhibition of the Arabidopsis BRASSINOSTEROID-INSENSITIVE 1 receptor kinase provides a possible link between calcium and brassinosteroid signalling. Biochem. J. 443, 515–523 (2012).

32. N. Abbas, J. P. Maurya, D. Senapati, S. N. Gangappa, S. Chattopadhyay, Arabidopsis CAM7 and HY5 physically interact and directly bind to the HY5 promoter to regulate its expression and thereby promote photomorphogenesis. Plant Cell 26, 1036–1052 (2014).

33. E. W. Gachomo, J. C. Jimenez-Lopez, L. J. Baptiste, S. O. Kotchoni, GIGANTUS1 (GTS1), a member of Transducin/WD40 protein superfamily, controls seed germination, growth and biomass accumulation through ribosome-biogenesis protein interactions in Arabidopsis thaliana. BMC Plant Biol. 14, 1–17 (2014).

34. L. Leng, et al., A subclass of HSP70s regulate development and abiotic stress responses in Arabidopsis thaliana. J. Plant Res. 130, 349–363 (2017).

35. P. Pulido, E. Llamas, M. Rodriguez-Concepcion, Both Hsp70 chaperone and Clp protease plastidial systems are required for protection against oxidative stress. Plant Signal. Behav. 12, e1290039 (2017).

